# Computationally guided exploration of borosin biosynthetic strategies

**DOI:** 10.1101/2024.01.09.574750

**Authors:** Aileen R. Lee, Riley S. Carter, Aman S. Imani, Shravan R. Dommaraju, Graham A. Hudson, Douglas A. Mitchell, Michael F. Freeman

## Abstract

Borosins are ribosomally synthesized and post-translationally modified peptides containing backbone α-*N*-methylations. Identification of borosin precursor peptides is difficult because (1) there are no conserved sequence elements among borosin precursor peptides and (2) the biosynthetic gene clusters contain numerous domain architectures and peptide fusions. To tackle this problem, we updated the genome mining tool RODEO to automatically evaluate putative borosin BGCs and identify precursor peptides. Enabled by the new borosin module, we analyzed all borosin BGCs found in available sequence data and assigned precursor peptides to previously orphan borosin methyltransferases. Additionally, we bioinformatically predict and experimentally characterize a new fused borosin domain architecture, in which the modified core is N-terminal to the methyltransferase domain. Finally, we demonstrate that a borosin precursor peptide is the native substrate of shewasin A, a previously characterized pepsin-like aspartic peptidase whose native biological function was unknown.

## Introduction

Borosins are a class of ribosomally synthesized and post-translationally modified peptides (RiPPs) characterized by the presence of amide backbone α-*N*-methylations.^1^ In RiPP biosynthesis, the N-terminus of the ribosomally produced precursor peptide typically contains a leader region, which is recognized by enzymes that modify the C-terminal core region into the mature RiPP. These modifying enzymes are often encoded near the precursor peptide, together forming a biosynthetic gene cluster (BGC). The maturation process includes cleavage of the core from the leader region by a protease that may or may not be encoded in the RiPP BGC.^2^Click or tap here to enter text. The first discovered borosin, omphalotin A, was isolated from the basidiomycete fungus *Omphalotus olearius*.^1,3^ The omphalotin BGC contained a novel RiPP domain architecture in which the core peptide is fused to the enzymatic α-*N*-methyltransferase (NMT) domain.^1^ Since the initial characterization of the omphalotin BGC, bioinformatic and biochemical studies have discovered additional borosin BGCs in fungi, archaea, and bacteria^4–6^ with a diverse set of BGC and NMT domain architectures. Among the domain architectures are three distinct types of “fused borosins,” such as the type found in *O. olearius*, and seven types of “split borosins” that encode the precursor peptide and NMT as separate genes. To date, >20 borosin NMTs have been biochemically characterized.^1,4–7^ However, other steps of borosin maturation remain understudied, including proteolytic cleavage of the precursor peptide into the mature RiPP. The only characterized protease involved in borosin biosynthesis is OphP, the macrocyclase from the omphalotin BGC.^8,9^

Computationally identifying borosins is challenging for numerous reasons. Borosin NMTs are homologs of the large tetrapyrrole (TP) methylase family (PF00590, >175,000 members) and can be difficult to distinguish from other TP methylases in the family. Furthermore, identification of borosin precursor peptides has several nontrivial obstacles. Whereas RiPP precursor peptides are typically ∼100 amino acids (AAs) or fewer, borosin precursors can vary in length from ∼60 to ∼600 AAs, and are not always encoded as discrete precursor peptides.^5,6^ Additionally, the α*-N*-methylation of the peptide backbone appears indiscriminate, as borosin core sequences often lack defining characteristics. Borosins may be enriched in aliphatic, polar or acidic residues, with the methylation occurring on any of these residue types. Borosins also vary in the number of methylations that occur; some sequences contain just two methylation sites while others can contain >40, as in the case of SliA from *Spirosoma liguale* DSM 74.^5^ When identifying precursor peptides, previous studies have relied on manual inspection of open reading frames (ORFs)^4,5^ or constructing sequence similarity networks (SSNs) of short ORFs found near the putative borosin NMTs.^6^ These methods are time-intensive and not preferable for large-scale bioinformatic analysis, thus, we sought to develop a streamlined approach to analyzing and assessing potential borosin BGCs.

RODEO (Rapid ORF Description & Evaluation Online) is a computational tool used to analyze genes and short ORFs neighboring an input protein query.^10^Several specialized modules which evaluate and identify putative precursor peptides for different classes of RiPPs have already been developed for RODEO.^10–15^In this work, we update the RODEO algorithm with the capability to automatically identify putative borosin precursor peptides using heuristic scoring, hidden Markov models (HMMs), motif analysis, and support vector machine classification. Using the new borosin module, we run a current analysis of borosin BGCs found in available sequence data and assign precursor peptides to previously orphan borosin NMTs. Additionally, we identify and biochemically verify a new type of fused borosin wherein the core sequence is found N-terminal to the NMT domain, which we designate as type 0 borosins. Finally, we demonstrate that a borosin precursor is the native substrate for shewasin A, a previously characterized pepsin-like aspartic peptidase from *Shewanella amazonensis*. Shewasin A requires an α*-N*-methylated leader peptide to release a core peptide devoid of α*-N*-methylations, a paradigm not previously observed in RiPP biosynthesis.

## Results & Discussion

### Development of a borosin RODEO scoring module

We first sought a method to robustly differentiate borosin NMTs from non-borosin methylases within the tetrapyrrole (TP) methylase family. Previous work relied on a combination of custom HMMs to identify putative borosin NMTs, but many of the sequences retrieved using the custom HMMs individually were found to be false positives when inspected manually.^5^ BLAST searches were performed against the NCBI non-redundant (nr) database using each characterized borosin NMT as a query (Table S1). Retrieved sequences were analyzed using hmmscan^16^ and a subset of the results was plotted based on the expectation value (e-value) for the TP methylases (PF00590) and BorosinMT HMMs (Fig. S1). We inspected and assigned borosin and non-borosin methyltransferase sequences (SI Methods). The graph comparing BorosinMT and PF00590 e-values showed two distinct trends that clearly separate borosin and non-borosin sequences, except at very poor e-values.

Next, we developed a set of heuristics to identify borosin precursor peptides based on accumulated knowledge of borosin BGCs (Table S2). Most identified borosin BGCs contain a conserved five-helical bundle, termed the Borosin Binding Domain (BBD), that is crucial for precursor peptide recognition by the NMT.^7^ Fused borosins (types I-III) contain a BBD and core sequence fused to an NMT domain. Some split borosin BGCs contain a BBD-NMT fusion (types V-VIII), while others contain a BBD-precursor peptide fusion (types IV, IX, and X) (Fig. 1). Several groups of split borosins also contain other conserved sequences within the BGC. These include several distinct sequence motifs found in a subset of borosin BGCs from Legionellales, which lack a BBD^6^ and a previously unreported conserved region often found in split borosin BGCs with a BBD-methyltransferase fusion (types VI, VII, and VIII). We made custom HMMs using these sequences that were used to aid precursor peptide identification (Dataset 1). The presence of these sequences, and in some cases the e-value similarity to HMMs, were included in the heuristic scoring. Additionally, borosin core peptides often contain repeating motifs or are enriched in certain amino acids at their C-terminus. Repeating motifs from characterized borosins were identified using the MEME suite^17^ and were incorporated into the heuristic scoring alongside the enrichment of certain amino acids in the precursor peptide.

**Figure 1:**
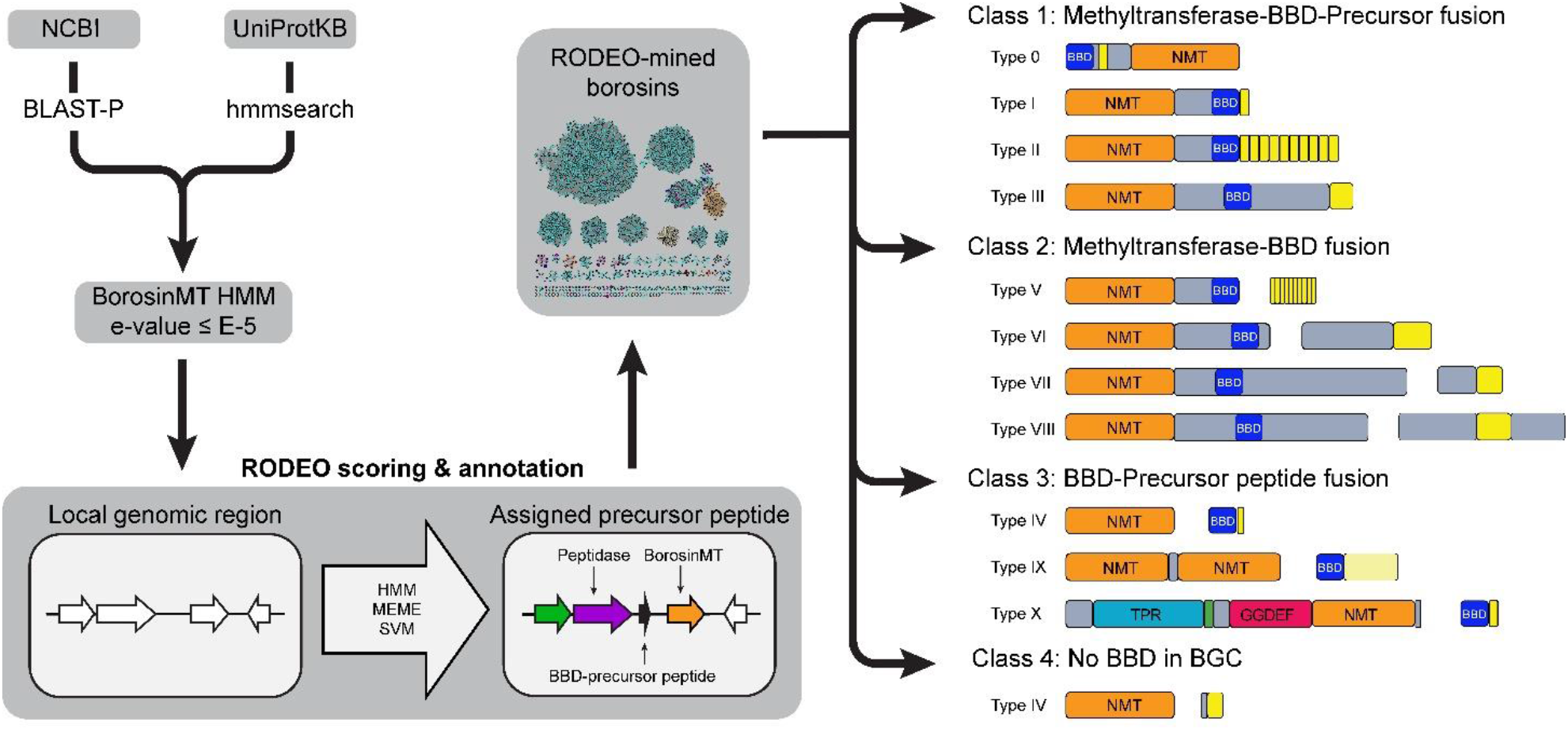
Workflow for bioinformatically mining borosins. Workflow used to create a bioinformatically mined borosin dataset. Borosin methyltransferase homologs are identified by BLAST-P and filtered by their e-value to the BorosinMT HMM to reduce computation time. Custom HMMs matches in UniProtKB are identified using hmmsearch. After removing duplicates, these lists of accessions were input into RODEO with the borosin scoring module. The RODEO-predicted precursors were then analyzed and assigned to different borosin types.

After defining the heuristic-, motif-, and HMM-based scoring metrics, we tested the borosin module on a set of high-confidence predicted borosins, but found it was insufficient to accurately identify borosin precursor peptides (Fig. S2, Fig. S3, SI methods). Support vector machine (SVM) classification, a supervised machine learning method, was trained to classify ORFs as borosin or non-borosin sequences using features from borosin and non-borosin BGCs (Table S3). Features used in this include gene co-occurrence, amino acid properties, and many of the heuristic scoring features. The SVM classification significantly increased module accuracy (Fig. S2, Fig. S3), and together with the heuristics, seemingly eliminated the issue of false positive BorosinMT matches. To decrease computation time, methyltransferases with an e-value higher than E-5 to the BorosinMT HMM were filtered out before analysis with the RODEO module (Fig. 1).

### Bioinformatics analysis of the borosin dataset

A dataset of borosin precursor peptides was curated using the new RODEO module (SI Methods). This dataset contains 2699 predicted borosin NMTs with 3064 predicted precursor peptides across 2596 BGCs (Dataset 2). The new RODEO module accurately predicted precursor peptides for 1565 previously identified putative borosin NMTs and identified precursor peptides for 1127 newly reported borosin BGCs. Previous bioinformatic studies on borosins identified an additional 733 NMTs that did not have a precursor identified by RODEO. After manual inspection, many of these BGCs still lacked an identifiable precursor peptide. These BGCs may represent new borosin architectures, those with distally encoded precursor peptides, or homologous non-borosin TP methylases. We assigned predicted borosin precursor peptides to their respective borosin types using similarity to HMMs, gene co-occurrence, and sequence information.^5^ An SSN was generated from NMTs in the RODEO-mined borosin dataset (Fig. 2, Fig. S4). Many of the largest SSN clusters contain biochemically characterized representatives, with clusters 7 and 8 as exceptions. Cluster 3 of the SSN is comprised of NMT-BBD fusions from borosin types I-III and VI-VIII. Cluster 3 is also more taxonomically diverse than clusters containing an NMT without a BBD, including members from both bacteria and fungi. Several unusual borosin BGC architectures were also identified, including examples containing a BBD-GNAT acetyltransferase fusion and a BGC with 11 BBD-fused precursor peptides (Fig. S5).

**Figure 2:**
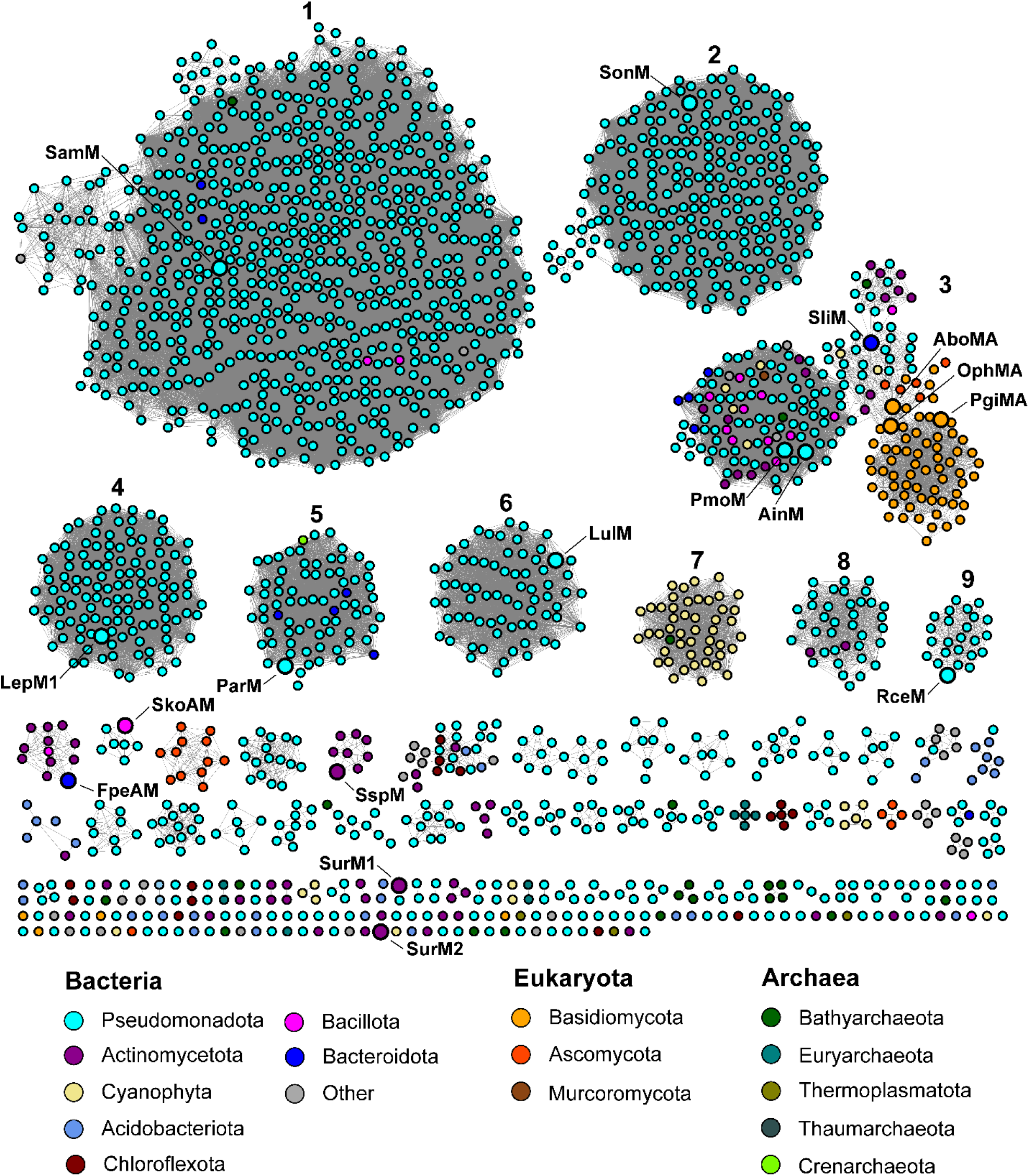
SSN of borosin methyltransferases. Borosin methyltransferases with a RODEO identified precursor peptide were used to generate an SSN (n=2,124). Sequences for OphMA, PgiMA and AboMA are not in the NCBI protein database and were added separately. An alignment score cutoff of 90 was used to clearly separate clusters.

We next investigated the evolution of borosin NMTs and BBDs. A phylogenetic tree was created using 469 diversity-maximized NMTs from the RODEO-mined borosin dataset, including members from each borosin type (SI methods, Fig. S6, Fig. S7). A custom script^14^ was employed to excise the region scoring highest for a BBD HMM from either the borosin NMT or split precursor in these BGCs (Fig. S8, Fig. S9). A phylogenetic tree was generated for these sequences, and the sum of the branch distances for the NMT and BBD trees were analyzed (Fig. S10). All characterized borosin NMTs are more similar to each other than to the PF00590 TP methylase outgroup, indicating these likely share a common ancestor. Excised BBDs also appear to have more sequence diversity and show similar relatedness between each other and the outgroup. This may result from conservation of secondary structure within the BBD rather than specific sequence motifs for precursor peptide recognition.

The borosin NMT from *Streptomyces* sp. NRRL S-118 (SspM; WP_158827804.1) and the associated BBD are the characterized split borosin pair most closely related to the fused borosins and outgroups. To get a more accurate root and branching order of borosin NMT evolution, SspM homologs in the NCBI nr database were identified using BLAST-P, and a larger phylogenetic tree was generated (Fig. S11, Fig. S12). Analysis of the resulting branch distances revealed several groups of RODEO-predicted type IV borosin NMTs that are positioned between the outgroups and characterized borosins (Fig. S13). As all characterized borosins are more similar to these predicted type IV borosins than the outgroups, borosins likely originated with a separate NMT and BBD-fused precursor peptide. The borosin NMTs from Legionellales lacking a BBD are closely related to other borosin NMTs, with distances comparable to other split borosin NMTs. This presumably indicates that the Legionellales BGCs originated from BBD-containing borosins that later lost the BBD.

While rare examples exist of fused borosins in bacteria, split borosins have not been identified in fungi. To look for evidence of fungi acquiring borosin BGCs through horizontal gene transfer (HGT), we compared borosin NMT gene G+C content to the content of surrounding contig genes. No evidence for HGT was observed for fungal borosins in this analysis. However, several promising examples were found in bacteria for HGT from lower G+C content organisms to organisms with a higher G+C content (Fig. S14). Additionally, clusters 2 and 4 of the borosin NMT SSN, which are mainly composed of *Shewanella* and *Legionella* species, respectively, contain many split borosin BGCs with nearby phage integrases. These mobile genetic elements may have enabled distribution of borosins within these genera (Fig. S15). Overall, borosins appear to have originated with a split BBD-fused precursor peptide, and diverged into two groups: a Pseudomonadota-dominated group with BBD-precursor peptides fusions and a taxonomically diverse group with NMT-BBD fusions.

### Discovery and characterization of type 0 borosins

Upon reviewing the excised BBDs from the analysis above, we discovered a previously uncharacterized group of fused borosins that we designated as type 0. Unlike other fused borosins (types I, II, and III), where the BBD and core peptide are encoded C-terminal to the NMT, the BBD and putative core of type 0 borosins are located at the N-terminus (Fig. 3). Thirty-two putative type 0 borosins were identified in the RODEO curated dataset based on the relative HMM alignment positions in BBD-fused NMTs (Fig. S4). Two of these type 0 borosins, FpeAM from *Flavihumibacter petaseus* NBRC 106054^18^ and SkoAM from *Segetibacter koreensis* DSM 18137,^19^ were selected for validation. To confirm NMT activity and core peptide sequences, we performed heterologous expression, purification, and analysis by high-resolution tandem mass spectrometry (LC-MS/MS). Up to 14 methylations in FpeAM and 10 in SkoAM were detected in the predicted core region between the BBD and methyltransferase domains (Fig. 3, Fig. S16).

**Figure 3:**
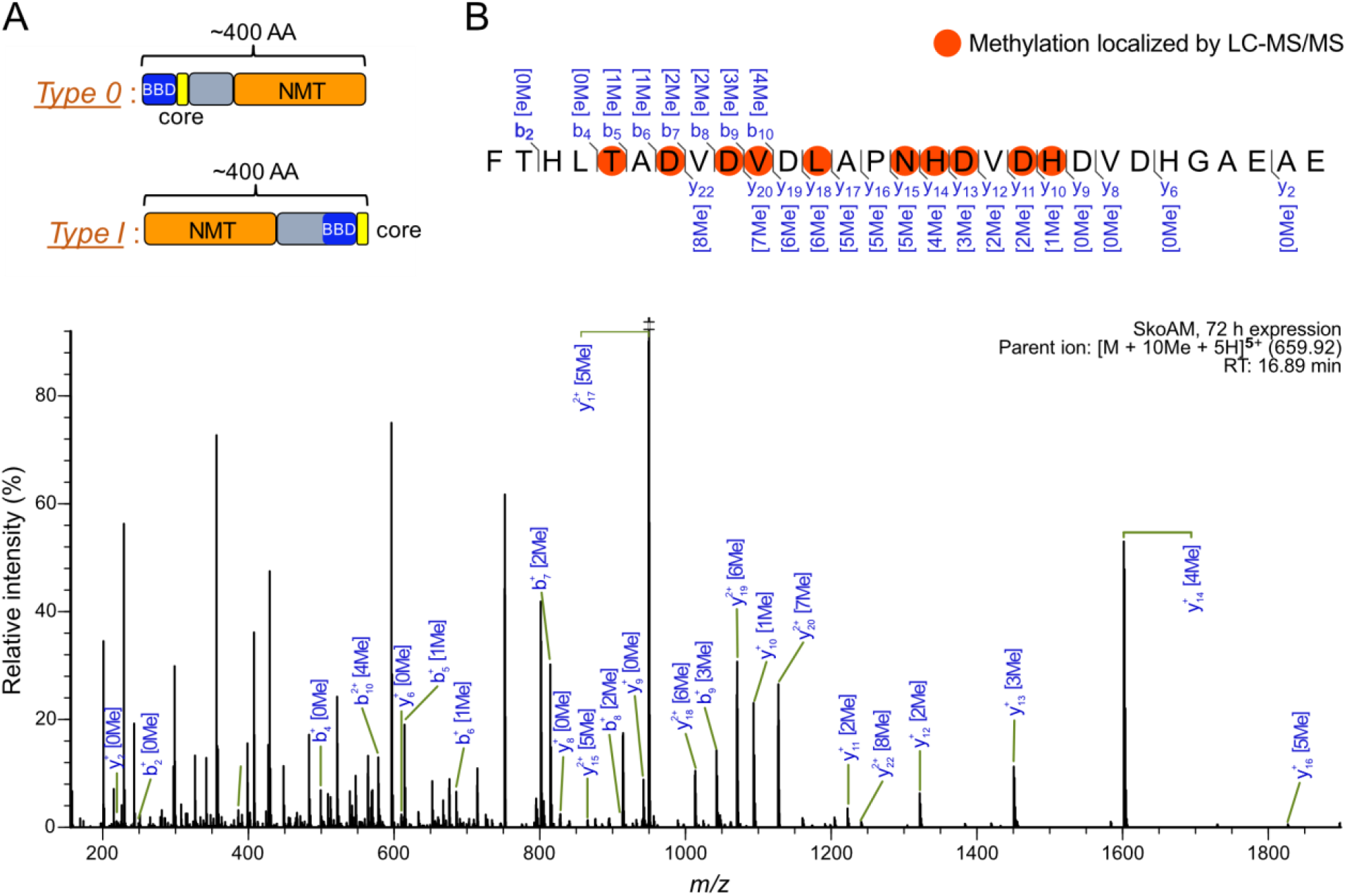
Type 0 borosin architecture and LC-MS/MS verification. **A**. Comparison of the overall domain architectures of the type 0 and type I fused borosins including the NMT (orange), BBD (blue), and core peptide (yellow) domains. **B**. LC-MS/MS data for example SkoAM heterologously produced in *E. coli* for 72 hrs. LC-MS/MS spectrum of SkoAM reveals methylated residues in a proteolytically released core peptide fragment. The borosin precursor peptide, time of expression, parent ion details, and LC retention times (RT) are denoted in the upper right-hand corner of the LC-MS/MS spectrum. Observed MS/MS fragmented masses are listed above (bions) and below (y-ions) the peptide sequence with grey lines marking sites of fragmentation. A mass cutoff of 10.0-ppm was used for the annotated LC-MS/MS peaks. Ion masses are denoted with varying numbers of methylations in brackets, where ‘Me’ marks a mass shift corresponding to methylation.

All currently published crystal structures of borosin NMTs show a dimeric conformation that facilitates an intermolecular mechanism in which one subunit modifies the core peptide of the other.^7,20,21^ In contrast, SkoAM was determined to be monomeric by size-exclusion chromatography (Fig. S17). This observation raises the possibility that type 0 borosins may catalyze intramolecular α-*N*-methylation, where each NMT modifies its own core peptide *in cis* rather than *in trans*. AlphaFold 2.0^22^ structural predictions of both monomeric and homodimeric SkoAM support the proposal of intramolecular catalysis based on the relative orientation of the BBD and NMT domains (Fig. S16). To confirm the size exclusion data was not an *in vitro* artifact, *trans*-complementation assays were performed similar to those used to first determine the intermolecular activity of OphMA.^1^ Briefly, an impaired mutant, SkoAM R214A, was expressed with an active SkoAM H121V core variant. Following expression, both proteins were co-purified, in-gel digested, and analyzed by LC-MS/MS (Fig. S18). Masses for the H121V core peptide variant containing 0-10 methylations were observed. In comparison, the wildtype core of the impaired R214A variant was predominantly unmodified, with a small amount of species containing 1-3 methylations (Fig. S19). Expression of the H121V mutant with wildtype SkoAM showed extensive methylation of both cores (Fig. S20). The robust methylation of the H121V variant coupled with the minimal modification of the impaired R214A variant supports an intramolecular mechanism for SkoAM.

### Shewasin A functions as a borosin RiPP peptidase

A significant advantage of the RODEO workflow is the ability to analyze ORFs neighboring the genes-of-interest. Since proteolytic cleavage is a pivotal step in the biosynthesis of many RiPPs^2^, we scanned for peptidases that commonly occur in borosin BGCs. Borosins from clusters 1 and 2 of the SSN either encode a PF00026 pepsin-like A1 aspartic peptidase in the BGC, or have one encoded distally in the organism (Fig. S20). Of the 2,699 Borosin NMTs in the RODEO-curated dataset, an aspartic peptidase is found within 194 BGCs. A PSI-BLAST of the NCBI nr database for homologs yielded 441 aspartic peptidases after removing fragmented sequencing reads. After submitting these proteins to the borosin RODEO module, an additional 73 putative borosin BGCs were identified. Of the remaining aspartic peptidases, 56 are found in organisms with a predicted borosin BGC, and 118 (including all 84 examples from Eukarya) were not associated with borosins. A phylogenetic tree of the peptidases suggests that a non-borosin peptidase gene was transferred to bacteria from a eukaryote before transitioning to become involved in borosin biosynthesis (Fig. S22). Inspection of the genomic neighborhood around the non-borosin peptidases did not reveal evidence of being involved in RiPP biosynthesis.

Curiously, A1 family peptidases are primarily found in eukaryotes, with few identified in bacteria.^23^ Two studies were conducted on the proteolytic activities of shewasin A and close homolog shewasin D, pepsin-like aspartic peptidases from *Shewanella amazonensis*^24,25^ and *Shewanella denitrificans*^25^, respectively. These studies revealed shewasins A and D had modest proteolytic activity under acidic conditions on a suite of model peptide substrates. Interestingly, our RODEO-mined borosin dataset revealed that both shewasin A and shewasin D are found in putative borosin BGCs, indicating that they may play a vital role in borosin RiPP maturation.

To determine if shewasin A (referred to hereafter as SamP) functions as a borosin RiPP peptidase, we first confirmed that the nearby borosin NMT (SamM1) is active on the predicted BBD-fused precursor peptide (SamA1). Genes *samM1* and *samA1* were heterologously expressed in *E. coli* and purified by nickel affinity chromatography. LC-MS/MS analysis of in-gel trypsinized SamA1 revealed a single α-*N-*methylation on Phe52 (Fig. S23). Expressions over 24 and 48 hours had >99% conversion to the methylated product compared to <10% conversion when expressed with a less active variant of SamM1 (Fig. S24-S26). These results confirm that SamM1 is an active borosin NMT, and SamA1 is the naturally encoded substrate.

Next, *samP* was heterologously expressed and purified, then assayed with methylated and unmethylated SamA1 using *in vitro* reactions monitored by SDS-PAGE (Fig. 4). Efficient cleavage of α-*N-*methylated SamA1 was observed, while only basal SamP activity was detected with unmethylated SamA1. Proteolytic activity was observed up to pH 6.0, while significant precipitation of SamP was seen at pH <6.0 (Fig. S27). Proteolytic activity was not observed using an inactive SamP D37A variant^24^ (Fig. S27). These results indicate that the RiPP precursor peptide, SamA1, is likely the native SamP substrate, and activity is α-*N*-methylation and pH dependent. Previous studies reported modest proteolytic activity of SamP only at pH 2.5-5.5.^24^ In addition, shewasin D was found localized to the pH-neutral cytosol in the native organism.^25^ Our results reconcile these seemingly contradictory observations by demonstrating that, while SamP is still active under acidic conditions, it is more stable and active on its native substrate at near-neutral pH. While most A1 family peptidases are only active at acidic pHs, some are active in near-neutral conditions.^26,27^ This shift is proposed to be caused by a Thr to Ala substitution near the active site, which is found in both shewasins (Fig. S28).^26^

**Figure 4:**
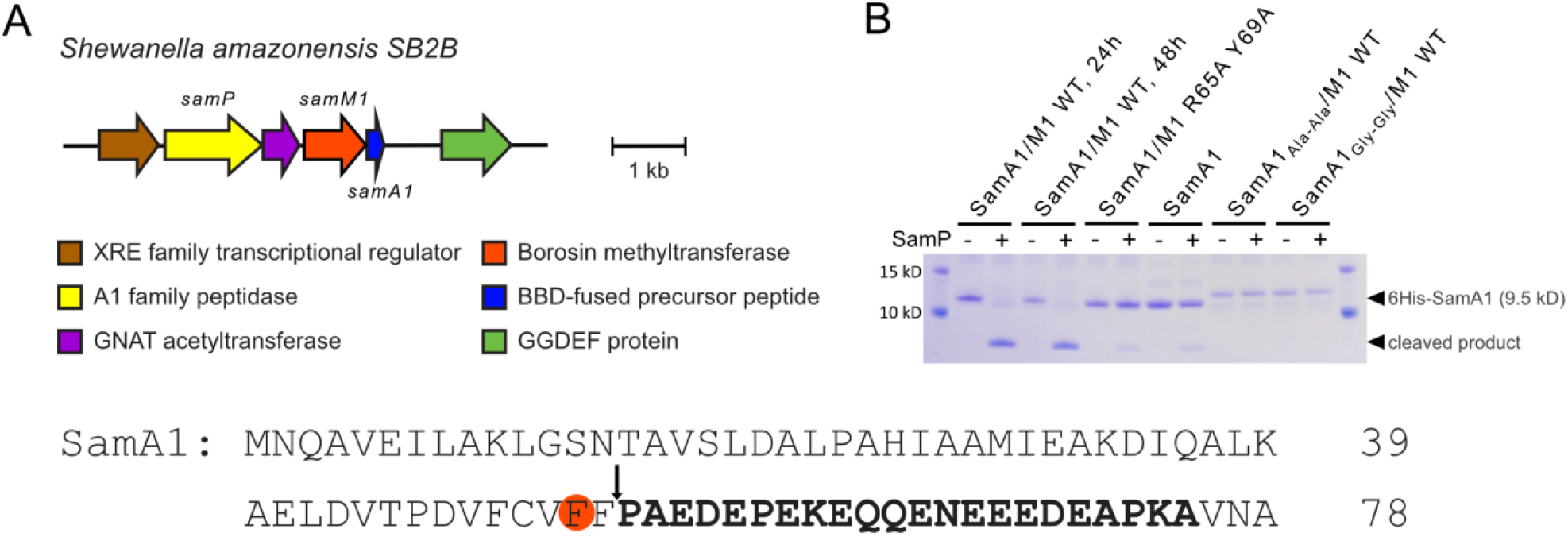
SamP acts as a borosin RiPP protease. A. Putative *S. amazonensis* borosin BGC. Genes expressed in this study are labeled. The amino acid sequence of the precursor peptide SamA1 is shown (bottom) with the putative core sequence in bold. LC-MS/MS localized α-*N-* methylated Phe52 is circled in orange, with an arrow denoting the SamP cut site. B. SDS-PAGE of *in vitro* reactions containing methylated (lanes 2-5) and unmethylated (lanes 6-9) SamA1 and variants (lanes 10-13) with and without SamP (+/-).

Finally, LC-MS/MS analysis of *in vitro* reactions localized SamP cleavage at the amide bond between Phe53 and Pro54 (Fig. S27). Previous work probing SamP substrate recognition found a strong preference for Leu, Phe, Tyr, and Asp at the P1 position as well as bulky, hydrophobic residues at P2.^25^Moderate preference for neutral residues at positions P3/P1’/P2’ and Asp at P4’ were also observed. The SamA1 cleavage site is mostly consistent with these results (Fig. 4). Notable exceptions are Pro at the P1’ position and Glu at P3’. Though previous studies observed no preference for these residues at these positions, there was also no indication of disfavor. Given the strong preference for bulky residues at P2 and P1, substrate variants replacing the Phe-Phe motif in the core peptide with Ala-Ala (SamA1_Ala-Ala_) or Gly-Gly (SamA1_Gly-Gy_) were assayed with SamP. A small amount of proteolytic activity was observed for SamA1_Ala-Ala_, but not for SamA1_Gly-Gly_ (Fig. S27, Fig. S29). LC-MS/MS analysis of trypsinized SamA1_Ala-Ala_ and SamA1_Gly-Gly_, revealed >97% and >72% methylation, respectively (Fig. S24-S26). This indicates that α-*N-*methylation is not solely sufficient for SamP recognition. Further work detailing SamP substrate scope and specificity is ongoing and will be reported elsewhere.

In attempts to identify SamP activity in the native host, LC-MS/MS analysis was performed on extracts from *S. amazonensis* cultures overexpressing a plasmid-encoding copy of the *sam* BGC. The peptide PAEDEPEKEQQENEEEDEAPKA was detected in cell extracts, corresponding to the C-terminus of SamA1 sans three C-terminal residues presumably removed by a host protease (Fig S30). These results support our SamP *in vitro* results and confirm α-*N*-methylation is not found in the proteolytically released C-terminal core peptide. Of note, a putative acetyltransferase (PF13302) is also encoded in the *sam* BGC, suggesting additional modifications may be present in the final natural product (Fig. 4). Future work will delve further into the biological activity and final structure of the *S. amazonensis* borosin metabolite, since studies have shown that anionic peptides can have specific and potent antimicrobial activities.^28,29^ Nonetheless, our data redefines the activity profile and substrate scope of rare bacterially encoded pepsins, now revealed as borosin pathway maturases. To our knowledge, this is the first example of a RiPP leader peptide modification required as a recognition element for proteolytic release of the natural product.

## Conclusions

Borosins are a rapidly expanding RiPP class with diverse core sequences and biosynthetic protein domain architectures. Several factors, including the indiscriminate nature of the class-defining backbone α-*N*-methylation, have necessitated time-consuming manual inspection to identify putative borosin precursor peptides. Our newly developed borosin RODEO module has enabled a streamlined approach for borosin BGC analysis, including precursor peptide identification. This large-scale identification allowed for evolutionary analysis of borosin BGCs which suggest borosin methyltransferases evolved once, then diverged into two main groups. The divide between the Pseudomonadota-dominated group containing BBD-fused precursor peptides and the taxonomically diverse group containing BBD-fused methyltransferase may correlate with differences in biological function; further studies on the bioactivities of the peptide products may shed light on the benefit and pattern of propagation of borosin BGCs. Additionally, analysis of >2,500 RODEO-mined borosin BGCs revealed a new type of fused borosin with a core sequence N-terminal to the methyltransferase domain. Biochemical characterization showed this type 0 borosin methyltransferase catalyzes intramolecular α-*N*-methylation unlike all other fused borosin methyltransferases studied to date. Finally, we demonstrated that the previously studied pepsin-like aspartic protease SamP (shewasin A) is involved in borosin RiPP maturation, marking only the second characterized borosin protease. *In vitro* and mutagenesis experiments revealed that SamP preferentially cleaves an α-*N-*methylated precursor peptide SamA1 to yield a highly anionic yet unmethylated peptide. While ongoing work will be necessary to confirm the final natural product and details of SamP specificity, these data report the first instance of a post-translational modification requirement on a RiPP leader peptide for natural product maturation. All in all, our newly developed RODEO module provides a new large-scale computational tool for studying borosin BGCs and demonstrates its utility to facilitate targeted enzyme and natural product discoveries.

## Materials and Methods

### Materials

HiFi DNA Assembly Master Mix, restriction enzymes, PNK, T4 ligase, OneTaq and Q5 High Fidelity DNA polymerase were purchased from New England Biolabs (NEB). Gene synthesis and codon optimization was performed by Twist Biosciences. Commercial proteases were purchased from Promega (sequencing-grade trypsin, AspN, GluC and chymotrypsin) or Gold Biotechnology (proteinase K). Primers were ordered from Integrated DNA Technologies. Unless otherwise stated, chemicals and reagents were purchased from Millipore Sigma.

*S. amazonensis* SB2B was obtained from Dr. Jeffrey Gralnick.

### Bioinformatic mining and analysis of borosin biosynthetic gene clusters (BGCs)

#### Borosin module

To generate a list of potential borosin BGCs, we used characterized borosin NMT sequences as queries for protein BLAST searches against the NCBI non-redundant (nr) database (Table S1). High confidence borosin BGCs were assigned by manually inspecting BGCs for precursor peptides and protein architectures similar to those already discovered. The methyltransferase and precursor peptide sequences were analyzed, and HMMER3^1^ was used to create custom profile Hidden Markov Models (HMMs) (Supplementary dataset 1) to identify more distant homologs. High confidence precursor peptides were analyzed for motifs using XSTREME from the MEME suite^17^. These features were used alongside amino acid content for heuristic scoring and support vector machine (SVM) classification (additional details found in SI).

#### Bioinformatic analysis of borosin BGCs

To generate a list of potential borosin BGCs for RODEO to analyze, we used characterized borosin NMT sequences as queries for BLAST-P searches against the NCBI non-redundant (nr) database (Table S3) keeping up to 10,000 hits for each query. Additionally, hmmsearch^1^ was used to query the UniProtKB database for each of the borosin NMT, BBD and precursor HMMs; these sequences were mapped to their corresponding NCBI accession. After removing duplicates, the gathered sequences were run through the borosin RODEO module. BGCs with multiple queried accessions were dereplicated and precursors with a cutoff score of 16 or higher were considered valid borosin precursors and included in the final borosin dataset.

### Cloning and gene synthesis

Genes *samA1, samM1*, and *samP*, were directly amplified from genomic DNA of *Shewanella amazonensis* SB2B using gene-specific primers (Supp. Dataset 4) and cloned into pET28a via Gibson assembly (NEBuilder® HiFi DNA Assembly Master Mix, NEB) using manufacturer’s protocols. The vector for co-expression of SamA1 and SamM1 was constructed using restriction enzyme-mediated ligation by digesting pCDFDuet-1 (NcoI-HF/HindIII-HF), pET28a-6His-*samA1* (NcoI-HF/HindIII-HF), and pET28-6His-*samM1* (NdeI-HF/XhoI-HF) followed by gel excision, column purification, and assembly by T4 Ligation (NEB). Gene synthesis for *skoAM* and *fpeAM* was performed by Twist Biosciences. Sequences were codon-optimized for expression in *E. coli* and designed to include hexahistidine or SUMO-tags. Site-directed mutagenesis for the *samA1, samP*, and *skoAM* mutants were performed by PCR using primers containing mutations (Supp. Dataset 4), followed by *DpnI* digestion, column cleanup, and single-tube *PNK* and T4 ligase reaction.

### Protein expression and purification

Protein expression and purification were performed as described previously. Briefly, genes were expressed in LOBSTR (Kerafast) *E*.*coli* or BL21(A1) *E. coli* in cases of *samA1/samM1* coexpression, at 16 °C for 24, 48, or 72h. Cells were harvested by centrifugation, resuspended in lysis buffer (50 mM HEPES pH 8.0, 300 mM NaCl, 25 mM imidazole, 10% (v/v) glycerol, 1 mg/mL lysozyme) and lysed by sonication on ice. Recombinant proteins were purified via nickel-affinity chromatography based on the manufacturer’s recommendations (Ni-NTA resin, Gold Biotechnology). Size exclusion chromatography of 6His-SkoAM was performed on a Superdex™ 75 column (Cytiva Life Sciences) using the running buffer 50 mM HEPES pH 8.0, 150 mM NaCl, 10% (v/v) glycerol over 120 minutes at a flowrate of 1 mL/min.

### *In vitro* assays of SamA1 and SamP

Purified 6His-SamA1 (with or without SamM1 in complex) was added to a tube with SamP at ratios ranging from 1:10 – 1:1,000 (enzyme: substrate). Reactions were performed in 50-100 mM sodium citrate pH 5.0, MES pH 6.0, HEPES pH 7.0 or HEPES pH 8.0 with 100 mM KCl, and with or without 8% DMSO. 50 mM MES pH 6.0 was used for subsequent assays once the optimal conditions were identified. For SDS-PAGE analysis, reactions were quenched by adding 1/4 volume 5X SDS-PAGE loading dye (250 mM TrisCl pH 6.8; 5% (v/v) beta-mercaptoethanol; 10% (w/v) SDS; 0.25% (w/v) bromophenol blue; 50% (v/v) glycerol) and boiling for 5 minutes before running on SDS-PAGE. For LC-MS/MS analysis, reactions were quenched by adding 1M ammonium bicarbonate solution, pH 8.0 to a final concentration of 100 mM. Samples were treated with iodoacetamide (described below), desalted using C18 ziptips, then run on LC-MS/MS (described below).

### Proteolytic digestion and peptide mass spectrometric analysis

Proteolytic digestion was performed as previously described.^5^ Briefly, proteins were in-gel digested with protease at enzyme:substrate mass ratios between 1:4 – 1:100 at 37°C o/n. After digestion, samples were desalted using C18 resin ZipTips (EMD Millipore, Burlington, MA). Cysteine-containing proteins were reduced then treated with iodoacetamide before digestion.

LC-MS/MS analysis of peptides was performed on a Thermo Scientific Fusion or Fusion Lumos mass spectrometer equipped with a Dionex Ultimate 3000 UHPLC system using a nLC column (200 mm × 75 μm) packed using Vydac 5-μm particles with a 300 Å pore size (Hichrom Limited). Elution was performed with an increasing linear acetonitrile gradient and data-dependent as well as targeted MS2 was performed using either higher-energy collisional dissociation (HCD) or electron-transfer dissociation (ETD). Methylation state abundance for trypsinated SamA1 variants were calculated by summing the peak areas (as ion counts) from extracted ion chromatograms (EICs) of the theoretical monoisotopic 2+ and 3+ parent masses +/-10 mmu for each methylation state, dividing by the total peak areas of all methylation states for that variant, and multiplied by 100%. Percentages were averaged and standard deviation was calculated from 3 technical replicates for each variant. Additional details can be found in the supplementary materials.

### Structure prediction of SkoAM

Structural predictions of monomeric and homodimeric SkoAM were created using Alphafold v2.1.1.

### *sam*BGC overexpression and metabolite extraction from *S. amazonensis* SB2B

The *sam*BGC was cloned into the pBBR1MCS5 expression vector using methods described above then transformed into the diaminopimelic acid (DAP) auxotrophic *E. coli* WM3064 strain. The plasmid was then conjugated into the *Shewanella amazonensis* SB2B host strain by mating the two strains on DAP-supplemented LB plates followed by selection on DAP-deficient LB supplemented with gentamicin. For expression, 1:100 volume of an overnight culture was inoculated into TB + gentamicin and grown at 30°C; 170 rpm until OD_600nm_ reached 0.4-0.6. Expression was induced with 0.2 mM IPTG and incubated at 30°C; 170 rpm for 24 hours. Cells were harvested by centrifugation at 4,000 x g for 20 minutes at 4°C then flash-frozen in liquid nitrogen and stored at -80°C until extraction. For extraction, the biomass was resuspended in 80% methanol and sonicated on ice for 5 minutes. Debris was removed by centrifugation at 4,000 x g for 30 minutes followed by sequential filtration through Whatman paper then a 0.22 μm filter. The sample was then dried down in a rotary evaporator then resuspended in water. Sample was then processed through a Pierce™ Graphite Spin Column (Thermo Scientific, Waltham, MA) following manufacturer’s instructions then analyzed by LC-MS/MS using methods described above.

## Supporting information

Supplemental dataset 1 - HMM profiles

Supplemental dataset 2 - Borosin precursors

Supplemental dataset 3 - Phylogenetic trees

Supplemental dataset 4 - Cloning primers_genes_plasmids_proteins

## Associated Content

### Accession Codes

FpeAM: WP_083990299.1

SkoAM: WP_018611879.1

LepM1: WP_061716904.1

ParM: WP_007623905.1

SspM: WP_031073184.1

SamM: WP_011758902.1

SonM: WP_011071665.1

RceM: WP_012568691.1

SliM: ADB39711.1

AinM: WP_083827882.1;

PmoM: WP_051555776.1

SurM1: WP_165337934.1

LulM: WP_149193073.1

### Supporting Information

Additional experimental methods, data, & figures (.pdf)

Supplemental Dataset 1: Custom HMMs (.txt)

Supplemental Dataset 2: Precursor Dataset (.xlsx)

Supplemental Dataset 3: Phylogenetic trees in Newick format (.nwk)

Supplemental Dataset 4: Primers, genes, plasmids, and protein sequences and accession codes used in this study (.xlsx)

## Notes

The authors declare the following competing financial interest(s): M.F.F. is an inventor on patents US20190112583A1, WO2017EP58327, and on patent application Nos. PCT/US2021/019009 and 62/979,947.

## Funding

This work was supported by the National Institutes of Health (R01 GM123998 for D.A.M.) and the National Institute of General Medical Sciences (R35 GM133475 for M.F.F.).

## Acknowledgements

We thank L. Daigh for the preliminary work on the borosin module and thank J. Gralnick for the strain *S. amazonensis* SB2B.

## Notes

### Competing Interest Statement

The authors have declared no competing interest.

