## Supporting Information for "Computationally guided exploration of borosin biosynthetic strategies"

**Additional experimental methods, data, & figures**

### Table of Contents

Materials and Methods……….……...………………………………………………………………...S3
Table S1. Accession codes and precursor peptide sequence for characterized borosins...……S8
Table S2.  Features and weights used for borosin precursor peptide scoring.....……….……S13
Table S3. Borosin module support vector machine parameters…...…….…..………………....S15
Figure S1. Expectation value of PF00590 (TP methylase) vs BorosinMT HMMs..…………….S18
Figure S2. Borosin module scoring accuracy……………………..…………..……………......S19
Figure S3. Precursor peptide scoring distribution.…………………………..…...…………...S20
Figure S4. SSN of borosin methyltransferases colored by predicted type.……………….…....S21
Figure S5. Unusual borosin BGC architectures..……………………………………..………S22
Figure S6. Phylogenetic distribution of diversity-maximized subset of RODEO curated
borosin methyltransferases…………………………………………………………………..…S23
Figure S7. Type distribution of diversity-maximized subset of RODEO curated
borosin methyltransferases ………………………………………………………………….S24
Figure S8. Phylogenetic distribution of excised BBDs ..………………...……….........….…S25
Figure S9. Type distribution of excised BBDs.……………………………………….……S26
Figure S10: Sum of branch distances from diversity-maximized tree……...………….…...……S27
Figure S11. Phylogenetic tree of SspM homologs..…………………..…………...………….S28
Figure S12. Type distribution of SspM homologs………….....……………………………..….S29
Figure S13. Sum of branch distances between SspM homologs…………………..…….....…..S30
Figure S14. G+C content analysis of borosin methyltransferases..…………...……………....S31
Figure S15. SSN of borosin methyltransferases with nearby phage integrases.…………….…S32
Figure S16. MS2 analysis of type 0 borosin core peptide regions..……………………………S33
Figure S17. SEC and Alphafold model for SkoAM…..…………………………..................S35
Figure S18. SkoAM trans-complementation assay.……………………...……………………..S36
Figure S19. MS1 spectra for SkoAM *trans*-complementation assay……..……..……………S37
Figure S20. SSN of borosin methyltransferases associated with a PF00026 eukaryotic
aspartyl protease.…………………………………………………………………………...….S39
Figure S21. Unrooted tree of SamP homologs ……………………………..……………...S40
Figure S22. Rooted tree of SamP homologs ………………………….…..……….…………S41
Figure S23. MS2 fragmentation of methylated SamA1…………………….……………......S42
Figure S24. Calculated methylation state abundances of SamA1 variants………………………S44
Figure S25. LC-MS1 EICs for 0-2 methylation states of trypsinized SamA1 variants…………S45
Figure S26. MS2 fragmentation of SamA1 variants………………………………………….....S49
Figure S27. SDS-PAGE gels of SamP *in vitro* reactions……………..………………………S60
Figure S28. Alignment of A1 peptidase family members..………………………..….........S63
Figure S29. LC-MS/MS analysis of SamP *in vitro* reactions..…...…………………………S64
Figure S30. LC-MS/MS analysis of metabolite extract from *sam*BGC overexpression………...S66
References…………………………………………………………………………………..…S68

### Materials and Methods

#### Materials

HiFi DNA Assembly Master Mix, restriction enzymes, PNK, T4 ligase, OneTaq and Q5 High Fidelity DNA polymerase were purchased from New England Biolabs (NEB). Gene synthesis and codon optimization was performed by Twist Biosciences. Commercial proteases were purchased from Promega (sequencing-grade trypsin, AspN, GluC and chymotrypsin) or Gold Biotechnology (proteinase K). Primers were ordered from Integrated DNA Technologies. Unless otherwise stated, chemicals and reagents were purchased from Millipore Sigma. *S. amazonensis* SB2B was obtained from Dr. Jeffrey Gralnick.

#### Bioinformatic mining and analysis of borosin biosynthetic gene clusters (BGCs)

*Borosin module.* To generate a list of potential borosin BGCs, we used characterized borosin NMT sequences as queries for protein BLAST searches against the NCBI non-redundant (nr) database (Table S1). High confidence borosin BGCs were assigned by manually inspecting BGCs for precursor peptides and protein architectures similar to those already discovered. The methyltransferase and precursor peptide sequences were analyzed, and HMMER3[1] was used to create custom profile Hidden Markov Models (HMMs) (Supplementary dataset 1) to identify more distant homologs. High confidence precursor peptides were analyzed for motifs using XSTREME from the MEME suite[2]. These features were used alongside amino acid content for heuristic scoring and support vector machine (SVM) classification.

The SVM was trained and tested with 414 positives and 7,838 negatives using a 5-fold cross validation. The positive dataset includes characterized and high confidence predicted borosin precursor peptides. The negative dataset includes hypothetical non-borosin open reading frames (ORFs) from within borosin and non-borosin RiPP BGCs. The borosin scoring module accuracy was assessed using the following equations (Fig. S2, Fig. S3):

Equation S1:

$$Precision = \frac{true\ positives}{true\ positives + false\ positives}$$

Equation S2:

$$Recall = \frac{true\ positives}{true\ positive + false\ negative}$$

Equation S3:

$$f1\ score = 2 * \frac{recall * precision}{recall + precision}$$

*Bioinformatic analysis of borosin BGCs.* To generate a list of potential borosin BGCs for RODEO to analyze, we used characterized borosin NMT sequences as queries for BLASTP searches against the NCBI non-redundant (nr) database (Table S3) keeping up to 10,000 hits for each query. Additionally, hmmsearch[1] was used to query the UniProtKB database for each of the borosin NMT, BBD and precursor HMMs; these sequences were mapped to their corresponding NCBI accession. After removing duplicates, the gathered sequences were run through the borosin RODEO module. BGCs with multiple queried accessions were dereplicated and precursors with a cutoff score of 16 or higher were considered valid borosin precursors and included in the final borosin dataset.

An analysis of the phylogenetic distribution of borosin methyltransferases with a RODEO predicted precursor peptide was performed. All sequence similarity networks (SSNs) were generated using the Enzyme Function Initiative Enzyme Similarity Tool (EFI-EST) (https://efi.igb.illinois.edu/)[3]. SSN visualization was performed using organic layout within Cytoscape[4]. Multiple sequence alignments were performed using MUSCLE[5], and phylogenetic trees were built using the FastTree[6] with the default Jones-Taylor-Thornton model. Trees were visualized using the interactive Tree of Life (iTOL) website (http://itol.embl.de/)[7]. The sum of branch lengths were compared using MEGA 11[8]. The protein sequence of OphMA, PgiMA1, and AboMA are not available for the UniProtKB or NCBI nr databases and were added manually to SSNs and trees.

### Cloning and gene synthesis

Genes *samA1*, *samM1*, and *samP* were directly cloned from genomic DNA of *Shewanella amazonensis* SB2B. Genomic DNA was extracted after the bacteria were grown overnight in LB media. Genes were PCR amplified with Q5 High-Fidelity DNA Polymerase (NEB) using standard conditions (1x Q5 Reaction buffer, 1x Q5 High GC Enhancer buffer, 200 µM dNTPs, 0.5 µM final concentration of each primer, 0.04 U Q5 polymerase / 50-µL reaction) with gene-specific primers (Table S1). The verified PCR products were excised, purified (Monarch DNA Gel Extraction Kit, NEB), and cloned into pET28a via Gibson assembly (NEBuilder® HiFi DNA Assembly Master Mix, NEB) using manufacturer's protocols. The vector for coexpression of *samA1* and *samM1* was constructed by digesting pCDFDuet-1 (NcoI-HF/HindIII-HF), pET28a-6His-*samA1* (NcoI-HF/HindIII-HF), and pET28-6His-*samM1* (NdeI-HF/XhoI-HF) followed by gel excision, column purification, and assembly by T4 ligation (NEB). Gene synthesis for *skoAM* and *fpeAM* was performed by Twist Biosciences. Sequences were codon-optimized for expression in *E. coli* and designed to include hexahistidine or SUMO-tags. Site-directed mutagenesis for the *samA1*, *samP*, and *skoAM* mutants were performed by PCR using primers containing mutations (Table S1), followed by DpnI digestion, column cleanup, and single-tube *PNK* and T4 ligase reaction.

### Protein expression and purification

Protein expression and purification were performed as described previously. Briefly, genes were expressed in LOBSTR (Kerafast) *E.coli* or BL21(A1) *E. coli* in cases of *samA1/samM1* co-expression, at 16 °C for 24, 48, or 72h. Cells were harvested by centrifugation, resuspended in lysis buffer (50 mM HEPES pH 8.0, 300 mM NaCl, 25 mM imidazole, 10% (v/v) glycerol, 1 mg/mL lysozyme) and lysed by sonication on ice. Recombinant proteins were purified via nickel-affinity chromatography based on the manufacturer's recommendations (Ni-NTA resin, Gold Biotechnology). Size exclusion chromatography of 6His-SkoAM was performed on a Superdex™ 75 column (Cytiva Life Sciences) using the running buffer 50 mM HEPES pH 8.0, 150 mM NaCl, 10% (v/v) glycerol over 120 minutes at a flowrate of 1 mL/min.

### *In vitro* assays of SamA1 and SamP

Purified 6His-SamA1 (with or without SamM1 in complex) was added to a tube with SamP at ratios ranging from 1:10 – 1:1,000 (enzyme:substrate). Reactions were performed in 50-100 mM sodium citrate pH 5.0, MES pH 6.0, HEPES pH 7.0 or HEPES pH 8.0 with 100 mM KCl, and with or without 8% DMSO. 50 mM MES pH 6.0 was used for subsequent assays once the optimal conditions were identified. For SDS-PAGE analysis, reactions were quenched by adding 1/4 volume 5X SDS-PAGE loading dye (250 mM TrisCl pH 6.8; 5% (v/v) β-mercaptoethanol; 10% (w/v) SDS; 0.25% (w/v) bromophenol blue; 50% (v/v) glycerol) and boiling for 5 minutes before running on SDS-PAGE. For LC-MS/MS analysis, reactions were quenched by adding 1 M ammonium bicarbonate solution, pH 8.0 to a final concentration of 100 mM. Samples were treated with iodoacetamide (described below), desalted using C18 ziptips, then run on LC-MS/MS (described below).

**Proteolytic digestion**

Proteolytic digestion was performed as previously described. Briefly, trypsin gold, chymotrypsin, AspN, GluC (Promega) or proteinase K (Gold Biotechnology) was used with either an in-solution or an in-gel digestion method. For in-solution digestion, an aliquot of protein containing 10-15 µg of protein-of-interest was transferred to a 1.5 mL protein LoBind tube and dried using a vacufuge concentrator (Eppendorf). Samples were resuspended in 25 µL of a solution containing 8 M urea, 0.5 M ammonium biocarbonate, pH 8.0 and 4 mM DTT. After brief vortexing (5 seconds) and centrifugation, the sample was incubated at 37 °C for 45 min. If the core peptide sequences contained cysteine residues, an alkylation treatment was performed as follows. After letting samples cool at room temperature for five minutes, 25 µl of 20 mM iodoacetamide (IAA) solution in ultrapure water was added and mixed by pipetting. Samples were incubated at room temperature in the dark for 30 minutes. Samples were then diluted with 50 µL of LC-MS-grade dd$H_2O$ to dilute urea below concentrations that would inhibit protease activity (< 2 M urea). Protease was added for a final protease:substrate (mass:mass) ratio between 1:25 – 1:100. Samples were incubated at 37 °C (or 25 °C for chymotrypsin) overnight. Extracted peptides were frozen at -80 °C for 30 min to deactivate the protease. Peptide solutions were then thawed and dried using a vacufuge concentrator (Eppendorf). Peptides were resuspended in 0.1% formic acid (FA) and further purified and desalted using C18 ZipTips (Millipore Sigma) according to the manufacturer's specifications. After drying the samples again, peptides were resuspended in 12-35 µL of 20% acetonitrile (ACN), 0.1% FA, and transferred to glass vials for MS analysis.

For in-gel digestion, bands from soluble fractions were excised from SDS-PAGE gels stained with Coomassie Blue and cut into ~2 mm x 2 mm cubes. Gel pieces were transferred to 1.5 mL Protein LoBind tubes (Eppendorf) and then washed with 50 mM ammonium bicarbonate (ABC), 50% v/v ACN three times until all the stain was removed. Gel pieces were then dehydrated in 100% ACN until semi-opaque (~30 sec), after which the ACN was discarded. If the putative RiPP core sequences contained cysteines, reduction (treatment with 10 mM DTT in a 65 °C water bath for 1 h before discarding DTT solution) and alkylation (treatment with 55 mM iodoacetamide in 50 mM ABC at room temperature for 30 min) were performed. After reduction and alkylation, gel pieces were washed twice using 50 mM ABC, 50% (v/v) ACN and then dehydrated in 100% ACN until semi-opaque as in the previous steps. Gel pieces were then rehydrated in digestion buffer (50 mM ABC, 5 mM $CaCl_2$, and appropriate units of protease) for 15 min before overnight incubation at 37 °C or 25 °C (chymotrypsin only).

Trypsin digests were performed in a 1:40 protease:protein ratio, whereas proteinase K digests were performed using a ~1:4 – 1:6 (mass:mass) ratio. Digestion supernatants were recovered the next day, and placed in fresh LoBind tubes. Cleaved peptides were recovered by dehydrating the gel pieces in three successive steps. First, ~60 µL of 50% ACN and 0.3% FA was added, incubated for 15 min at room temperature and recovered. Second, ~60 µL of 80% ACN and 0.3% FA was added, incubated and recovered. Finally, ~60 uL of 95% ACN and 0.1% FA was added, incubated, and recovered. The extracted peptides were pooled and frozen at -80 °C for 30 min to deactivate the protease. Peptide solutions were then thawed and dried using a vacufuge concentrator (Eppendorf). Peptides were resuspended in 0.1% FA and further purified and desalted using C18 ZipTips (Millipore Sigma) according to the manufacturer's specifications. After drying the samples again, peptides were resuspended in 12-35 μL of 20% ACN, 0.1% FA, and transferred to glass vials for MS analysis.

**Peptide mass spectrometric analysis**

LC-MS/MS analysis of digested peptides was performed as described previously. Briefly, data were recorded on a Thermo Scientific Fusion or Fusion Lumos mass spectrometer equipped with a Dionex Ultimate 3000 UHPLC system using a nLC column (200 mm × 75 μm) packed using Vydac 5-μm particles with a 300 Å pore size (Hichrom Limited). Elution was performed with a linear gradient using water with 0.1% FA (solvent A) and ACN with 0.1% FA (solvent B) at a flow rate of 0.3 μL $min^{-1}$. The column was equilibrated with 20% solvent B for 5 min, followed by a linear increase of solvent B to 95% over 32 min and a final elution step with 95% solvent B for 2 min. Mass spectra were acquired in positive ion mode. Full MS was done at a resolution of 60,000 [automatic gain control (AGC) target, $4 \times 10^5$; maximum injection time (IT), 50–100 ms; range, 300–1800 m/z], and data-dependent as well as targeted MS/MS was performed at a resolution of 15,000 (AGC target, $5 \times 10_5$; maximum IT, 100–500 ms; isolation window, 2.2 m/z) using higher-energy collisional dissociation (HCD) or electron-transfer dissociation (ETD). Data were processed using Thermo Fisher Xcalibur software and MaxQuant v1.6.10.145. Methylation state abundance for trypsinized SamA1 variants were calculated by summing the peak areas (as ion counts) from extracted ion chromatograms (EICs) of the theoretical monoisotopic $2^+$ and $3^+$ parent masses +/- 10 amu for each methylation state, dividing by the total peak areas of all methylation states for that variant, and multiplied by 100%. Percentages were averaged and standard deviation was calculated from three technical replicates for each variant.

**Structure prediction of SkoAM**

Structural predictions of monomeric and homodimeric SkoAM were created using Alphafold v2.1.1.

***sam*BGC overexpression and metabolite extraction from *S. amazonensis* SB2B**

The *sam*BGC was cloned into the pBBR1MCS5 expression vector using methods described above then transformed into the diaminopimelic acid (DAP) auxotrophic *E. coli* WM3064 strain. The plasmid was then conjugated into the *S. amazonensis* SB2B host strain by mating the two strains on DAP-supplemented LB plates followed by selection on DAP-deficient LB supplemented with gentamicin. For expression, 1:100 volume of an overnight culture was inoculated into TB + gentamicin and grown at 30 °C; 170 rpm until $OD_{600}$ reached 0.4-0.6. Expression was induced with 0.2 mM IPTG and incubated at 30 °C; 170 rpm for 24 h. Cells were harvested by centrifugation at 4,000 x g for 20 min at 4 °C then flash-frozen in liquid nitrogen and stored at -80 °C until extraction. For extraction, the biomass was resuspended in 80% methanol and sonicated on ice for 5 min. Debris was removed by centrifugation at 4,000 x g for 30 min followed by sequential filtration through Whatman paper and then a 0.22 μm filter. The sample was then dried down in a rotary evaporator then resuspended in water. Sample was then processed through a Pierce™ Graphite Spin Column (Thermo Scientific, Waltham, MA) following manufacturer's instructions then analyzed by LC-MS/MS using methods described above.

**Table S1. Accession codes and precursor peptide sequence for characterized borosins.** Sequences for the associated methyltransferases were used as queries for a protein BLAST search to yield the input for the RODEO module.

| Name | NCBI Protein Accession Code | Precursor peptide sequence | Ref |
|---|---|---|---|
| OphMA | N/A | METSTQTKAGSLTIVGTGIESIGQMTLQALSYIEAAAKVFY<br>CVIDPATEAFILTKNKNCVDLYQYYDNGKSRLNTYTQMSE<br>LMVREVRKGLDVVGVFYGHPGVFVNPSHRALAIAKSEGY<br>RARMLPGVSAEDCLFADLCIDPSNPGCLTYEASDFLIRDR<br>PVSIHSHLVLFQVGCVGIADFNFTGFDNNKFGVLVDRLEQ<br>EYGAEHPVVHYIAAMMPHQDPVTDKYTVAQLREPEIAKRV<br>GGVSTFYIPPKARKASNLDIIRRLELLPAGQVPDKKARIYPA<br>NQWEPDVPEVEPYRPSDQAAIAQLADHAPPEQYQPLATS<br>KAMSDVMTKLALDPKALADYKADHRAFAQSVPDLTPQER<br>AALELGDSWAIRCAMKNMPSSLLDAARESGEEASQNGFP<br>WVIVVGVIGVIGSVMSTE | 9,10 |
| DbiMA1 | THU83556.1 | MESSTQTKPGSLIVVGTGIESIGQMTLQALSYIEAASKVFY<br>CVIDPATEAFILTKNKNCVDLYQYYDNGKSRMDTYTQMAE<br>LMLKEVRNGLDVVGVFYGHPGVFVNPSHRALAIARSEGY<br>QARMLPGVSAEDCLFADLCIDPSNPGCLTYEASDFLIRER<br>PVNVHSHLILFQVGCVGIADFNFSGFDNSKFTILVDRLEQE<br>YGPDHTVVHYIAAMMPHQDPVTDKFTIGQLREPEIAKRVG<br>GVSTFYIPPKARKDINTDIIRLLEFLPAGKVPDKHTQIYPPN<br>QWEPDVPTLPPYGQNEQAAITRLEAHAPPEEYQPLATSK<br>AMTDVMTKLALDPKALAEYKADHRAFAQSVPDLTPQERA<br>ALELGDSWAIRCAMKNMPSSLLEAASQSVEEASMNGFPW<br>VIVTGIVGVIGSVVSSA | 11,12 |
| LedMA | GAW09067.1 | METPTLNKSGSLTIVGTGIESIGQMTLQTLSYIEAADKVFYC<br>VIDPATEAFILTKNKDCVDLYQYYDNGKSRMDTYTQMSEV<br>MLREVRKGLDVVGVFYGHPGVFVNPSLRALAIAKSEGFKA<br>RMLPGVSAEDCLYADLCIDPSNPGCLTYEASDFLIRERPT<br>NIYSHFILFQVGCVGIADFNFTGFENSKFGILVDRLEKEYG<br>AEHPVVHYIAAMLPHEDPVTDQWTIGQLREPEFYKRVGG<br>VSTFYIPPKERKEINVDIIRELKFLPEGKVPDTRTQIYPPNQ<br>WEPEVPTVPAYGSNEHAAIAQLDTHTPPEQYQPLATSKA<br>MTDVMTKLALDPKALAEYKADHRAFAQSVPDLTANERTAL<br>EIGDSWAFRCAMKEMPISLLDNAKQSMEEASEQGFPWIIV<br>VGVVGVVGSVVSSA | 12,13 |
| CmiMA | TEB34317.1 | MIGASLAKKGQLTIVGSGIASISHLTLQAVSAIENADIVCYV<br>VADGATEAFIRKKNPNSLDLYHLYGEDKQRTDTYIQMAEF<br>MLIRVRQGQNVVGVFYGHPGVFVCPTHRALYIARSEGYK<br>ARMLPGLSAEDCLFADLGIDPSSVGCVTYEATDLLVFKRPI<br>NPASHLVLYQVGIVGKSNFKFDYTSDENIHFTKLLDRLEEA<br>YGPEHSVTHYIAPLFPTEDPIAEEYTIAQLRLPEIRDKIHTIS<br>TFYVPPKTSESLIYDEVLLASLGVTHKPSVPYPWNPEATPY<br>GPREKKAIELLAEHEPPKGYRPLKERSGLLAVLEKLCLEPL<br>EMKKYNEDRQAYADGLKGLTENEKEALVKGDHRTLAGAL<br>KVGDTPTNPAALVFTFIITRLD | 13 |

| | | | |
|---|---|---|---|
| CmaMA | TFK23735.1 | MDATANPKAGQLTIVGSGIASINHMTLQAVACIETADVVCY<br>VVADGATEAFIRKKNENCIDLYPLYSETKERTDTYIQMAEF<br>MLNHVRAGKNVVGVFYGHPGVFVCPTHRAIYIARNEGYR<br>AVMLPGLSAEDCLYADLGIDPSTVGCITYEATDMLVYNRP<br>LNSSSHLVLYQVGIVGKADFKFAYDPKENHHFGKLIDRLEL<br>EYGPDHTVVHYIAPIFPTEEPVMERFTIGQLKLKENSDKIAT<br>ISTFYLPPKAPSAKVSLNREFLRSLNIADSRDPMTPFPWNP<br>TAAPYGEREKKVILELESHVPPPGYRPLKKNSGLAQALEK<br>LSLDTRALAAWKTDRKAYADSVSGLTDDERDALASGKHA<br>QLSGALKEGGVPMNHAQLTFFFIISNL | 13,14 |
| SveMA | PVG01141.1 | MASSTHPKRGSLTIAGTGIATLAHMTLETVSHIKEADKVYYI<br>VTDPVTQAFIEENAKGPTFDLSVYYDADKYRYTSYVQMAE<br>VMLNAVREGCNVLGLFYGHPGIFVSPSHRALAIAREEGYE<br>ARMLPGVSAEDYMFADLGLDPALPGCVCYEATNFLIRNKP<br>LNPATHNILWQVGAVGITAMDFENSKFSLLVDRLERDLGP<br>NHKVVHYVGAVLPQSATIMETYTIAELRKPEVIKRISTTSST<br>FYIPPRDSEAIDYDMVARLGIPPEKYRKIPSYPPNQWAGP<br>NYTSTPAYGPEEKAAVSQLANHVVPNSYKTLHASPAMKK<br>VMIDLATDRSLYKKYEANRDAFVDAVKGLTELEKVALKMG<br>TDGSVYKVMSATQADIELGKEPSIEELEEGRGRLLLVVITA<br>AVVV | 13,14 |
| GjuMA | NA | MATPIATTTNTPTKAGSLTIAGSGIASVGHITLETLAYIKESH<br>KVFYLVCDPVTEAFIQENGKGPCINLSIYYDSQKSRYDSYL<br>QMCEVMLRDVRNGLDVLGVFYGHPGVFVSPSHRAIALAR<br>EEGFNAKMLAGVSAEDCLFADLEFDPASFGCMTCEASEL<br>LIRNRPLNPYIHNVIWQVGSVGVTDMTFNNNKFPILIDRLE<br>KDFGPNHTVIHYVGRVIPQSVSKIETFTIADLRKEEVMNHF<br>DAISTLYVPPRDISPVDPTMAEKLGPSGTRVEPIEAFRPSL<br>KWSAQNDKRSYAYNPYESDVVAQLDNYVTPEGHRILQGS<br>PAMKKFLITLATSPQLLQAYRENPSAIVDTVEGLNEQEKYG<br>LKLGSEGAVYALMSRPTGDIAREKELTNDEIANNHGAPYA<br>FVSAVIIAAIICAL | 13 |
| CeuMA | NA | MATQKSGSLTIAGSGIASIGHITLETLSYIEQADKVYYAVAD<br>PATEAFIQDKSKVECFDLTVYYDKDKIRFETYIQMSEVMLR<br>DVRAGHSVLGIFYGHPGVFVCPSHRAIAIALSEGYKARML<br>PGISAEDYMFSDIGFDPALPGCTTQEATHLLLHNKKLDPS<br>MHNIIWQVGGVGADTMNFDNRQFHQLVDCLERDFGSSH<br>KVVHYIGAVMPQSTTIMDEFSIADLRKEEVVKQFTTWSTFY<br>IPPRDAAPVDEGIMQSLGLSSNDMQYTMYPPSSTMRLGIR<br>SPNLDVYGRAGRAAIEKLDHHTPAARHQVLRASPAIRKFM<br>EDLALKSDLRDRYKADPHTVLDAIPGLTSQEKIALGFGKPG<br>PVYKVMRATGRETADGQEHVPHDLTTTDEPGAPVLLLLLL<br>QTT | 13 |
| MroMA1 | NA | MALKKPGSLTIAGSGIASIGHITLETLALIKEADKIFYAVTDP<br>ATECYIQENSRGDHFDLTTFYDTNKKRYESYVQMSEVML<br>RDVRAGRNVLGIFYGHPGVFVAPSHRAIAIAREEGFQAKM<br>LPGISAEDYMFADLGFDPSTYGCMTQEATELLVRNKKLDP<br>SIHNIIWQVGSVGVDTMVFDNGKFHLLVERLEKDFGLDHKI<br>QHYIGAILPQSVTVKDTFAIRDLRKEEVLKQFTTTSTFYVPP<br>RTPAPIDPKAVQALGLPATVTKGAQDWTGFQSVSPAYGP | 13,14 |

| | | | |
|---|---|---|---|
| | | DEMRAVAALDSFVPSQEKAVVHASRAMQSLMVDLALRPALLEQYKADPVAFANTRNGLTAQEKFALGLKKPGPIFVVMRQLPSAIASGQEPSQEEIARADDATAFIIIYIVQG | |
| MroMA2 | NA | MALNKPGSLTIAGSGIASIGHITLETLALIKEADKIFYAVTDPATECYIQENSRGDHFDLTTFYDTNKKRYESYVQMSEVMLREVRAGRNVLGIFYGHPGVFVAPSHRAIAIAREEGFQAKMLPGISAEDYMFADLGFDPSTQGCMTQEATELLVRNKKLDPSVHNIIWQVGSVGVDTMVFDNGKFHLLVERLEKDFGLDHKIQHYIGAILPQSVTVKDAFAIRDLRKEEVLKQFTTTSTFYIPPRAPAPIDAKVLQALGLPPPAQATKDRTGYGPLEKQAVAALDSFIPSQEKQVVHASPAMQSLMADLALRPALFEQYKADPVGFANTRNLNGLTAQEKFALGFNKSGPIFAVMRHLPSAIASGQERSQEEIAHAADDKELLALVVVIVQ | 14 |
| PocMA | NA | MPVSTTTTKNGTLVIAGSGIASIAHITLETLSHIKESDRVYYIVGDPATEAFIQDNASGTCFDLTIFYDTNKVRYDSYVQMCEVMLRDVRAGHTVLGVFYGHPGVFVSPSHRAIAIARDEGYKARMLPGVSAEDYLFADLGFDPATHGCTSYEATDLLVRNKPLNASTHNIIWQVGGVGVGTMVFDNAKFHLLVDRLEKDFGPSHTVVHYIGAVLPQSITTMDKLTIADLRKDAVVKQFNPTSTFYIPPRDISLPLDTMAKKLGMDDASARPVSLYPPSRWTGTKFTTAPAYGPREKDVIAKIDTYAAPKDHKILHASRSMKKLMTDLALNPKLLEKYRANTKAVVEATEGLSAQEKAALNMDLAGPVHAVMKATPSDITDGREMSVDAVASATEPSAALILLLV | 13 |
| PgiMA1 | NA | MSSASSDSNTGSLTIAGSGIASVRHMTLETLAHVQEADIVFYVVADPVTEAYIKKNARGPCKDLEVLFDKDKVRYDTYVQMAETMLNAVREGQKVLGIFYGHPGVFVSPSRRALSIARKEGYQAKMLPGISSEDYMFADLEFDPAVHGCCAYEATQLLLREVSLDTAMSNIIWQVGGVGVSKIDFENSKVKLLVDRLEKDFGPDHHVVHYIGAVLPQSATVQDVLKISDLRKEEIVAQFNSCSTLYVPPLTHANKFSGNMVKQLFGQDVTEVSSALCPTPKWAAGSHLGDVVEYGPREKAAVDALVEHTVPADYRVLGGSLAFQQFMIDLALRPAIQANYKENPRALVDATKGLTTVEQAALLLRQPGAVFGVMKLRASEVANEQGHPVAPASLDHVAFTAPSPASLDHVAFSAPNPASLDHVAFIAPTPASLDHVAFSAPTPASLDHVSFGTPTSASLDHVAFEAPVPASLDHVAFAAPVPASLDHVAFAAPTPASLDHVAFAAPTPASLDHVAFAVPVPASLDHIAFSVPTPASLDHVAFAVPVPDHVAGIPCM | 13 |
| AboMA | NA | MSSPAVETKVPASPDVTAEVIPAPPSSHRPLPFGLRPGKLVIVGSGIGSIGQFTLSAVAHIEQADRVFFVVADPATEAFIYSKNKNSVDLYKFYDDKKPRMDTYIQMAEVMLRELRKGYSVVGVIYGHPGVFVTPSHRAISIARDEGYSAKMLPGVSAEDNLFADIGIDPSRPGCLTYEATDLLLRNRTLVPSSHLVLFQVGCIGLSDFRFKGFDNINFDVLLDRLEQVYGPDHAVIHYMAAVLPQSTTTIDRYTIKELRDPVIKKRITAISTFYLPPKALSPLHEESAAKLGLMKAGYKILDGAQAPYPPFPWAGPNVPIGIAYGRRELAAVAKLDSHVPPANYKPLRASNAMKSTMIKLATDPKAFAQYSRNPALLANSTPGLTTPERKALQTGSQGLVRSVMKTSPEDVAKQFVQAELRDPTLAKQYSQECYDQTGNTDGIAVISAWLKSKGYDTTPTAINDAWADMQANSLDVYQSTYNTMVDGKSGPAITIKSGVVYIGNTVVKKFAFSKSVLTWSSTD | 13 |

| | | | |
|---|---|---|---|
| | | GNPSSATLSFVVLTDDDGQPLPANSYIGPQFTGFYWTSG AKPAAANTLGRNGAFPSGGGGGSGGGGGSSSQGADIST WVDSYQTYVVTTAGSWKDEDILKIDDDTAHTITYGPLKIVK YSLSNDTVSWSATDGNPFNAVIFFKVNKPTKANPTAGNQF VGKKWLPSDPAPAAVNWTGLIGSTADPKGTAAANATASM WKSIGINLGVAVSAMVLGTAVIKAIGAAWDKGSAAWKAAK AAADKAKKDAEAAEKDSAVDDEKFADEEPPDLEELPIPDA DPLVDVTDVDVTDVDVTDVDVTDVDVTDVDVTDVDVTDV DVTDVDVTDVDVVDVLDVVVI | |
| LepM1 | WP_061716904.1 | MDLLSKLEQAVGNEYLSNQNGLTIKINQEKQRQADSNKIV SLQPDSNKIVNFKKINA | 15 |
| LepM2 | WP_072355894.1 | MSILIKFESLAKQAIFSSNKDIILQSIGLLDTIKAKNNQKIRMT LSNNSEQFADMSRVVR | 15 |
| ParM | WP_007623905.1 | MMTKSLVDFLEELDSNAELKEAYLKDPVATATSYGLADAD VKIIKNKDWDTVKSKFENLNKDIKVHSY | 15 |
| PruM | WP_138552900.1 | MTVMKPKSIEIEYEIEQTAALEATSILSDLPNVVARKLTVDE VVAHTDKETHSDVDVDVDIDIDIDVDAIAVVDVDIDIDVDVV VDVTTHTDKD | 15 |
| SspM | WP_031073184.1 | MPAAVVDFMEELVTQPRRQHAYRRSAEAYVADSALTASE REAVVSGDVDRMRAVLAEHSGVKEECHAVLVVIIFDPDEV PSGA | 16 |
| SamM1 | WP_011758902.1 | MNQAVEILAKLGSNTAVSLDALPAHIAAMIEAKDIQALKAEL DVTPDVFCVFFPAEDEPEKEQQENEEEDEAPKAVNA | This work |
| SonM | WP_011071665.1 | MSGLSDFFTQLGQDAQLMEDYKQNPEAVMRAHGLTDEQI NAVMTGDMEKLKTLSGDSSYQSYLVISHGNGD | 16 |
| SahM | WP_037458248.1 | MSALSDFFTQLGQNAQLMEDYKQDPEAVMRAHGLTDEEI TAVMTGDMEKLKNLSGDESYQSYLVISHGNGD | 15 |
| SanM | WP_102440610.1 | MSNLTEFFTQLSSDAKLLETYKKDPRGVMQANGLSDKDID AVMSGDMAQVKSLVGDVELKSILLVHHLS | 15 |
| RceM | WP_012568691.1 | MTTIVPTELDQPDVIELSGGELDVAELSGGELDVAELFGG ELDVAELSGGELDVAELSGGELDVAELSGGELDVAELSG GELDVAELSGGELDVAELSGGELDVAELSGGELDVAEIGII NTFDL | 15,17 |
| SliM | ADB39711.1 | MTATNFSGNYMSFVLGVDGNTWGPGPVVAVNTTSQTLFI DGVQVNNPTFTSTSVMWMASSNNPSSGSIKFYSSAPDTG GFIGTYVEGTAPLPSSNNFSGVSSAAPDDLSVWNGTYNT FILNGSKWTQDSTLIVAAPNISYNGKDISNYIYMGTKSTQN LDQLSWFVAGGNAQNAIVEFFKDSSGYLTFGGTQWVSGD APAANNFIGTTKATPEPPTITAVVAIVINSSTTEVAEVTEVT EVTEVTEVTEVVEVVEVVEVVEVIAAAEEAEVKGQQQQSK ISTAQQDQEAAYKLSRKSNG | 17 |
| AinM | WP_083827882.1 | MANTRLLEGGTCQDLEDDFDEQSEEFDNASDDASLNNPE ELDIGDDVEIPDDPNVDVDVDADVEADIETEVDVDVDVDV TLEMFLEEDLERSEPGIDFVTGADGLDGDERP | 17 |
| PmoM | WP_051555776.1 | MATYPVQITNNTNLALDIYTTLNKNPATDPPSTNPADYTAV YTLQGSVGANQSTTLPLTESLARLVIVRQSDQFPLLVQVA NALLPDSEQVQVGNQDVTTANSGWAFYQSFISQPFTPTA LEFSELVSETPANALNDKAASFFAANGYPGVSFALFSALG YWANNQLYAYPGTYYCYEPPSGNSMGFILPTTSVGTLTIA | 17 |

| | | | |
|---|---|---|---|
| | | GGKANYSPSTGGSTALQFQYGQLTSPGADDKHGFNTTG FIRDLTWEGKPDVITWAFVGTYDGQQFIAQSYQNPQLPW YAVAYDMAYGALFTVQLAMTLDAAINLLGTVANGMQWLA QNTGKLISRIQDSLNSTGDTAGAGSGVGDAADPVNVDVDI DVDIDVDVDVDIDIDIDVDVDVDIDIDVDVDFIAVVDVDVDV DIDVDIDVVTDTETDIDIDVDVDIDTDVNVEPGALMKVVNG VGNWIMTKALPTLIEGAVIYVAFQSVGAIFQAWKNQDEKDI ENLQPRQSTGVGVLVNYMLQDDKPVAARWQTFSQYVAE VQGDPKTVGVTISTLLQTGNTKADNDAANWRWSSDDEN QVVASMAPYTGDQACKAFVILGNATYQGKPLPVKVGASV AMKYLAAQGA | |
| SurM1 | WP_165337934.1 | MATSALTNLLTEISEDPARLTEWLEDPDSYMNRAGLTEEE TRALRSASGGAIRHSIINSVTLEADRSELMAEILFKAWSDD DFREQLRTDPRAVLERLFTRRLPGIEFQVHEETPDVRHIVI PWLPPDVEEETRKNSRKLREIAEAGGPRTLLRPGNDEIPV VEAQGSESAVVPPPDTSNLQVVTVTDDIIIVDAIFAQEE | 17 |
| SurM2 | WP_165337936.1 | See SurM1 | 17 |
| LulM | WP_149193073.1 | MSNLLDLMRKLSGDAALAAEYEKDPDAVTRRAGLSDDER KAMLGKDYAAIKRLTGLADGKFATNHIIQAYDE | 15 |
| SkoAM | WP_018611879.1 | MENVVSSNTPNKKTTNSDFNYPFTDFIISLADSANFKNYLN NDEQYMTDVGLSEEQKLSVKAKEFYRVRGFLAREMANDS IHARLLNEFIESFNKYSKNDEFTHLTADVDVDLAPNHDVDH DVDHGAEAESHVLQKNYYNYLFKNWQRLHTNEVIFIGTGI KGANHISAEADAYIANASKVLYCVADLAVERKILLLNQNSE DLYIYYGDNKPRRETYEQMVDRILEVIKTEKLVCVVFYGHP GIFVWPSYQAIEKARGMGYKASMLPAISSLDCLFADIGFDP SRYSCQIVEATDLLIRSRNLDVSASVIIFQIGLVGDLGYSSK GFDGRNVPILADYLSSFYGDDYEVIIYEAAQYPIFEPKIQLIK INELSKAKLTGISTMYIPPKVSRATNQEMISKLGIKQSANST QKTH | This work |
| FpeAM | WP_083990299.1 | MPDTPLTDFLASLADPVKLELFTENPVAYLEKAGLSQELTN LILRGHSGAIRIEAVRELERAGLSPVVSDKFNPADQVSQQP MSISMNSYTSSTYTANSSTSTTFTSNTTDFVTTTSDTTTTT THNKFPGDFTTIDIVEEAIRYFEEGRSRKAGQLVIVGSGIRA IADLTFDAEAQIRTAQKVLYCVADPVIERRLHLLNSTAESLY GLYGNNKPRIETYHAMVDALLAPVRAGLRVCGVFYGHPG SFAWPTHQAIRVARRDGFKAEMHASVSADASLFADLGIDP SQPGCHSLEATEFLIHRRVPDTSSHLLVWQAECVGDPGF NFAGYKRHNFDILINQLQKFYPADHPVFIYEAASLPQGRPK IIKSTLDTIRKEDLTGISTLYLPPSIFNEIDQEMCLKLGL | This work |

**Table S2: Features and weights used for borosin precursor peptide scoring.** Scoring criteria and weights used by the RODEO borosin module to identify borosin precursor peptides. Custom Hidden Markov Models (HMMs) (Supplemental Dataset 1) are identified using HMMER[1], while motifs are identified using the MEME suite[2].

| | Feature | Weight |
|---|---|---|
| Heuristic scoring | No local* matches for the BorosinMT HMM | -5 |
| | No local* matches for a Borosin Binding Domain (BBD) HMM | -2 |
| | Local* ORF matches the BorosinMT HMM | +1 |
| | Local* ORF matches the NMT_1 HMM | +1 |
| | Local* ORF matches the NMT_2 HMM | +1 |
| | Local* ORF matches the PF03819 (MazG) or PF12643 (MazG-like) HMM | -4 |
| | Best BorosinMT HMM match has an e-value ≤ $10^{-40}$ | +1 |
| | Best BorosinMT HMM match has an e-value ≤ $10^{-30}$ | +1 |
| | Best BorosinMT HMM match has an e-value ≤ $10^{-25}$ | -2 |
| | Best BorosinMT HMM match has an e-value ≤ $10^{-15}$ | -3 |
| | Best BorosinMT HMM match has e-value ≤ $10^{-35}$ for PF00590 | -3 |
| | Best BorosinMT HMM match has e-value ≤ $10^{-25}$ for PF00590 | -2 |
| | Best BorosinMT HMM match has e-value ≤ $10^{-35}$ for PF03819 | -5 |
| | Best BorosinMT HMM match has e-value ≤ $10^{-25}$ for PF03819 | -3 |
| | Best BorosinMT HMM match has e-value ≤ $10^{-15}$ for PF03819 | -1 |
| | Local* ORF matches the PF07746 (LigA) or a BBD HMM | +2 |
| | ORF is encoded < 100 nucleotides from a methyltransferase | +1 |
| | ORF is encoded < 200 nucleotides from a methyltransferase | +1 |
| | ORF is encoded < 1,000 nucleotides from a methyltransferase | +1 |
| | ORF is encoded > 3,000 nucleotides from a methyltransferase | -2 |
| | ORF matches a BBD HMM | +3 |
| | ORF matches a Legionellales HMM | +4 |
| | ORF matches the Boro_6_8 HMM | +4 |
| | ORF matches both the Boro_6_8 and a methyltransferase HMM | -10 |
| | Core region has >10% Asp | +1 |
| | Core region has >10% Glu | +1 |
| | Core region has >10% Ile | +1 |
| | Core region has >10% Val | +1 |
| | Core region has >10% Thr | +1 |
| Motifs | No MEME motifs | -3 |
| | 3 or more MEME motifs | +1 |
| | 5 or more MEME motifs | +1 |
| | 7 or more MEME motifs | +1 |
| SVM | SVM classifies as valid | +10 |

*Within gene window called by RODEO, default is 8 open-reading frames (ORFs) in either direction from query protein

**Final positive cut-off score:** 16

**Table S3: Borosin module support vector machine parameters.** A support vector machine (SVM) was trained using 414 borosin precursor peptides with a 5-fold cross validation. Features were identified from characterized borosin BGCs.

| Heuristic | Type |
|---|---|
| ORF matches the PF00590 (TP methylase) or a borosin methyltransferase HMM | Boolean |
| ORF matches a Borosin Binding Domain (BBD) HMM | Boolean |
| ORF length | Integer |
| Length of the best scoring BorosinMT HMM match | Integer |
| ORF is the best scoring BorosinMT HMM match | Boolean |
| Length of the best scoring BBD HMM match | Integer |
| ORF is the best scoring BBD HMM match | Boolean |
| Multiple local* BorosinMT HMM matches | Boolean |
| Multiple local* BBD HMM matches | Boolean |
| Local* ORF matches both a methyltransferase and a BBD HMM | Boolean |
| Local* ORF matches both a methyltransferase and a GGDEF HMM | Boolean |
| Local* ORF matches the PF00590 HMM | Boolean |
| Local* ORF matches the BorosinMT HMM | Boolean |
| Local* ORF matches the NMT_1 HMM | Boolean |
| Local* ORF matches the NMT_2 HMM | Boolean |
| Local* ORF matches the PF03819 (MazG) or PF12643 (MazG-like) HMM | Boolean |
| E-value for the best scoring BorosinMT HMM match | Float |
| Best BorosinMT HMM match has an e-value ≤ $10^{-40}$ | Boolean |
| Best BorosinMT HMM match has an e-value ≤ $10^{-30}$ | Boolean |
| Best BorosinMT HMM match has an e-value ≤ $10^{-25}$ | Boolean |
| Best BorosinMT HMM match has an e-value ≤ $10^{-15}$ | Boolean |
| E-value of the best BorosinMT match for the PF00590 HMM | Float |
| Best BorosinMT HMM match has e-value ≤ $10^{-35}$ for PF00590 | Boolean |
| Best BorosinMT HMM match has e-value ≤ $10^{-25}$ for PF00590 | Boolean |
| E-value of the best BorosinMT match for the PF03819 HMM | Float |
| Best BorosinMT HMM match has e-value ≤ $10^{-35}$ for PF03819 | Boolean |
| Best BorosinMT HMM match has e-value ≤ $10^{-25}$ for PF03819 | Boolean |
| Best BorosinMT HMM match has e-value ≤ $10^{-15}$ for PF03819 | Boolean |
| Best BorosinMT HMM match has e-value ≤ $10^{-5}$ for PF03819 | Boolean |
| Local* ORF matches the PF07746 (LigA) HMM | Boolean |
| Local* ORF matches a BBD HMM | Boolean |
| E-value for the best PF07746 HMM match | Float |
| E-value for the best BBD_A HMM match | Float |
| E-value for the best BBD_B HMM match | Float |
| E-value for the best BBD_C HMM match | Float |
| E-value for the best BBD_old HMM match | Float |
| ORF is encoded < 100 nucleotides from a methyltransferase | Boolean |

| | |
|---|---|
| ORF is encoded < 200 nucleotides from a methyltransferase | Boolean |
| ORF is encoded < 1,000 nucleotides from a methyltransferase | Boolean |
| ORF is encoded > 3,000 nucleotides from a methyltransferase | Boolean |
| ORF and borosin methyltransferase are encoded on the same strand | Boolean |
| Local* matches for both a BBD and the PF00590 HMMs | Boolean |
| Local* GGDEF | Boolean |
| Local* acetyltransferase | Boolean |
| Local* peptidase | Boolean |
| Local* ORF matches a Legionellales HMM | Boolean |
| Local* ORF matches a Boro_6_8 HMM | Boolean |
| ORF matches the PF00590 (TP methylase) HMM | Boolean |
| ORF matches the BorosinMT HMM | Boolean |
| ORF matches the NMT_1 HMM | Boolean |
| ORF matches the NMT_2 HMM | Boolean |
| ORF matches the PF07746 (LigA) HMM | Boolean |
| ORF matches the BBD_A HMM | Boolean |
| ORF matches the BBD_B HMM | Boolean |
| ORF matches the BBD_C HMM | Boolean |
| ORF matches the BBD_old HMM | Boolean |
| ORF matches a Legionellales HMM | Boolean |
| ORF matches the Boro_6_8 HMM | Boolean |
| ORF matches both a borosin methyltransferase and a BBD HMM | Boolean |
| ORF matches both a Legionelalles and a BBD HMM | Boolean |
| ORF matches both the Boro_6_8 and a BBD HMM | Boolean |
| ORF matches a Legionelalles and a borosin methyltransferase HMM | Boolean |
| ORF matches the Boro_6_8 and a borosin methyltransferase HMM | Boolean |
| ORF is >900 AA and matches a BBD HMM | Boolean |
| ORF is >700 AA and matches a BBD HMM | Boolean |
| ORF is <400 AA and matches a BBD HMM | Boolean |
| ORF is <250 AA and matches a BBD HMM | Boolean |
| ORF is <100 AA and matches a BBD HMM | Boolean |
| ORF is <400 AA and matches the Boro_6_8 HMM | Boolean |
| ORF is <100 AA and matches a Legionellales HMM | Boolean |
| ORF contains MEME motifs (1-27) | Boolean |
| Total MEME motifs detected in ORF | Integer |
| Occurrences of each MEME motif detected in ORF | Integer |
| Minimum distance between ORF and borosin methyltransferase | Integer |
| Net charge of predicted “core region” of ORF | Float |
| Net charge of ORF | Float |
| Total charge of predicted “core region” of ORF | Float |
| Total charge of ORF | Float |

*Within gene window called by RODEO, default is 8 open-reading frames (ORFs) in either direction from query protein.

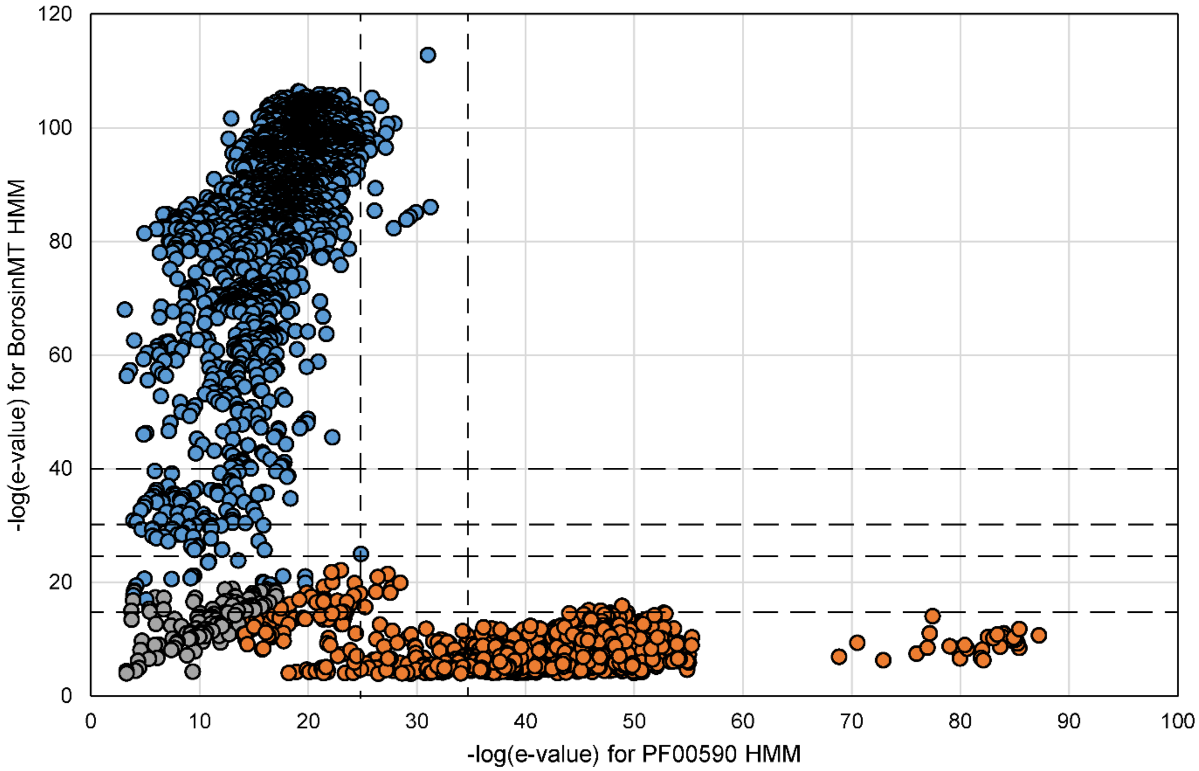


**Figure S1: Expectation value of PF00590 (TP methylase) vs BorosinMT HMMs.** A protein BLAST search of characterized borosins yielded borosin and non-borosin methyltransferases. Using hmmscan[1] we calculated the expectation value (e-value) of the BorosinMT (Supplementary dataset 1) and PF00590 HMMs for 4,697 of the retrieved sequences and then plotted the data on a negative log scale. Borosins were assigned based on the presence of precursor and methyltransferase HMMs; sequences which scored poorly on these HMMs were manually inspected, with ambiguous BGCs left undetermined. Heuristic weights were added for e-value cutoffs for the BorosinMT and PF00590 (TP methylase) HMM based on analyzing the trends above (Table S1).

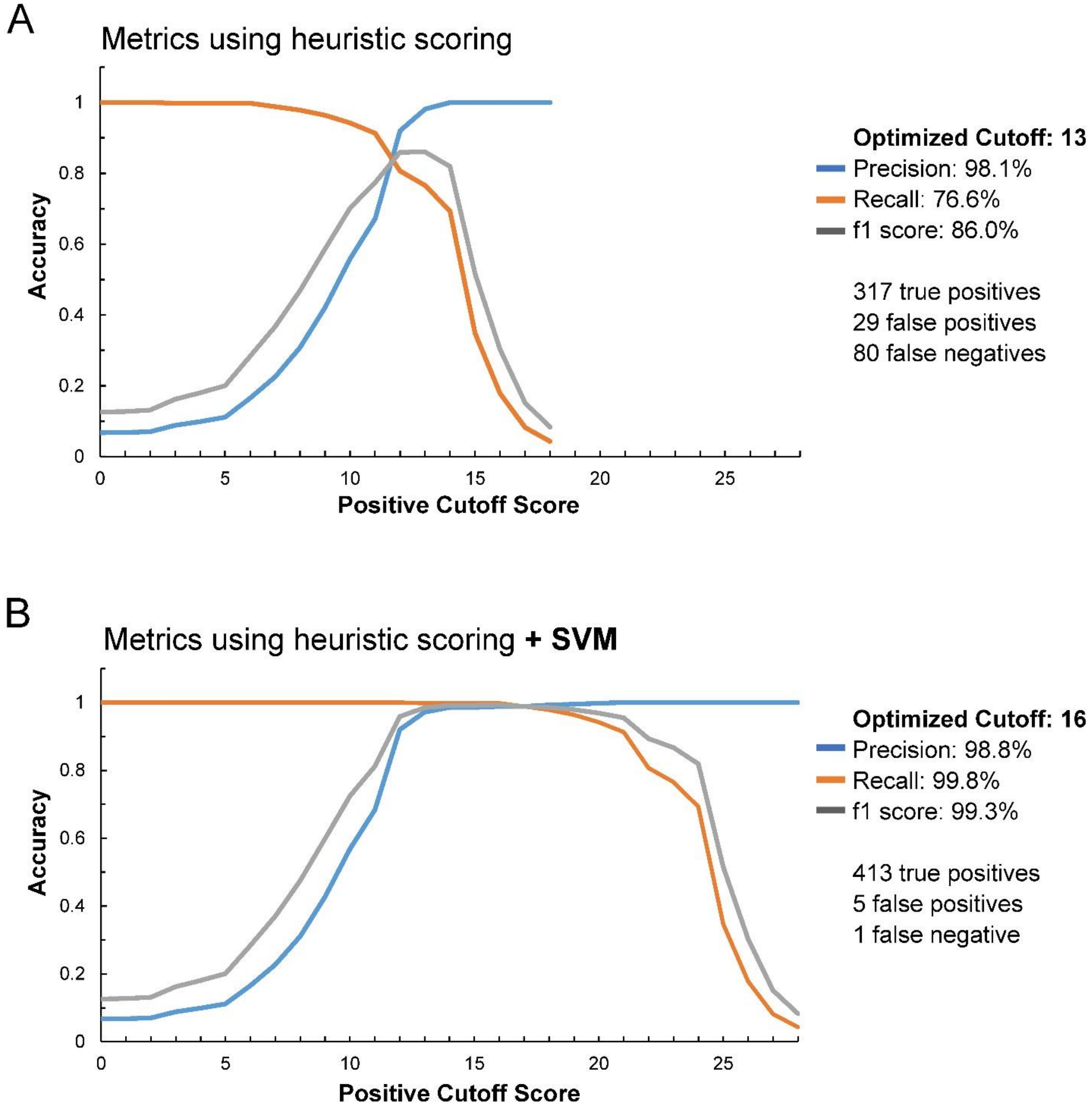


**Figure S2: Borosin module scoring accuracy.** Scoring accuracy of the borosin module on a test set of 414 high confidence predicted borosins **A.** with only the heuristic and motif scoring weights and **B.** with the addition of the SVM scoring weights. Precision (the ability to minimize false positives), Recall (the ability to minimize false negatives), and the f1 score (used to optimize the balance between precision and recall) were calculated using equations S1-S3.

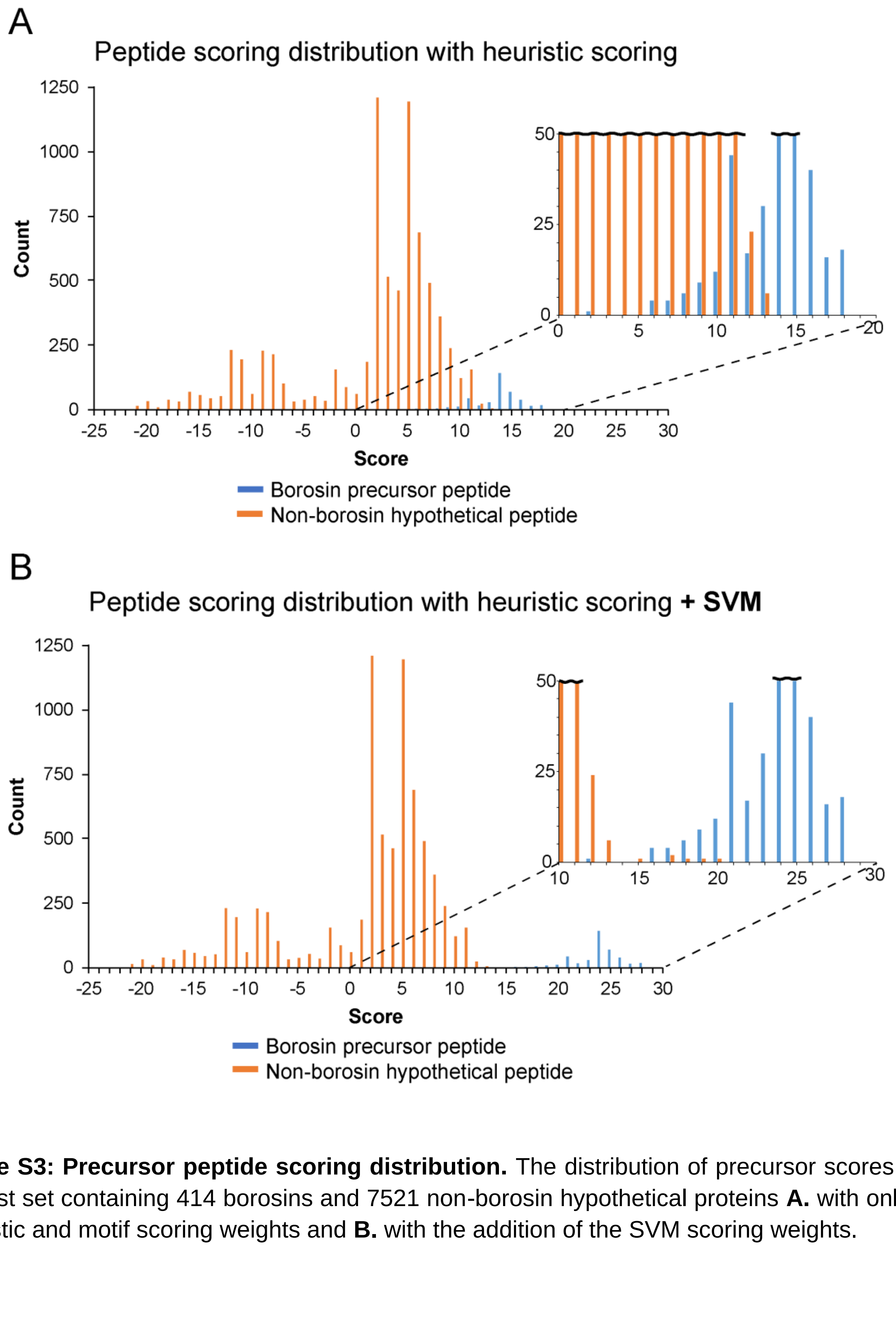


**Figure S3: Precursor peptide scoring distribution.** The distribution of precursor scores from the test set containing 414 borosins and 7521 non-borosin hypothetical proteins **A.** with only the heuristic and motif scoring weights and **B.** with the addition of the SVM scoring weights.

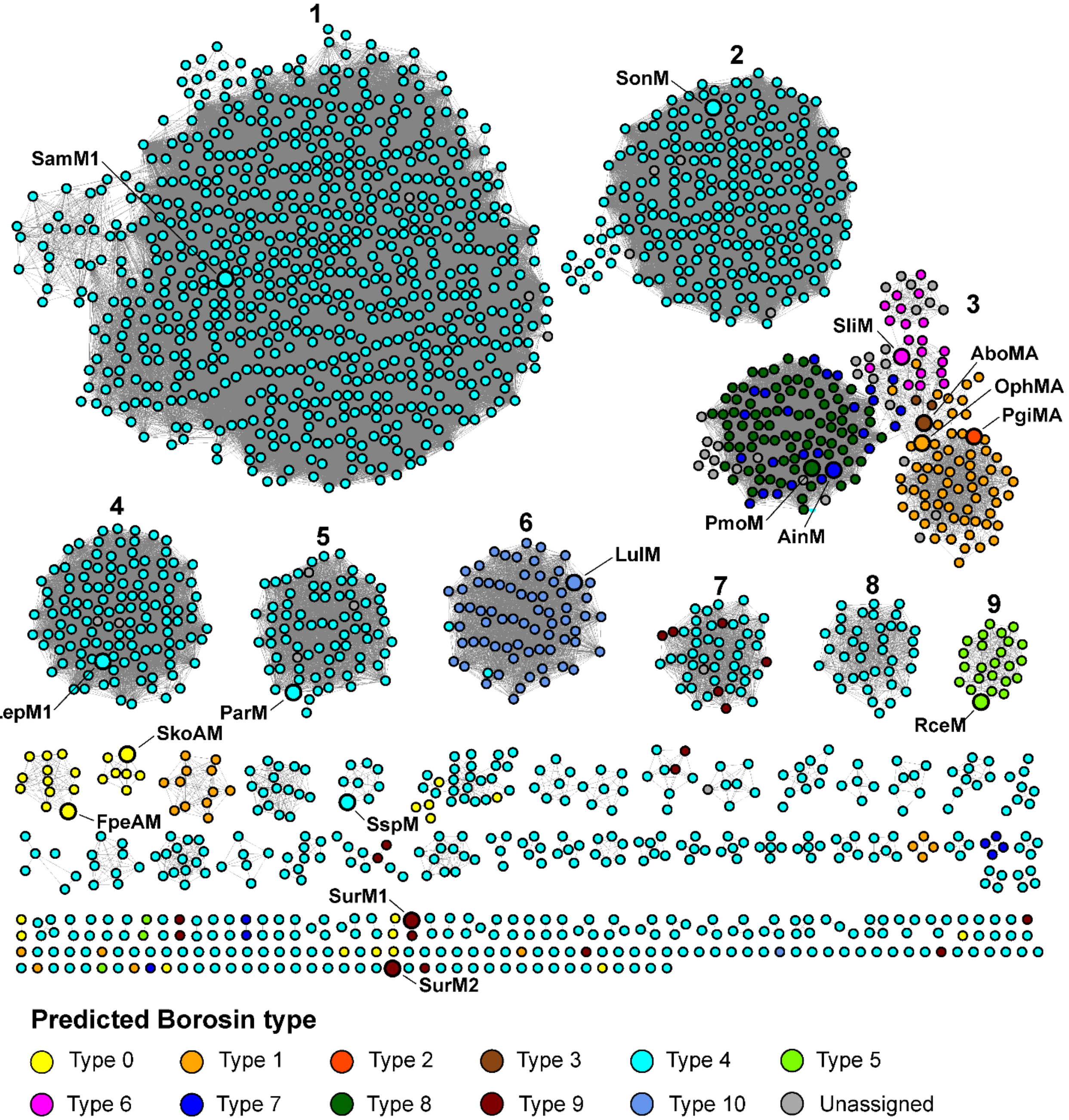


**Figure S4: SSN of borosin methyltransferases colored by predicted type.** Borosin methyltransferases with a RODEO-identified precursor peptide were used to generate this SSN (*n* = 2,124, alignment score = 90). Sequences for known borosin methyltransferase that are absent from the NCBI protein database were manually added (OphMA, PgiMA and AboMA) (Table S3).

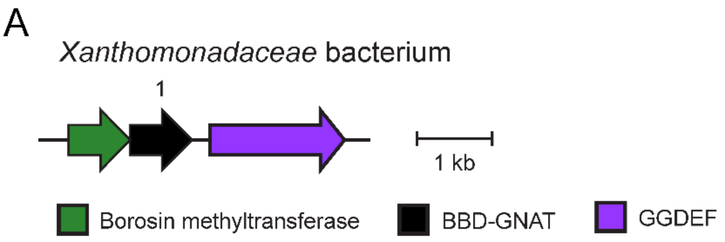


```
1:   MIVTPRLRLRALALRDMALFRALYCDPGTMRHIGKPLSRAVAVASLRATVAATRKPAGLRFFVIAECQS    69
     RRGVGLCSLRPAAWDKRGIELGIMLVHTARRHGYAREAFAALINYAFRALPIDTIWVQYRRANAGMARL   138
     CDILGFEGNFLPRSGTSVRRCVRIRRDRNGAVFSNQPTRGITMSNIIGFLEQAGSNAAMRHASREALLR   207
     TMQDKEIENVLRQPHRSLLDDLTGARETMYCKNTAIAPPKQKAPAKKKPAKKTPAKKPAKKAPAKRKR    275
```

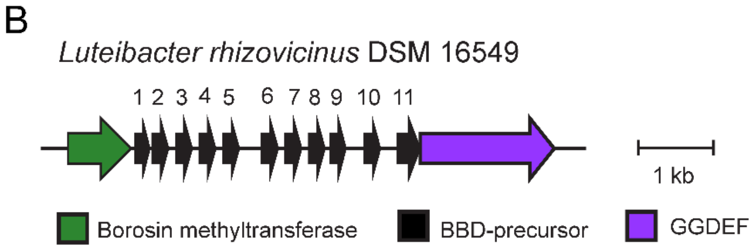


```
1:   MLDTIDILETIGSDASLRHAHTDALSAALEQLHASPALTSAAVTGDSSELFAELGVKRMFLPQVSQV            67
2:   MLDTLEMLEAIGSDASLRYASTAELTTVLERGKASEAFTAAVASGDATRLDEEFGHRKMYQPQASQTFLSSELG     74
3:   MLDTLEMLEAIGSDASLRYASTVELTTVLERGKASEAFTAAVVSGDASRLDEEFGYRKMYQPQATQSFCSSELS     74
4:   MLDTLEMLEAIGSDASLRYASTVELTTVLERGKASEAFTAAVASGDASRLDEEFGYRKMYQPQATQSFCSSELS     74
5:   MLDTIEMLEAIGSDASLRYASTVELTNVLEKAQASAAFTAAVASGDTSRLDQEFDGRKMYAPQAVQTFLASDPD     74
6:   MLDTLEMLEAIGSDASLRYASTVELTTVLEAGRASDALTAAVASGDSSHLAREFGNIKNFSPQVTQQIFVSSDID    75
7:   MLDTIEMLEAIGSDASLRYASTVELTNVLEKAQASAAFTAAVASGDTSRLDQEFDGRKMYAPQAVQTFLASDPD     74
8:   MLDTLEMLEAIGSDASLRYAPTVDLASALENAKAIEALTEAVVSGDITRLRQDFGGSQMHAPQISQSFFDRP       72
9:   MLDTIEMLEVIGSDASLRYASAVELTNALEQAEATEALTAAVASGDSTHLLRELGSKRMYLPQVIQSYS          69
10:  MLDTIEILEAIGRDASLRHASKAKLADVLSNAHASEALASAVKYGDASHLAREFGTAANFVPQAIQTFFAPG       72
11:  MSDTIELLEAIGRDASLRHASSEHLANVLQESQASAGLTAAVASGDSSLLSEELGHKHMDLPQIVQVPGHDEDDM    75
     ESGDGDSSDKPSKPDDGKQPSEESTK                                                    101
```

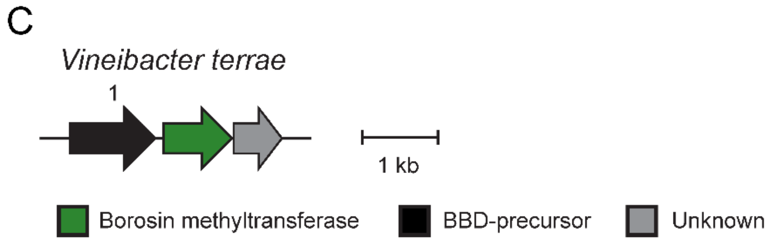


```
1:   MATRKRGGTGPVAEPIIPPGEATPPDEFRSFMVALATDPAKLGAFIKDPDGTMQAAGISDVDQVILKSG    69
     HPWAIHGRLSGQKYTFTPPSPATPVTVLVVDMTKPPGADQAAAADEPAVRAQPAFTTPPSQGSAAMFPN   138
     IPQQIFPQVHPQIFPQVHPQIWPQVHPQVVVHPQVWPQVHPQLVVHPQVWPQVHPQLVVHPQVWPQVHP   207
     QLVVHPQIWPQVHPQLVVHPQIWPQIHPQLVVHPQVHPQLVQPNAALPQFPQLVLHPQVHPQIWPQVHP   276
     QVVVHPQIWPQVHPQLVVHPQIWPQVHPQVVVHPQIWPQVHPQVVVHPQIWPQVHPQVVVHPQIWPQVH   345
     PQLVVHPQIWPQVHPQLVTPPPATPGTPG                                            374
```

**Figure S5: Unusual borosin BGC architectures. A.** A borosin BGC with a BBD-fused GNAT acetyltransferase (MBU6199443.1) identified as a precursor peptide by RODEO. **B.** A borosin methyltransferase (WP_046969431.1) with 11 nearby BBDs. C. A borosin BGC with a BBD-fused precursor (WP_178133946.1) 374 amino acids long containing repeating motifs.

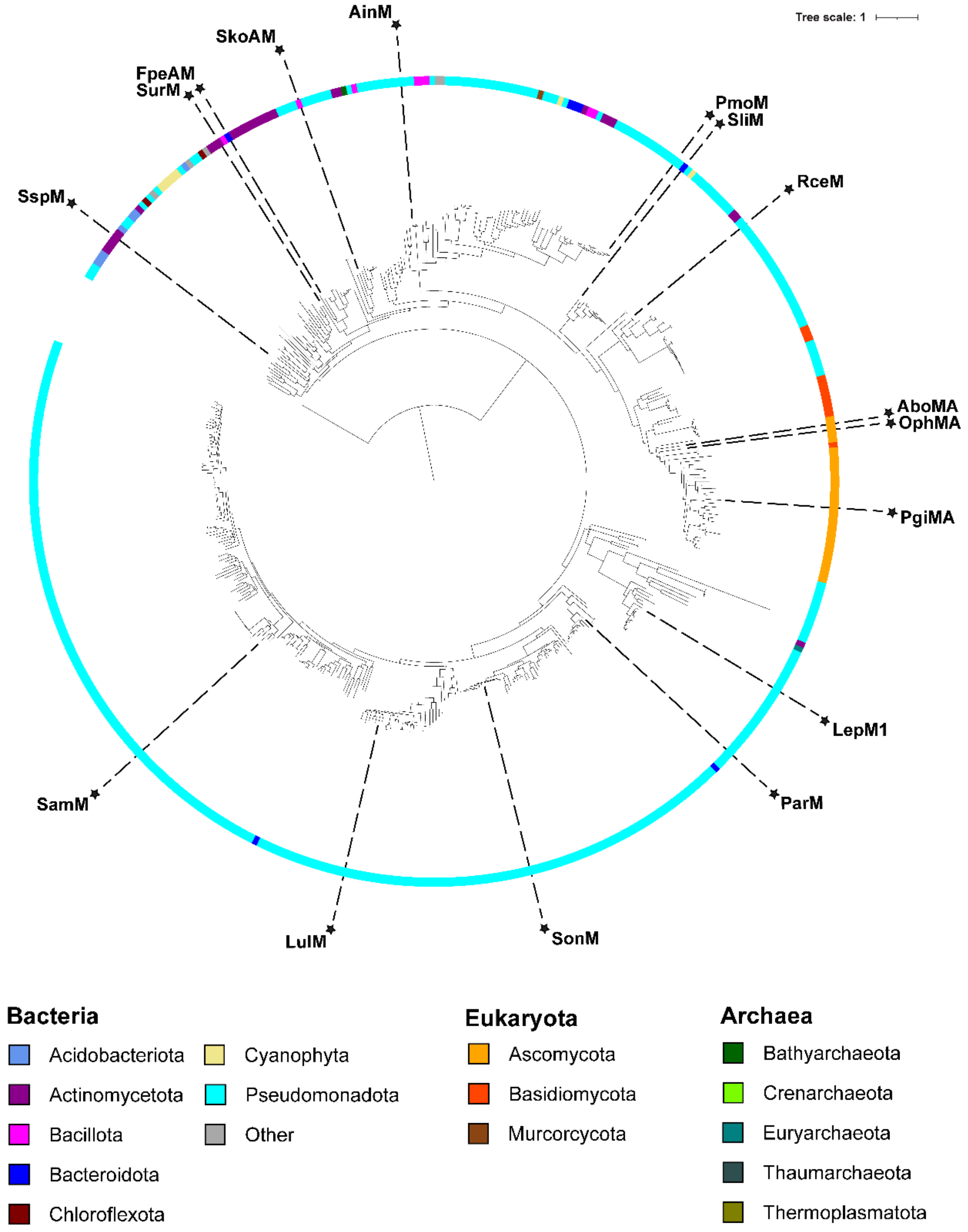


**Figure S6: Phylogenetic distribution of diversity-maximized subset of RODEO curated borosin methyltransferases.** A maximum likelihood tree of a diversity-maximized sample of borosin methyltransferases. The tree was rooted using the non-borosin TP methylase MCB1244216.1.

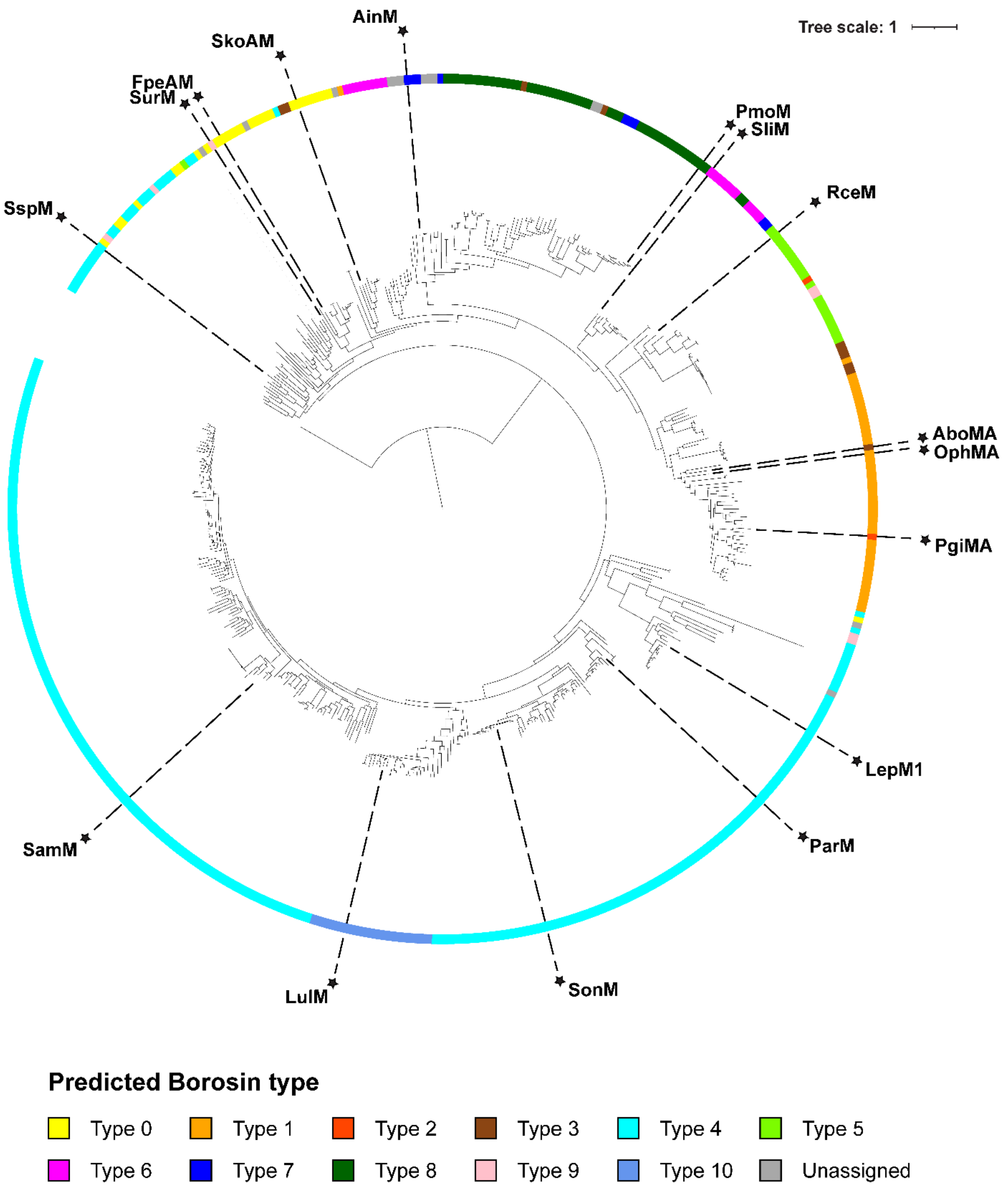


**Figure S7: Type distribution of diversity-maximized subset of RODEO curated borosin methyltransferases.** A maximum likelihood tree of a diversity-maximized sample of borosin methyltransferases. The tree was rooted using the non-borosin TP methylase MCB1244216.1.

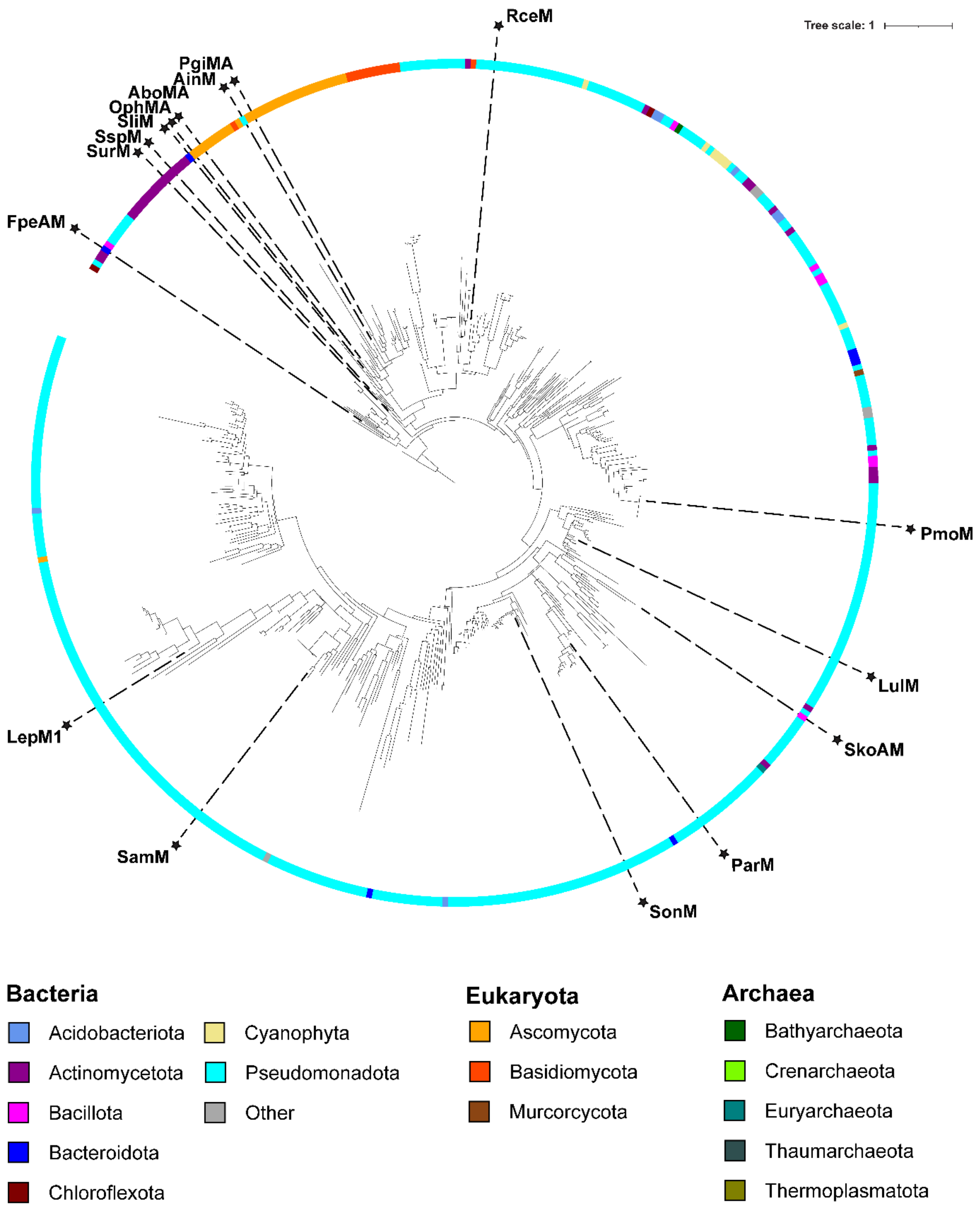


**Figure S8: Phylogenetic distribution of excised BBDs.** A maximum likelihood tree of the excised BBD domain from the diversity-maximized sample of borosin methyltransferases or their cognate BBD-fused precursor peptide. The tree was rooted using a homologous LigA protein APV50782.1.

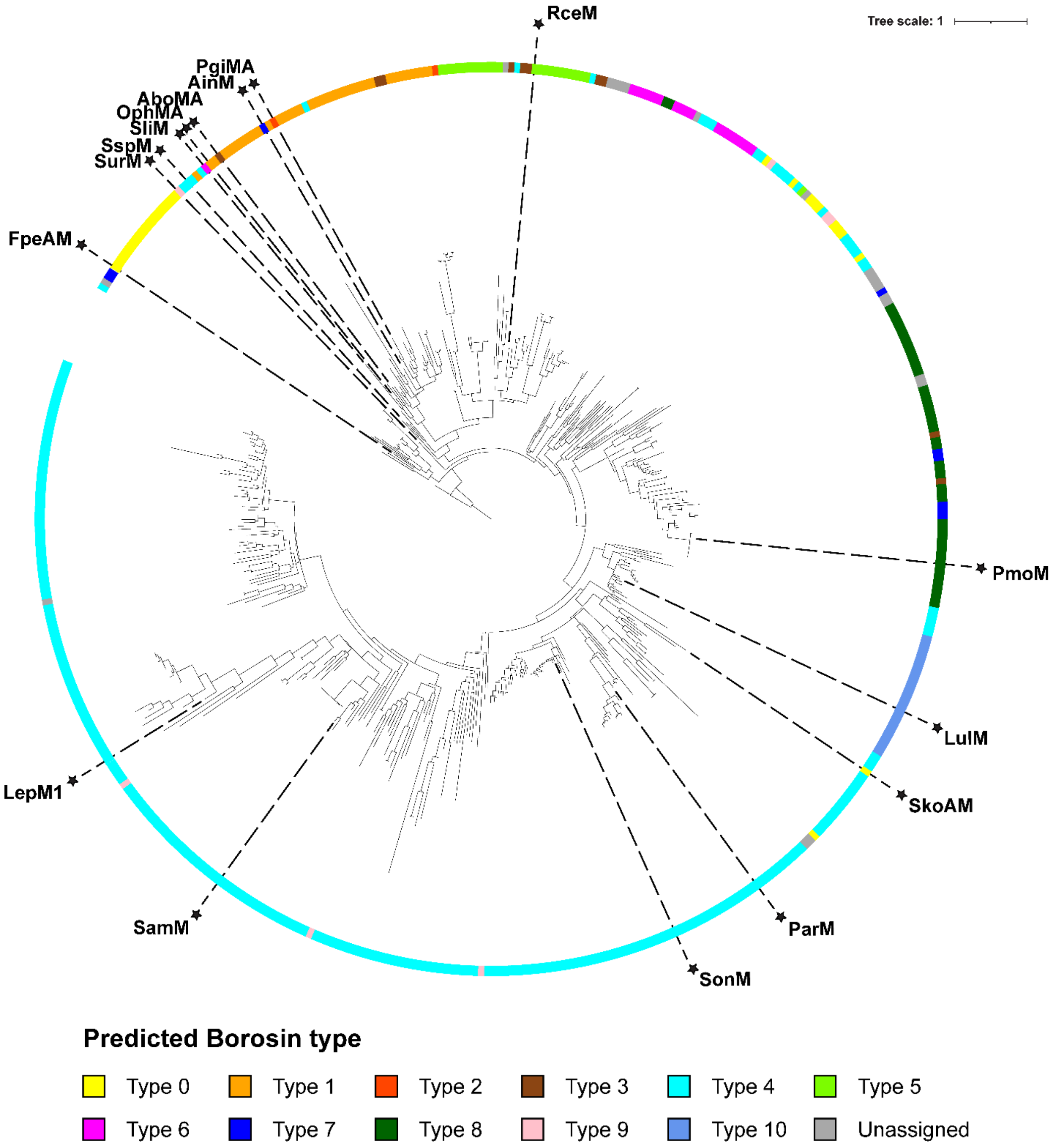


**Figure S9: Type distribution of excised BBDs.** A maximum likelihood tree of the excised BBD domain from the diversity-maximized sample of borosin methyltransferases or their cognate BBD-fused precursor peptide. The tree was rooted using a homologous LigA protein APV50782.1.

A

| | MCB1244216.1 | LulM | SurM1 | PmoM | AinM | SliM | RceM | LepM1 | SspM | ParM | SonM | SamM | AboMA | PgiMA1 | OphMA | FpeAM | SkoAM |
|---|---|---|---|---|---|---|---|---|---|---|---|---|---|---|---|---|---|
| SkoAM | 5.3 | 3.9 | 2.2 | 3.8 | 3.1 | 2.5 | 3.1 | 3.8 | 2.8 | 2.8 | 3.0 | 3.6 | 2.9 | 3.8 | 2.9 | 2.3 | 0.0 |
| FpeAM | 5.1 | 3.7 | 1.7 | 3.8 | 3.1 | 2.5 | 3.1 | 3.7 | 2.6 | 2.6 | 2.8 | 3.4 | 2.9 | 3.9 | 3.0 | 0.0 | |
| OphMA | 6.0 | 4.6 | 2.8 | 3.7 | 3.0 | 2.2 | 2.2 | 4.5 | 3.5 | 3.5 | 3.7 | 4.3 | 1.2 | 2.0 | 0.0 | | |
| PgiMA1 | 6.9 | 5.5 | 3.7 | 4.6 | 3.9 | 3.1 | 3.1 | 5.4 | 4.4 | 4.4 | 4.6 | 5.2 | 2.1 | 0.0 | | | |
| AboMA | 6.0 | 4.5 | 2.8 | 3.7 | 3.0 | 2.1 | 2.2 | 4.5 | 3.4 | 3.5 | 3.6 | 4.3 | 0.0 | | | | |
| SamM | 5.6 | 2.7 | 3.3 | 5.2 | 4.5 | 3.9 | 4.5 | 3.9 | 3.2 | 2.4 | 1.9 | 0.0 | | | | | |
| SonM | 5.0 | 2.2 | 2.7 | 4.5 | 3.8 | 3.2 | 3.8 | 3.3 | 2.6 | 1.7 | 0.0 | | | | | | |
| ParM | 4.8 | 2.7 | 2.5 | 4.4 | 3.7 | 3.1 | 3.7 | 3.1 | 2.5 | 0.0 | | | | | | | |
| SspM | 4.9 | 3.5 | 2.5 | 4.3 | 3.7 | 3.0 | 3.6 | 3.5 | 0.0 | | | | | | | | |
| LepM1 | 5.8 | 4.2 | 3.5 | 5.4 | 4.7 | 4.1 | 4.7 | 0.0 | | | | | | | | | |
| RceM | 6.1 | 4.7 | 3.0 | 3.9 | 3.2 | 2.3 | 0.0 | | | | | | | | | | |
| SliM | 5.6 | 4.1 | 2.4 | 3.3 | 2.6 | 0.0 | | | | | | | | | | | |
| AinM | 6.2 | 4.8 | 3.0 | 2.6 | 0.0 | | | | | | | | | | | | |
| PmoM | 6.9 | 5.4 | 3.7 | 0.0 | | | | | | | | | | | | | |
| SurM1 | 5.0 | 3.6 | 0.0 | | | | | | | | | | | | | | |
| LulM | 5.9 | 0.0 | | | | | | | | | | | | | | | |
| MCB1244216.1 | 0.0 | | | | | | | | | | | | | | | | |

B

| | APV50782.1 | LulM | SurM1 | PmoM | AinM | SliM | RceM | LepM1 | SspM | ParM | SonM | SamM | AboMA | PgiMA1 | OphMA | FpeAM | SkoAM |
|---|---|---|---|---|---|---|---|---|---|---|---|---|---|---|---|---|---|
| SkoAM | 3.2 | 2.3 | 2.8 | 3.4 | 3.7 | 4.0 | 3.7 | - | 3.1 | 2.6 | 1.9 | 2.0 | 2.0 | 4.1 | 2.0 | 2.0 | 0.0 |
| FpeAM | 1.6 | 1.8 | 1.3 | 2.6 | 2.2 | 2.6 | 2.2 | - | 1.7 | 2.7 | 2.0 | 3.4 | 2.0 | 2.0 | 2.0 | 0.0 | |
| OphMA | 2.1 | 2.1 | 1.7 | 2.9 | 1.2 | 1.0 | 1.8 | - | 1.7 | 3.0 | 2.3 | 3.7 | 2.0 | 2.0 | 0.0 | | |
| PgiMA1 | 2.8 | 2.8 | 2.4 | 3.6 | 1.1 | 2.3 | 2.6 | - | 2.5 | 3.7 | 3.0 | 4.4 | 1.8 | 0.0 | | | |
| AboMA | 2.3 | 2.3 | 1.9 | 3.1 | 1.4 | 1.5 | 2.0 | - | 1.9 | 3.2 | 2.5 | 3.8 | 0.0 | | | | |
| SamM | 3.5 | 2.5 | 3.1 | 3.7 | 4.0 | 4.3 | 3.9 | - | 3.4 | 3.3 | 2.5 | 0.0 | | | | | |
| SonM | 2.1 | 1.2 | 1.7 | 2.3 | 2.6 | 2.9 | 2.6 | - | 2.0 | 1.5 | 0.0 | | | | | | |
| ParM | 2.9 | 1.9 | 2.5 | 3.0 | 3.3 | 3.7 | 3.3 | - | 2.7 | 0.0 | | | | | | | |
| SspM | 1.8 | 1.8 | 1.4 | 2.6 | 2.0 | 2.4 | 2.0 | - | 0.0 | | | | | | | | |
| LepM1 | - | - | - | - | - | - | - | 0.0 | | | | | | | | | |
| RceM | 2.4 | 2.4 | 2.0 | 3.2 | 2.1 | 2.5 | 0.0 | | | | | | | | | | |
| SliM | 2.7 | 2.7 | 2.3 | 3.5 | 1.8 | 0.0 | | | | | | | | | | | |
| AinM | 2.4 | 2.4 | 2.0 | 3.2 | 0.0 | | | | | | | | | | | | |
| PmoM | 2.8 | 2.1 | 2.3 | 0.0 | | | | | | | | | | | | | |
| SurM1 | 1.4 | 1.5 | 0.0 | | | | | | | | | | | | | | |
| LulM | 2.0 | 0.0 | | | | | | | | | | | | | | | |
| APV50782.1 | 0.0 | | | | | | | | | | | | | | | | |

**Figure S10: Sum of branch distances from diversity-maximized tree**. A. The sum of the branch distances from the diversity-maximized tree between select borosin methyltransferases and B. the cognate excised BBDs. MCB1244316.1, a PF00590 TP methylase, was used for the methyltransferase outgroup, while APV50782.1, a LigA protein, was used for the BBD outgroup.

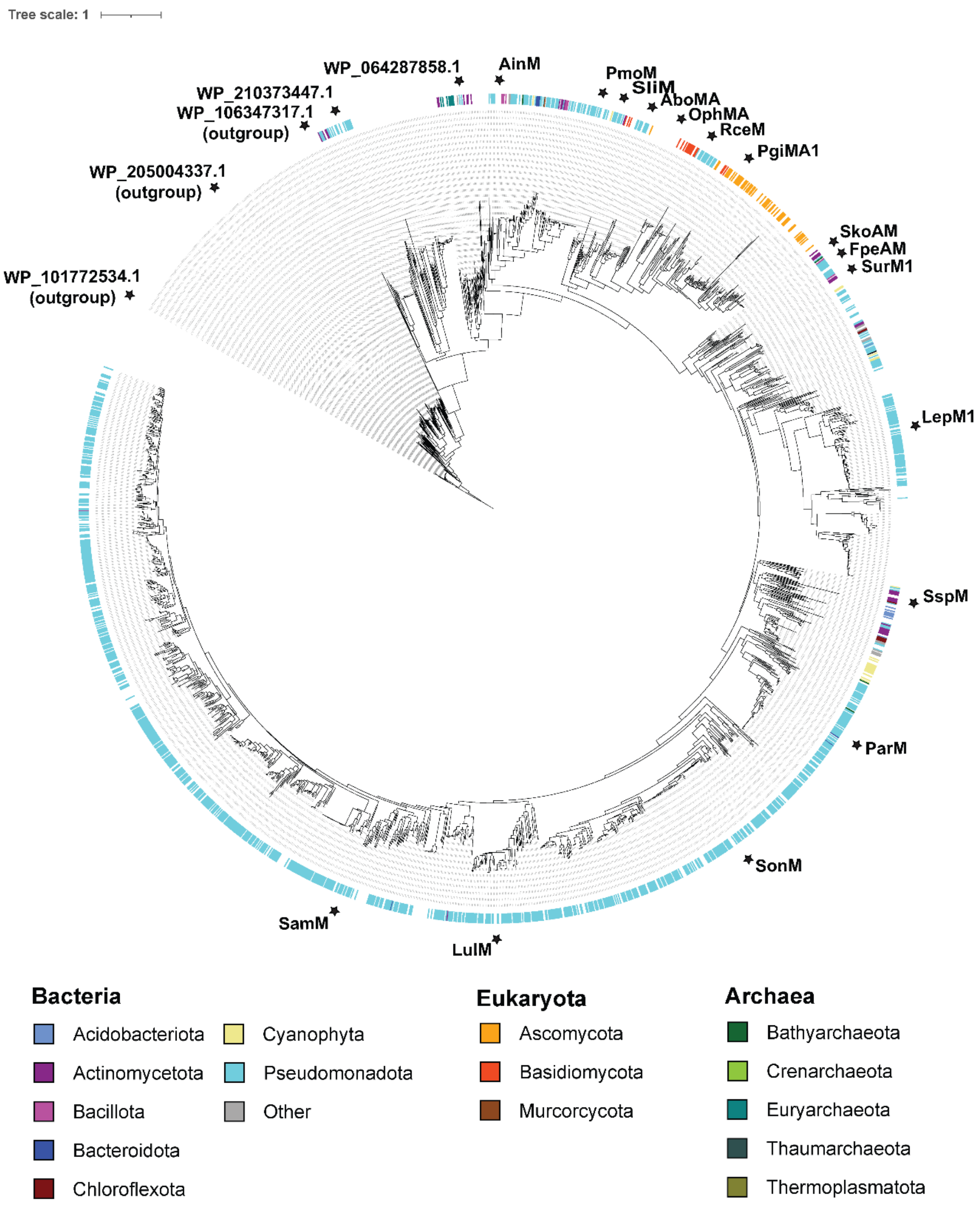


**Figure S11: Phylogenetic tree of SspM homologs.** A phylogenetic tree composed of the top 2375 homologs for SspM (WP_031073184.1), identified by a BLAST search of the NCBI non-redundant (nr) protein database. Members of the RODEO curated borosin dataset are colored by phylum, while gaps in the coloring scheme represent homologs without a RODEO identified precursor. WP_106347317.1, WP_205004337.1, and WP 101772534.1 are non-borosin PF00590 TP methylase outgroups, with WP 101772534.1 used to root the tree. WP_210373447.1 and WP 064287858.1 are members of RODEO predicted borosins groups with a BBD-precursor fusion (type IV) that are more closely related to the outgroups than any of the characterized borosins.

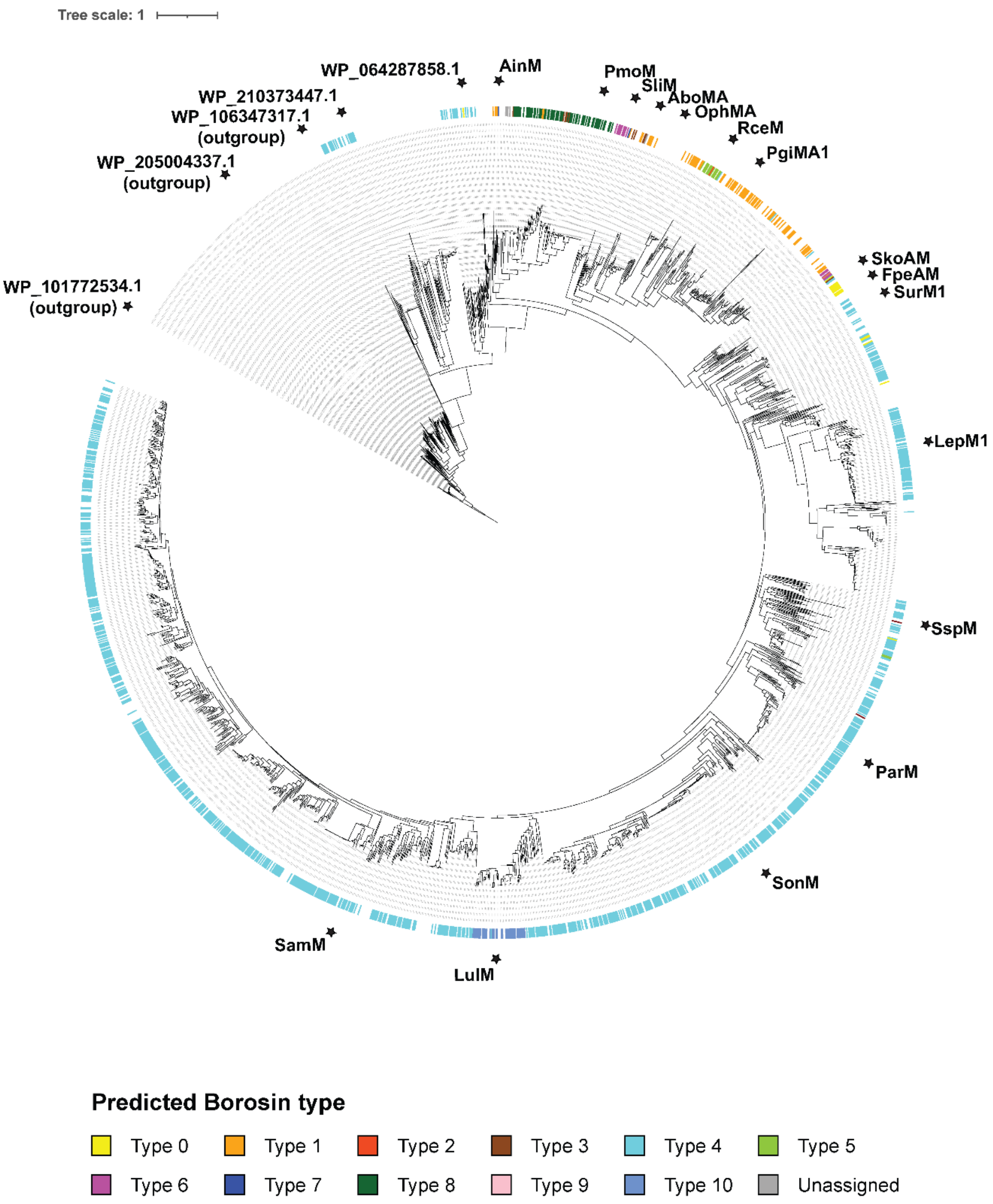


**Figure S12: Type distribution of SspM homologs.** A phylogenetic tree of 2,375 SspM (WP_031073184.1) homologs identified by a BLAST search of the NCBI non-redundant (nr) protein database. Members of the RODEO curated borosin dataset are colored by borosin type, while gaps in the coloring scheme represent homologs without a RODEO identified precursor. WP_106347317.1, WP_205004337.1, and WP 101772534.1 are non-borosin PF00590 TP methylase outgroups, with WP 101772534.1 used to root the tree. WP_210373447.1 and WP 064287858.1 are members of RODEO predicted borosins groups with a BBD-precursor fusion (type IV) that are more closely related to the outgroups than any of the characterized borosins.

| | WP_106347317.1 | WP_101772534.1 | WP_205004337.1 | WP_064287858.1 | WP_210373447.1 | LulM | ParM | SonM | SahM | SanM | SamM | SspM | LepM2 | LepM1 | SurM2 | SurM1 | AinM | PruM | PmoM | SliM | RceM | AboMA | PgiMA1 | PocMA | MroMA2 | MroMA1 | CeuMA | GjuMA | SveMA | DbiMA2 | DbiMA1 | OphMA | LedMA | CmiMA | CmaMA | SkoAM | FpeAM |
|---|---|---|---|---|---|---|---|---|---|---|---|---|---|---|---|---|---|---|---|---|---|---|---|---|---|---|---|---|---|---|---|---|---|---|---|---|---|
| FpeAM | 4.9 | 4.9 | 4.4 | 3.1 | 3.4 | 2.5 | 2.0 | 2.0 | 2.0 | 2.4 | 2.4 | 1.8 | 2.6 | 2.6 | 1.3 | 1.2 | 2.0 | 2.2 | 2.3 | 1.9 | 2.4 | 1.7 | 2.4 | 2.3 | 2.5 | 2.4 | 2.5 | 2.3 | 2.1 | 2.1 | 1.9 | 1.9 | 1.9 | 2.4 | 2.2 | 1.5 | 0.0 |
| SkoAM | 4.8 | 4.8 | 4.4 | 3.0 | 3.3 | 2.5 | 2.0 | 2.0 | 2.0 | 2.4 | 2.4 | 1.8 | 2.6 | 2.6 | 1.3 | 1.2 | 2.0 | 2.1 | 2.3 | 1.9 | 2.3 | 1.6 | 2.4 | 2.2 | 2.4 | 2.3 | 2.4 | 2.3 | 2.1 | 2.0 | 1.8 | 1.8 | 1.8 | 2.3 | 2.1 | 0.0 | |
| CmaMA | 5.0 | 5.0 | 4.5 | 3.2 | 3.5 | 3.2 | 2.8 | 2.8 | 2.8 | 3.1 | 3.1 | 2.6 | 3.3 | 3.3 | 2.1 | 2.0 | 2.1 | 2.3 | 2.4 | 1.9 | 1.8 | 1.0 | 1.9 | 1.7 | 1.9 | 1.9 | 2.0 | 1.8 | 1.6 | 1.4 | 1.2 | 1.3 | 1.3 | 0.4 | 0.0 | | |
| CmiMA | 5.2 | 5.1 | 4.7 | 3.3 | 3.7 | 3.4 | 2.9 | 2.9 | 2.9 | 3.3 | 3.3 | 2.7 | 3.5 | 3.5 | 2.3 | 2.1 | 2.3 | 2.4 | 2.6 | 2.1 | 2.0 | 1.1 | 2.0 | 1.9 | 2.1 | 2.0 | 2.1 | 1.9 | 1.7 | 1.6 | 1.4 | 1.4 | 1.4 | 0.0 | | | |
| LedMA | 4.7 | 4.7 | 4.3 | 2.9 | 3.2 | 2.9 | 2.5 | 2.5 | 2.5 | 2.8 | 2.9 | 2.3 | 3.0 | 3.0 | 1.8 | 1.7 | 1.8 | 2.0 | 2.2 | 1.6 | 1.5 | 0.8 | 1.6 | 1.4 | 1.6 | 1.6 | 1.7 | 1.5 | 1.3 | 0.5 | 0.2 | 0.2 | 0.0 | | | | |
| OphMA | 4.7 | 4.7 | 4.2 | 2.9 | 3.2 | 2.9 | 2.4 | 2.4 | 2.5 | 2.8 | 2.8 | 2.3 | 3.0 | 3.0 | 1.8 | 1.7 | 1.8 | 2.0 | 2.1 | 1.6 | 1.5 | 0.8 | 1.6 | 1.4 | 1.6 | 1.6 | 1.7 | 1.5 | 1.3 | 0.4 | 0.1 | 0.0 | | | | | |
| DbiMA1 | 4.7 | 4.6 | 4.2 | 2.8 | 3.2 | 2.9 | 2.4 | 2.4 | 2.4 | 2.8 | 2.8 | 2.2 | 3.0 | 3.0 | 1.8 | 1.6 | 1.8 | 1.9 | 2.1 | 1.6 | 1.5 | 0.7 | 1.6 | 1.4 | 1.6 | 1.5 | 1.6 | 1.4 | 1.2 | 0.4 | 0.0 | | | | | | |
| DbiMA2 | 4.9 | 4.9 | 4.4 | 3.0 | 3.4 | 3.1 | 2.6 | 2.6 | 2.7 | 3.0 | 3.0 | 2.5 | 3.2 | 3.2 | 2.0 | 1.8 | 2.0 | 2.1 | 2.3 | 1.8 | 1.7 | 0.9 | 1.8 | 1.6 | 1.8 | 1.7 | 1.8 | 1.7 | 1.4 | 0.0 | | | | | | | |
| SveMA | 4.9 | 4.9 | 4.5 | 3.1 | 3.5 | 3.2 | 2.7 | 2.7 | 2.7 | 3.0 | 3.1 | 2.5 | 3.2 | 3.3 | 2.0 | 1.9 | 2.1 | 2.2 | 2.4 | 1.8 | 1.4 | 1.1 | 0.8 | 0.7 | 0.9 | 0.8 | 0.9 | 0.7 | 0.0 | | | | | | | | |
| GjuMA | 5.1 | 5.1 | 4.7 | 3.3 | 3.7 | 3.4 | 2.9 | 2.9 | 2.9 | 3.2 | 3.3 | 2.7 | 3.4 | 3.5 | 2.2 | 2.1 | 2.3 | 2.4 | 2.6 | 2.0 | 1.6 | 1.3 | 0.9 | 0.6 | 0.8 | 0.8 | 0.9 | 0.0 | | | | | | | | | |
| CeuMA | 5.3 | 5.3 | 4.9 | 3.5 | 3.8 | 3.5 | 3.1 | 3.1 | 3.1 | 3.4 | 3.5 | 2.9 | 3.6 | 3.6 | 2.4 | 2.3 | 2.5 | 2.6 | 2.8 | 2.2 | 1.7 | 1.5 | 1.1 | 0.7 | 0.7 | 0.7 | 0.0 | | | | | | | | | | |
| MroMA1 | 5.2 | 5.2 | 4.8 | 3.4 | 3.7 | 3.4 | 3.0 | 3.0 | 3.0 | 3.3 | 3.4 | 2.8 | 3.5 | 3.5 | 2.3 | 2.2 | 2.4 | 2.5 | 2.7 | 2.1 | 1.6 | 1.4 | 1.0 | 0.6 | 0.1 | 0.0 | | | | | | | | | | | |
| MroMA2 | 5.2 | 5.2 | 4.8 | 3.4 | 3.8 | 3.5 | 3.0 | 3.0 | 3.0 | 3.4 | 3.4 | 2.8 | 3.6 | 3.6 | 2.3 | 2.2 | 2.4 | 2.5 | 2.7 | 2.1 | 1.7 | 1.4 | 1.0 | 0.6 | 0.0 | | | | | | | | | | | | |
| PocMA | 5.1 | 5.1 | 4.6 | 3.3 | 3.6 | 3.3 | 2.8 | 2.8 | 2.9 | 3.2 | 3.2 | 2.7 | 3.4 | 3.4 | 2.2 | 2.1 | 2.2 | 2.4 | 2.5 | 2.0 | 1.5 | 1.2 | 0.9 | 0.0 | | | | | | | | | | | | | |
| PgiMA1 | 5.2 | 5.2 | 4.8 | 3.4 | 3.8 | 3.5 | 3.0 | 3.0 | 3.0 | 3.3 | 3.4 | 2.8 | 3.6 | 3.6 | 2.3 | 2.2 | 2.4 | 2.5 | 2.7 | 2.1 | 1.7 | 1.4 | 0.0 | | | | | | | | | | | | | | |
| AboMA | 4.5 | 4.5 | 4.1 | 2.7 | 3.0 | 2.7 | 2.3 | 2.3 | 2.3 | 2.6 | 2.7 | 2.1 | 2.8 | 2.8 | 1.6 | 1.5 | 1.7 | 1.8 | 2.0 | 1.4 | 1.3 | 0.0 | | | | | | | | | | | | | | | |
| RceM | 5.2 | 5.2 | 4.7 | 3.4 | 3.7 | 3.4 | 2.9 | 2.9 | 3.0 | 3.3 | 3.3 | 2.8 | 3.5 | 3.5 | 2.3 | 2.2 | 2.3 | 2.4 | 2.6 | 2.1 | 0.0 | | | | | | | | | | | | | | | | |
| SliM | 4.7 | 4.7 | 4.3 | 2.9 | 3.3 | 2.9 | 2.5 | 2.5 | 2.5 | 2.8 | 2.9 | 2.3 | 3.0 | 3.1 | 1.8 | 1.7 | 1.9 | 2.0 | 2.2 | 0.0 | | | | | | | | | | | | | | | | | |
| PmoM | 4.9 | 4.9 | 4.4 | 3.1 | 3.4 | 3.4 | 2.9 | 2.9 | 2.9 | 3.2 | 3.3 | 2.7 | 3.4 | 3.5 | 2.2 | 2.1 | 1.4 | 1.3 | 0.0 | | | | | | | | | | | | | | | | | | |
| PruM | 4.7 | 4.7 | 4.3 | 2.9 | 3.3 | 3.2 | 2.7 | 2.7 | 2.7 | 3.1 | 3.1 | 2.5 | 3.3 | 3.3 | 2.0 | 1.9 | 1.2 | 0.0 | | | | | | | | | | | | | | | | | | | |
| AinM | 4.6 | 4.6 | 4.1 | 2.8 | 3.1 | 3.0 | 2.6 | 2.6 | 2.6 | 2.9 | 3.0 | 2.4 | 3.1 | 3.2 | 1.9 | 1.8 | 0.0 | | | | | | | | | | | | | | | | | | | | |
| SurM1 | 4.7 | 4.7 | 4.2 | 2.8 | 3.2 | 2.2 | 1.7 | 1.7 | 1.7 | 2.1 | 2.1 | 1.5 | 2.3 | 2.3 | 0.7 | 0.0 | | | | | | | | | | | | | | | | | | | | | |
| SurM2 | 4.8 | 4.8 | 4.3 | 3.0 | 3.3 | 2.3 | 1.8 | 1.8 | 1.8 | 2.2 | 2.2 | 1.7 | 2.4 | 2.4 | 0.0 | | | | | | | | | | | | | | | | | | | | | | |
| LepM1 | 6.0 | 6.0 | 5.6 | 4.2 | 4.6 | 3.0 | 2.6 | 2.5 | 2.6 | 2.9 | 2.9 | 2.4 | 1.7 | 0.0 | | | | | | | | | | | | | | | | | | | | | | | |
| LepM2 | 6.0 | 6.0 | 5.5 | 4.2 | 4.5 | 3.0 | 2.5 | 2.5 | 2.6 | 2.9 | 2.9 | 2.4 | 0.0 | | | | | | | | | | | | | | | | | | | | | | | | |
| SspM | 5.3 | 5.3 | 4.8 | 3.5 | 3.8 | 2.2 | 1.8 | 1.8 | 1.8 | 2.1 | 2.2 | 0.0 | | | | | | | | | | | | | | | | | | | | | | | | | |
| SamM | 5.8 | 5.8 | 5.4 | 4.0 | 4.4 | 1.9 | 2.0 | 1.5 | 1.6 | 1.9 | 0.0 | | | | | | | | | | | | | | | | | | | | | | | | | | |
| SanM | 5.8 | 5.8 | 5.3 | 4.0 | 4.3 | 2.0 | 1.9 | 0.6 | 0.6 | 0.0 | | | | | | | | | | | | | | | | | | | | | | | | | | | |
| SahM | 5.5 | 5.5 | 5.0 | 3.6 | 4.0 | 1.6 | 1.6 | 0.1 | 0.0 | | | | | | | | | | | | | | | | | | | | | | | | | | | | |
| SonM | 5.4 | 5.4 | 5.0 | 3.6 | 4.0 | 1.6 | 1.6 | 0.0 | | | | | | | | | | | | | | | | | | | | | | | | | | | | | |
| ParM | 5.4 | 5.4 | 5.0 | 3.6 | 4.0 | 2.0 | 0.0 | | | | | | | | | | | | | | | | | | | | | | | | | | | | | | |
| LulM | 4.8 | 5.9 | 5.5 | 4.1 | 4.4 | 0.0 | | | | | | | | | | | | | | | | | | | | | | | | | | | | | | | |
| WP_210373447.1 | 3.8 | 3.8 | 3.4 | 2.4 | 0.0 | | | | | | | | | | | | | | | | | | | | | | | | | | | | | | | | |
| WP_064287858.1 | 3.8 | 3.8 | 3.4 | 0.0 | | | | | | | | | | | | | | | | | | | | | | | | | | | | | | | | | |
| WP_205004337.1 | 3.9 | 1.9 | 0.0 | | | | | | | | | | | | | | | | | | | | | | | | | | | | | | | | | | |
| WP_101772534.1 | 4.3 | 0.0 | | | | | | | | | | | | | | | | | | | | | | | | | | | | | | | | | | | |
| WP_106347317.1 | 0.0 | | | | | | | | | | | | | | | | | | | | | | | | | | | | | | | | | | | | |

**Figure S13: Sum of branch distances between SspM homologs.** The sum of the branch distances between a subset of SspM homologs. Included are the distances for all characterized borosin methyltransferases, predicted type IV borosin methyltransferases (WP_210373447.1 and WP 064287858.1), and non-borosin TP methylase outgroups (WP_106347317.1, WP_205004337.1, and WP 101772534.1).

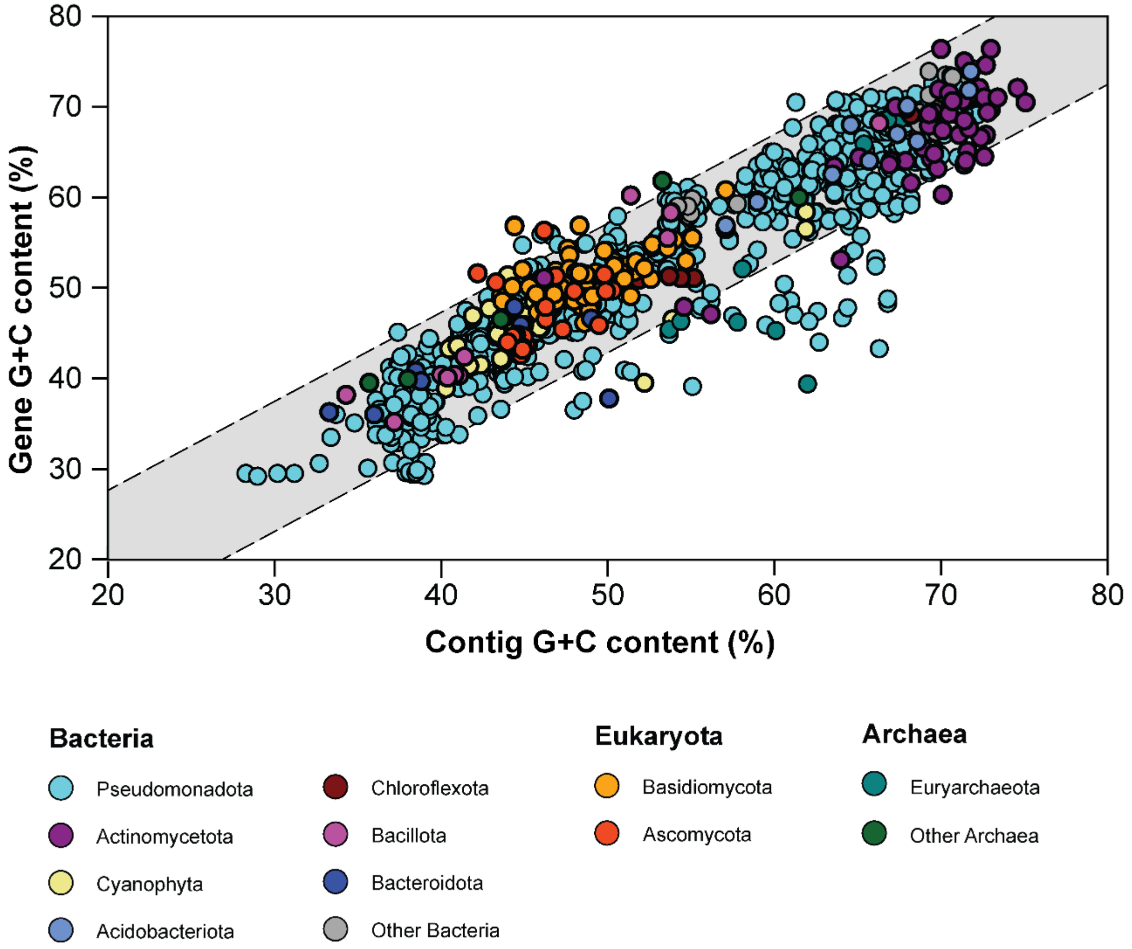


**Figure S14: G+C content analysis of borosin methyltransferases.** The G+C content of borosin methyltransferases compared to the G+C content of the contig on which they are encoded. The gray highlighted area shows the genes with a G+C difference within 2 standard deviations of the mean. The bottom right corner of the graph represents genes likely originating in lower G+C content organisms that were horizontally transferred to a higher G+C content organism.

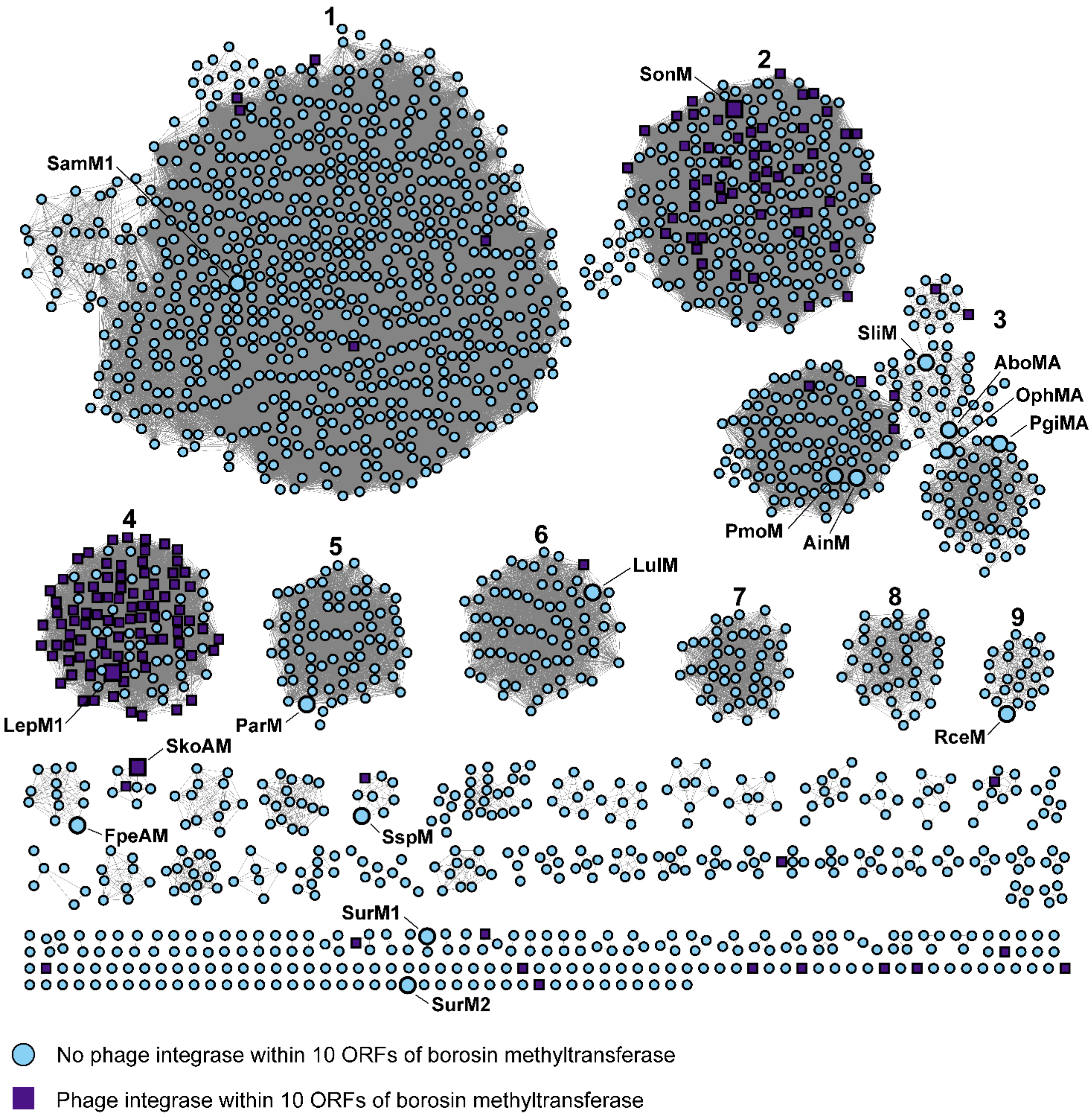


**Figure S15: SSN of borosin methyltransferases with nearby phage integrases.** Borosin methyltransferases with a RODEO-identified precursor peptide were used to generate this SSN (*n* = 2,124, alignment score = 90).

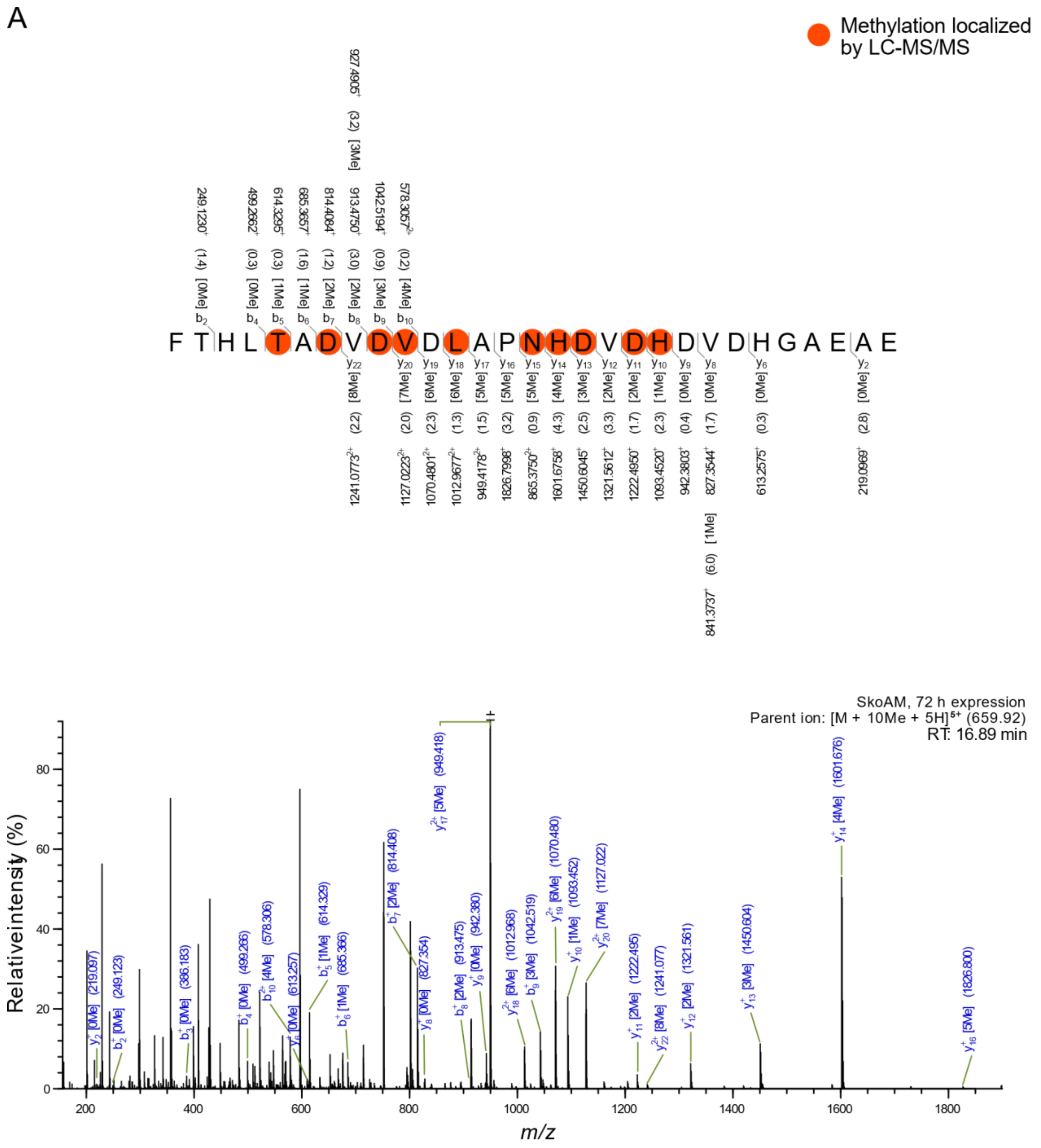
A
Methylation localized by LC-MS/MS
F T H L T A D V D V D L A P N H D V D H D V D H G A E A E
SkoAM, 72 h expression
Parent ion: [M + 10Me + 5H]5+ (659.92)
RT: 16.89 min
Relative intensity (%)
m/z

B

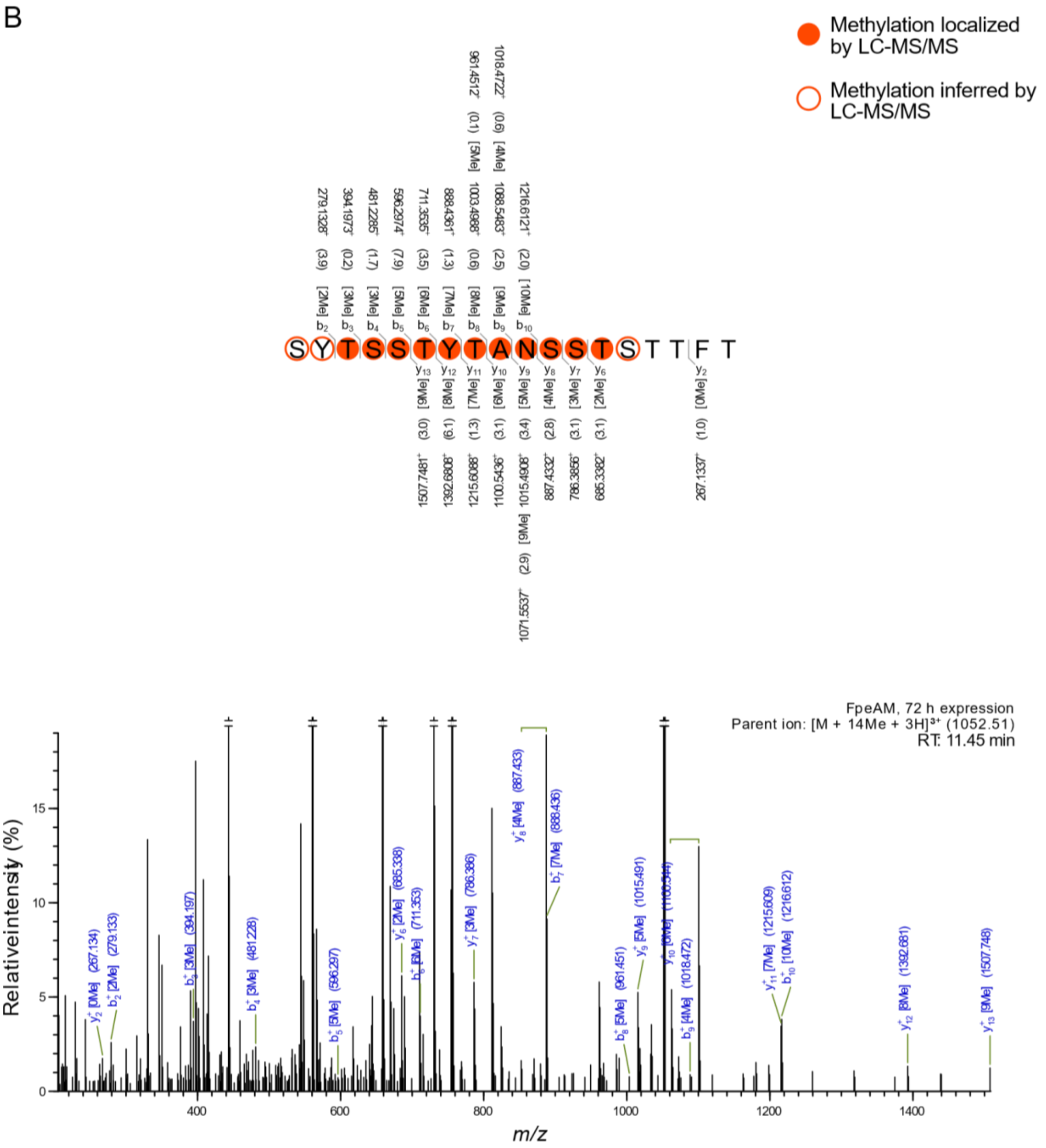


**Figure S16: MS2 analysis of type 0 borosin core peptide regions.** Nickel affinity resin-purified proteins were in-gel digested then analyzed using high resolution LC-MS/MS with HCD fragmentation to identify modified residues. Methylation state of each peptide fragment is included in brackets, where 'Me' marks a mass shift corresponding to methylation, and methylated residues are indicated by orange circles. $b^+$ and $y^+$ ions are labeled above and below the peptide sequence with the observed m/z and the deviation from the theoretical mass in parentheses (in ppm). A mass cutoff of 10.0 ppm and minimum relative peak intensity of 0.1% was used for the annotated masses. **A.** MS2 fragmentation and methylation pattern for GluC digested SkoAM (72 hour expression). **B.** MS2 fragmentation and methylation pattern for Proteinase K digested FpeAM (72 hour expression).

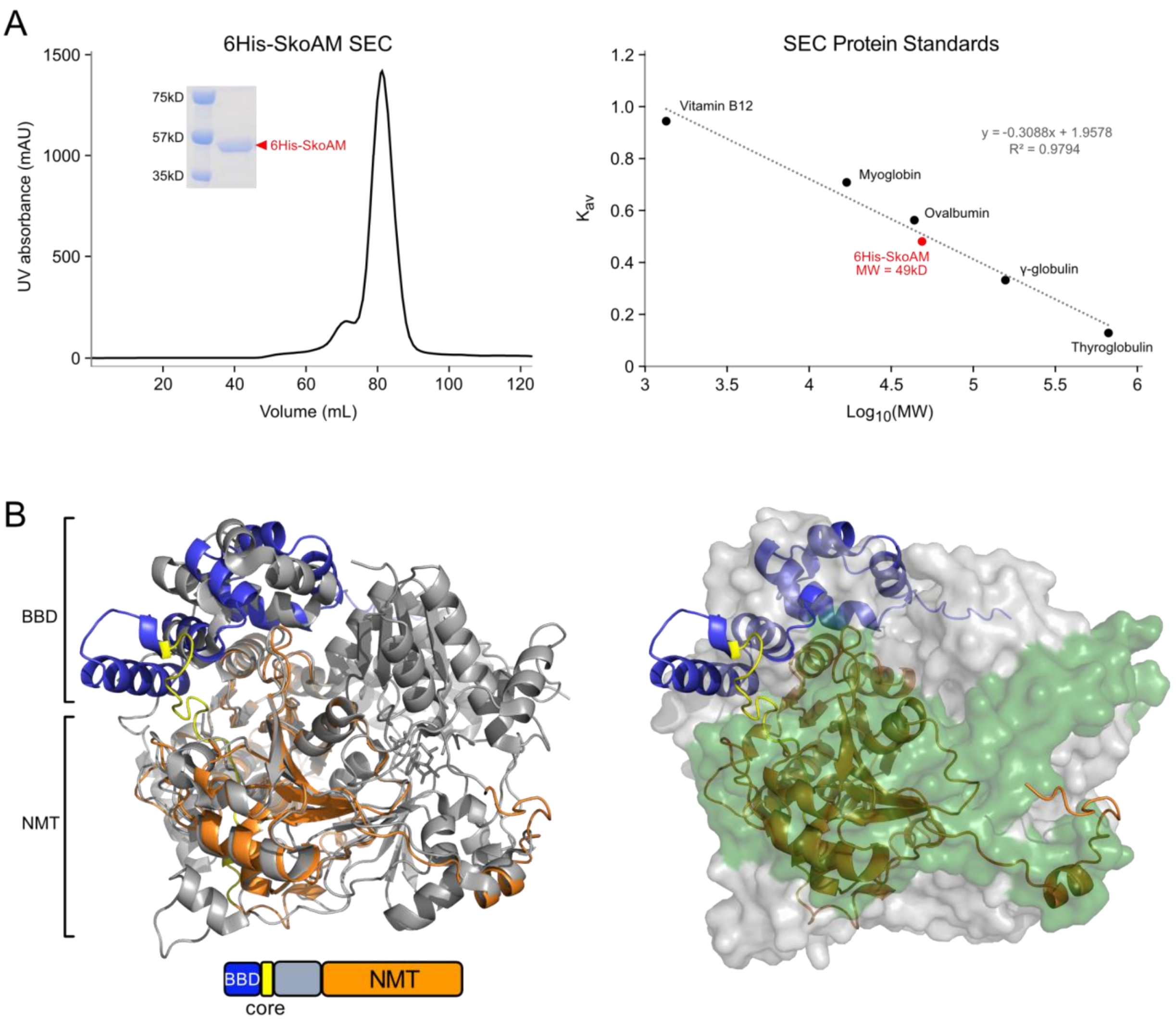


**Figure S17: SEC and Alphafold model for SkoAM.** SEC and Alphafold structural predictions suggest 6His-SkoAM is a monomer **A.** 6His-SkoAM plotted onto a calibration curve (right) using its theoretical monomeric mass (49 kD) and the partition coefficient ($K_{av}$) calculated from the observed retention time from the SEC chromatogram (left). An inset panel of the SDS-PAGE profile for the SEC peak is included. For calibration of the column the following size markers were used: vitamin B12 (1.35 kD), myoglobin (17 kD), ovalbumin (44 kD), γ-globulin (158 kD), and thyroglobulin (670 kD). The Y-axis represents the partition coefficient ($K_{av}$) and X-axis represents the log of the molecular weight (MW). **B.** Superimpositions of an Alphafold generated structure of monomeric SkoAM (cartoon representation) and the OphMA crystal structure (PDB: 5N0X) in grey cartoon representation (left) and surface representation (right). The SkoAM BBD domain is colored blue, the core yellow, while the NMT domain is colored in orange. The two subunits of the surface representation of the OphMA homodimer are separately colored green and grey. An Alphafold generated structure of homodimeric SkoAM showed a similar configuration (not shown).

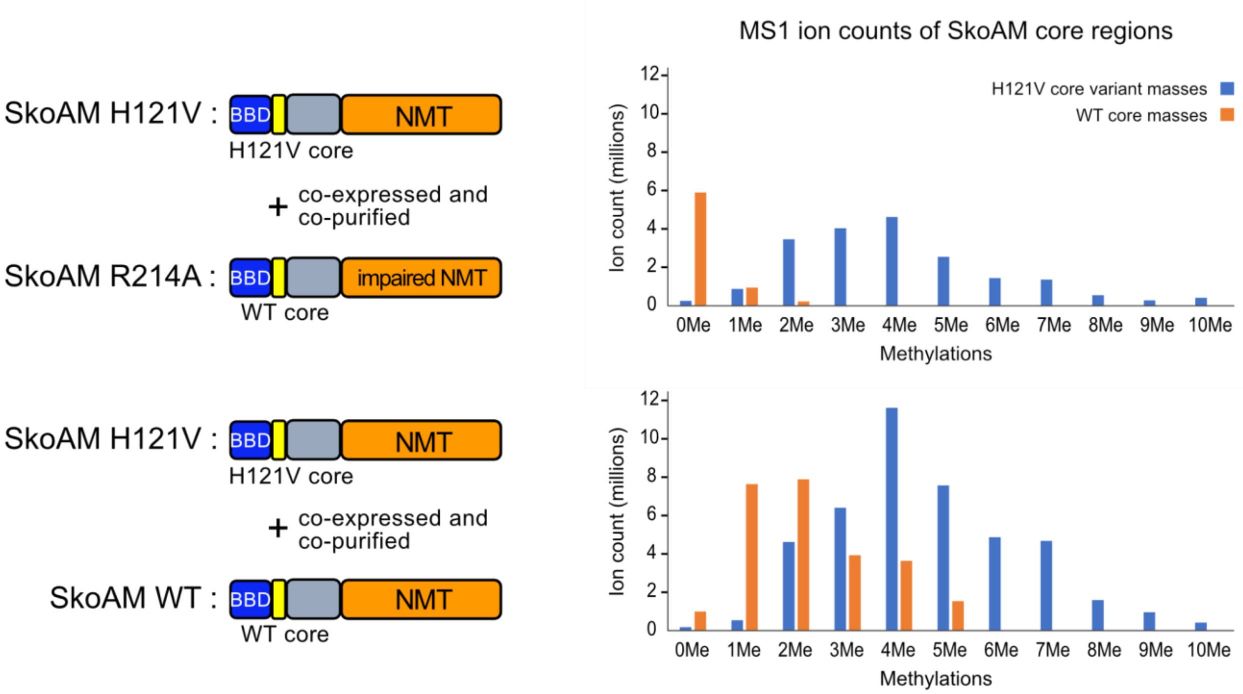


**Figure S18: SkoAM *trans*-complementation assay.** Complementation experiments to determine the intra- or inter- molecular mechanism of SkoAM. A distinguishable SkoAM core mutant (H121V) was co-expressed and co-purifed with either an impaired SkoAM R214A mutant (top) or wildtype (WT) (bottom) SkoAM then analyzed by LC-MS/MS. The ion counts (y-axis) for MS1 parent ions (including 2+ and 3+ charge states with a mass difference cutoff of 10 mmu) containing 0-10 methylations (x-axis) are shown with masses representing the H121V core variant shown in blue and those for the WT core shown in orange.

A

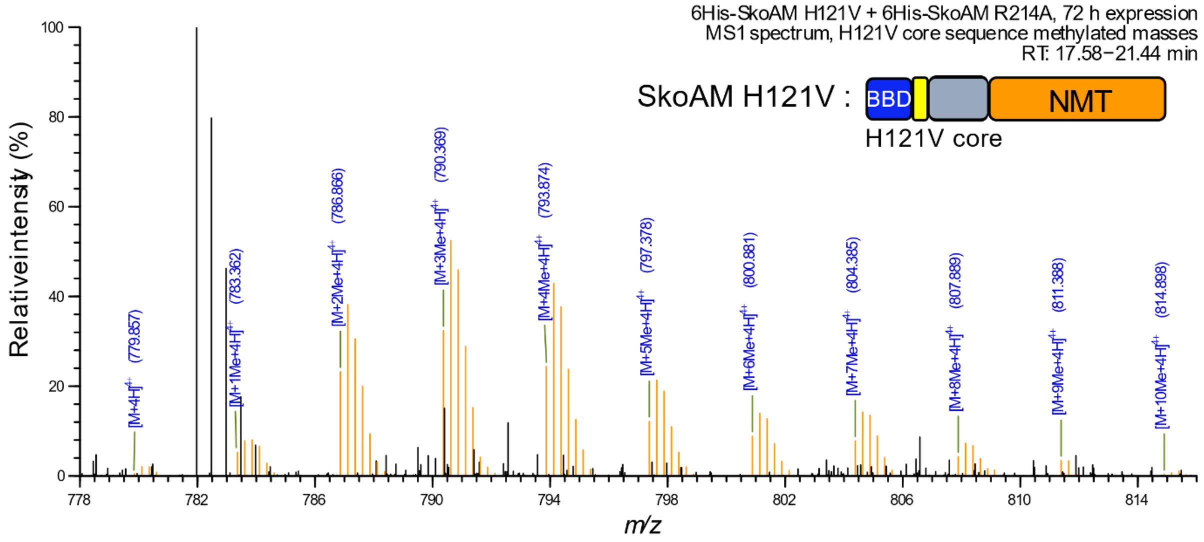
6His-SkoAM H121V + 6His-SkoAM R214A, 72 h expression
MS1 spectrum, H121V core sequence methylated masses
RT: 17.58−21.44 min
SkoAM H121V :
BBD
NMT
H121V core
Relative intensity (%)
m/z
[M+4H]4+ (779.857)
[M+1Me+4H]4+ (783.362)
[M+2Me+4H]4+ (786.866)
[M+3Me+4H]4+ (790.369)
[M+4Me+4H]4+ (793.874)
[M+5Me+4H]4+ (797.378)
[M+6Me+4H]4+ (800.881)
[M+7Me+4H]4+ (804.385)
[M+8Me+4H]4+ (807.889)
[M+9Me+4H]4+ (811.388)
[M+10Me+4H]4+ (814.898)

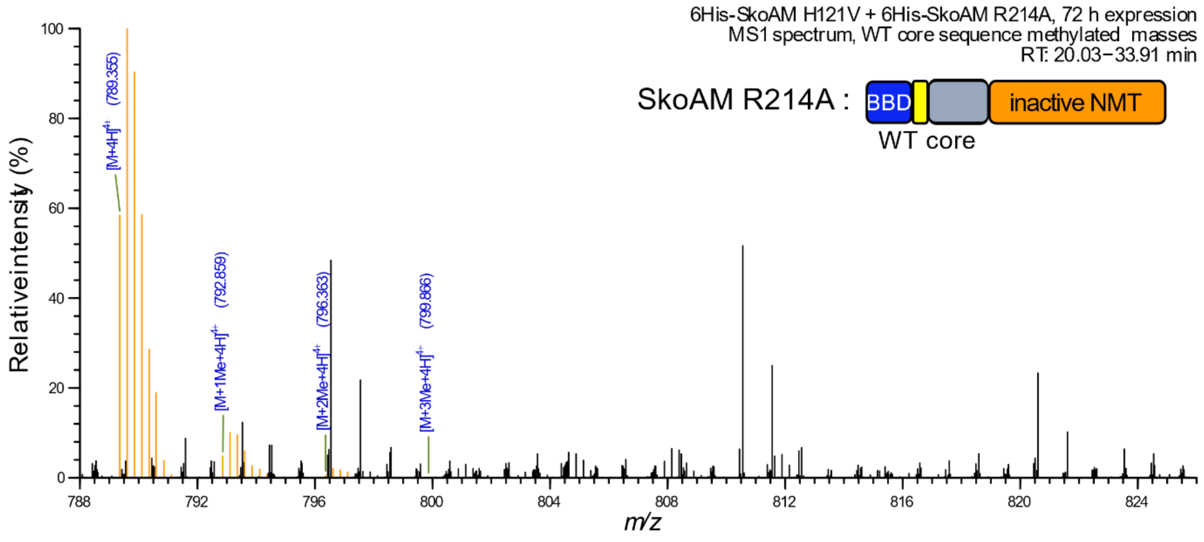
6His-SkoAM H121V + 6His-SkoAM R214A, 72 h expression
MS1 spectrum, WT core sequence methylated masses
RT: 20.03−33.91 min
SkoAM R214A :
BBD
inactive NMT
WT core
Relative intensity (%)
m/z
[M+4H]4+ (789.355)
[M+1Me+4H]4+ (792.859)
[M+2Me+4H]4+ (796.363)
[M+3Me+4H]4+ (799.866)

B

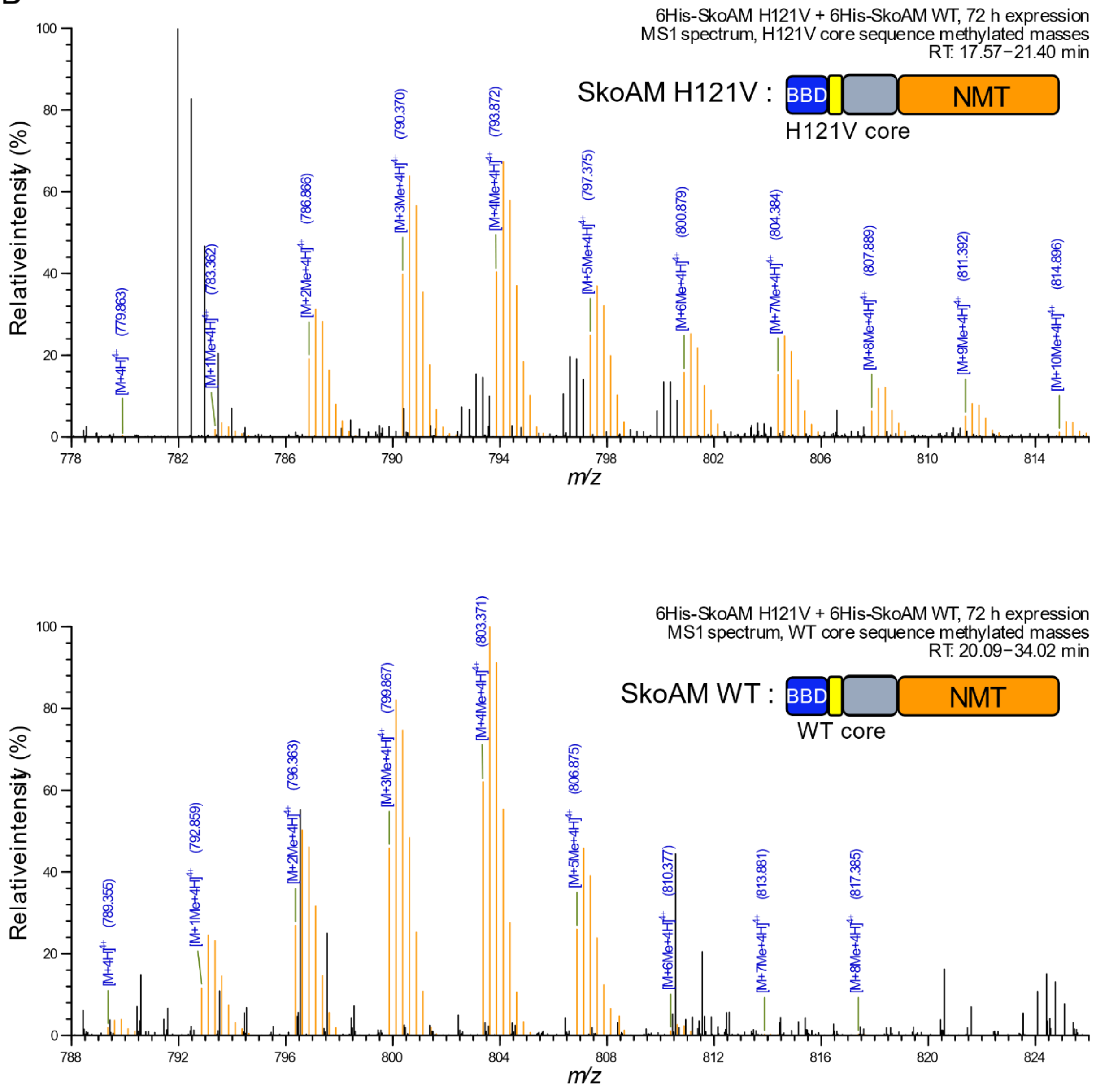


**Figure S19: MS1 spectra for SkoAM *trans*-complementation assay.** Observed MS1 masses (4+ charge state) for SkoAM core regions containing 0-10 methylations. Peaks within 10 ppm of the theoretical masses and their isotopic masses are labeled with their observed masses and corresponding methylation state and colored orange. **A.** Observed methylation states from the co-expression of 6His-SkoAM H121V + 6His-SkoAM R214A during retention times 17.58-21.44 min for the H121V mutant core (top) and 20.03-33.91 min for the WT core (bottom). **B.** Observed methylation states from the co-expression of 6His-SkoAM H121V + 6His-SkoAM WT during retention times 17.57-21.40 min for the H121V mutant core (top) and 20.09-34.02 min for the WT core (bottom).

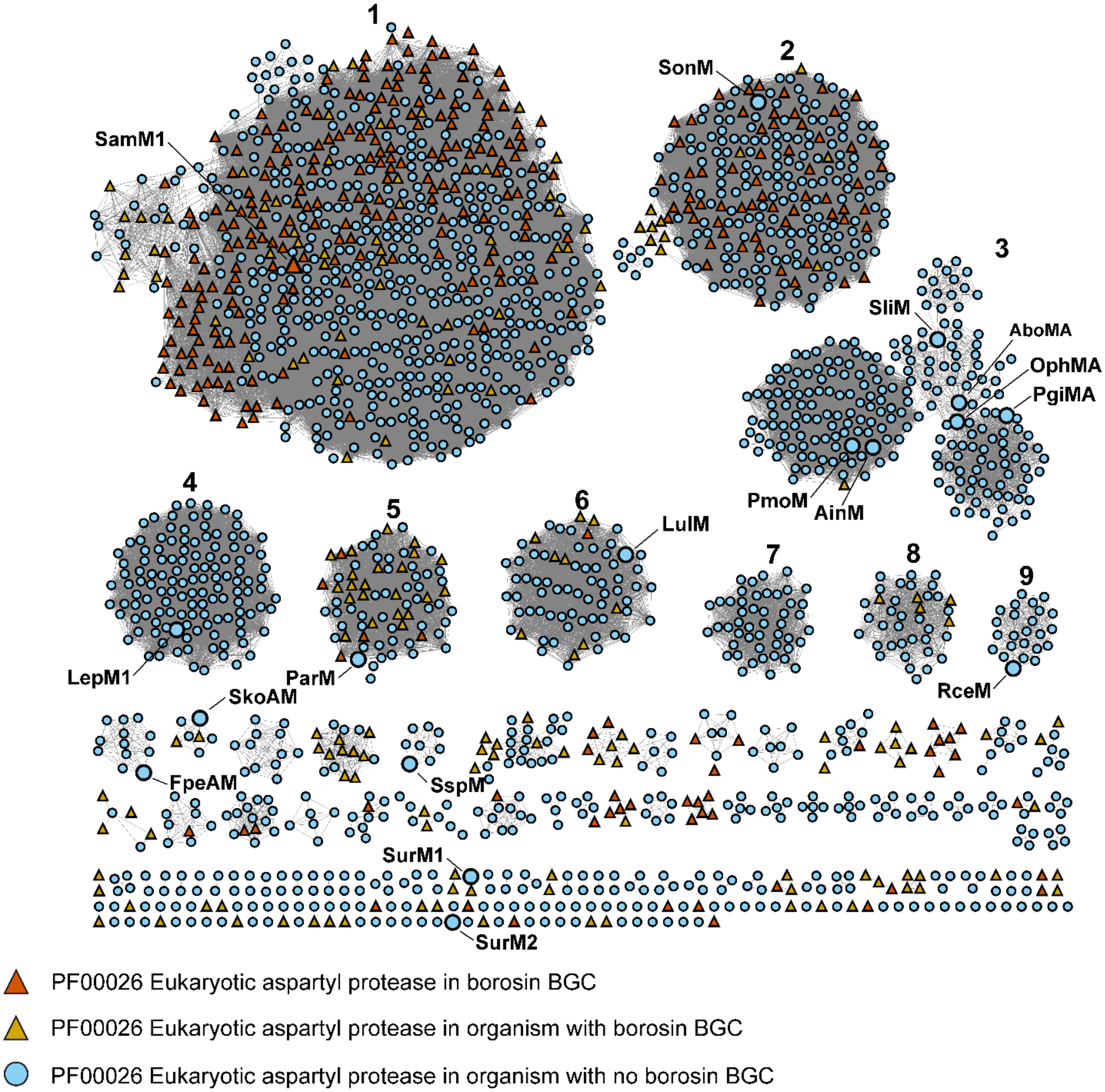


**Figure S20: SSN of borosin methyltransferases associated with a PF00026 eukaryotic aspartyl protease.** Borosin methyltransferases with a RODEO identified precursor peptide were used to generate this SSN (*n* = 2,124, alignment score = 90).

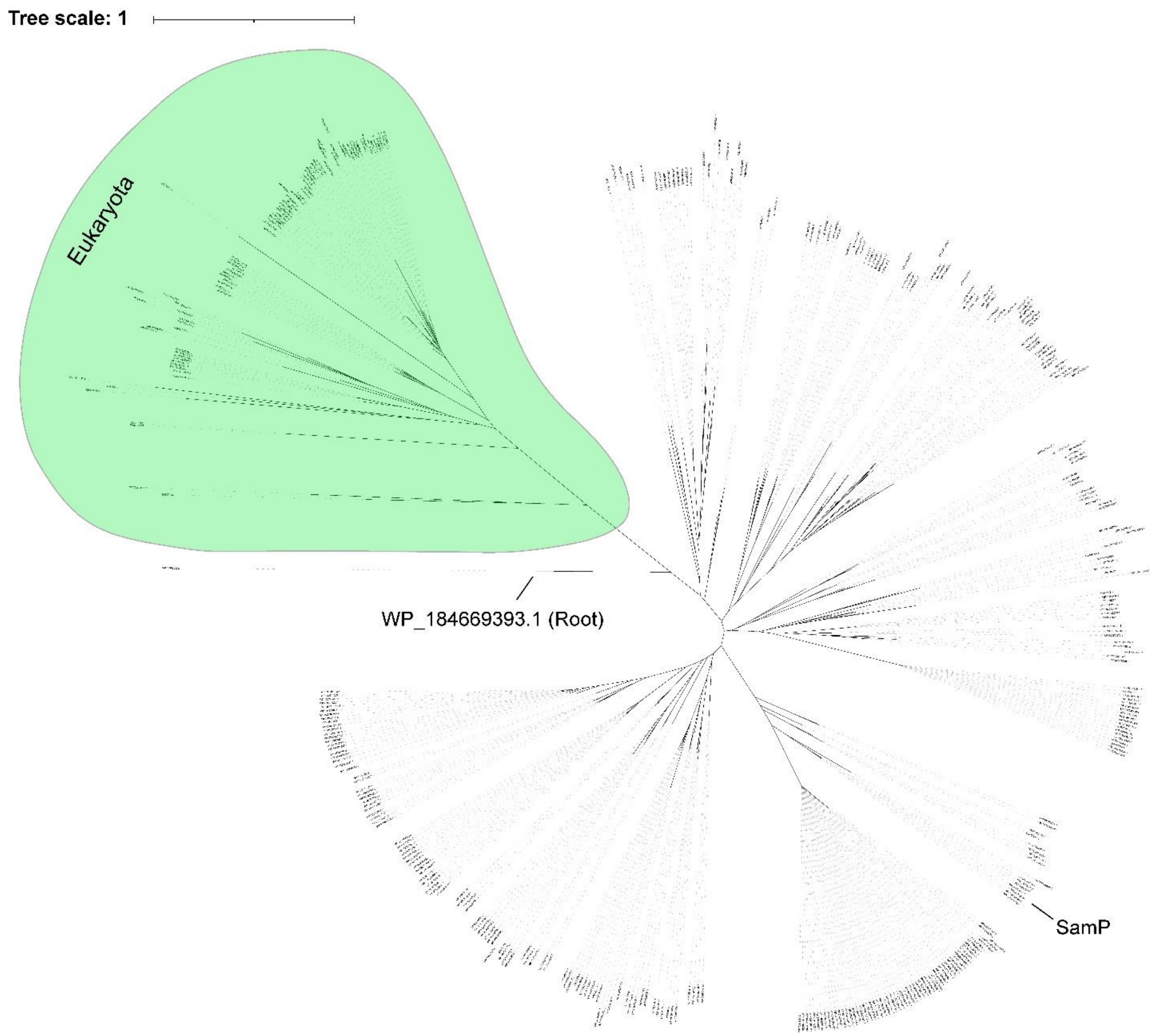


**Figure S21: Unrooted tree of SamP homologs.** A phylogenetic tree of 441 SamP homologs found in Bacteria and Eukarya.

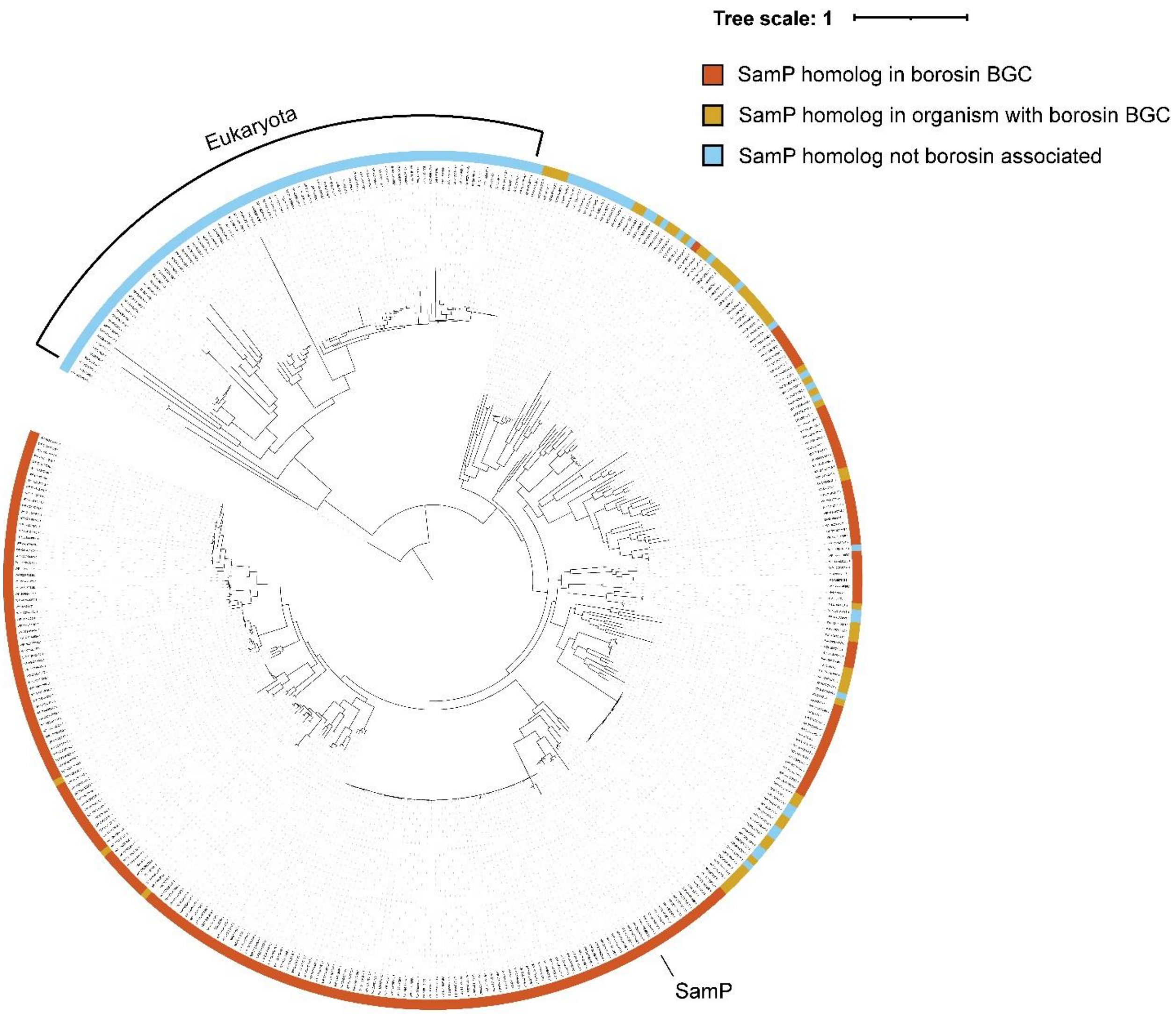


**Figure S22: Rooted tree of SamP homologs.** A phylogenetic tree of 441 SamP homologs found in Bacteria and Eukarya rooted using WP_184669393.1 to show separation between Eukaryotic and Bacterial sequences.

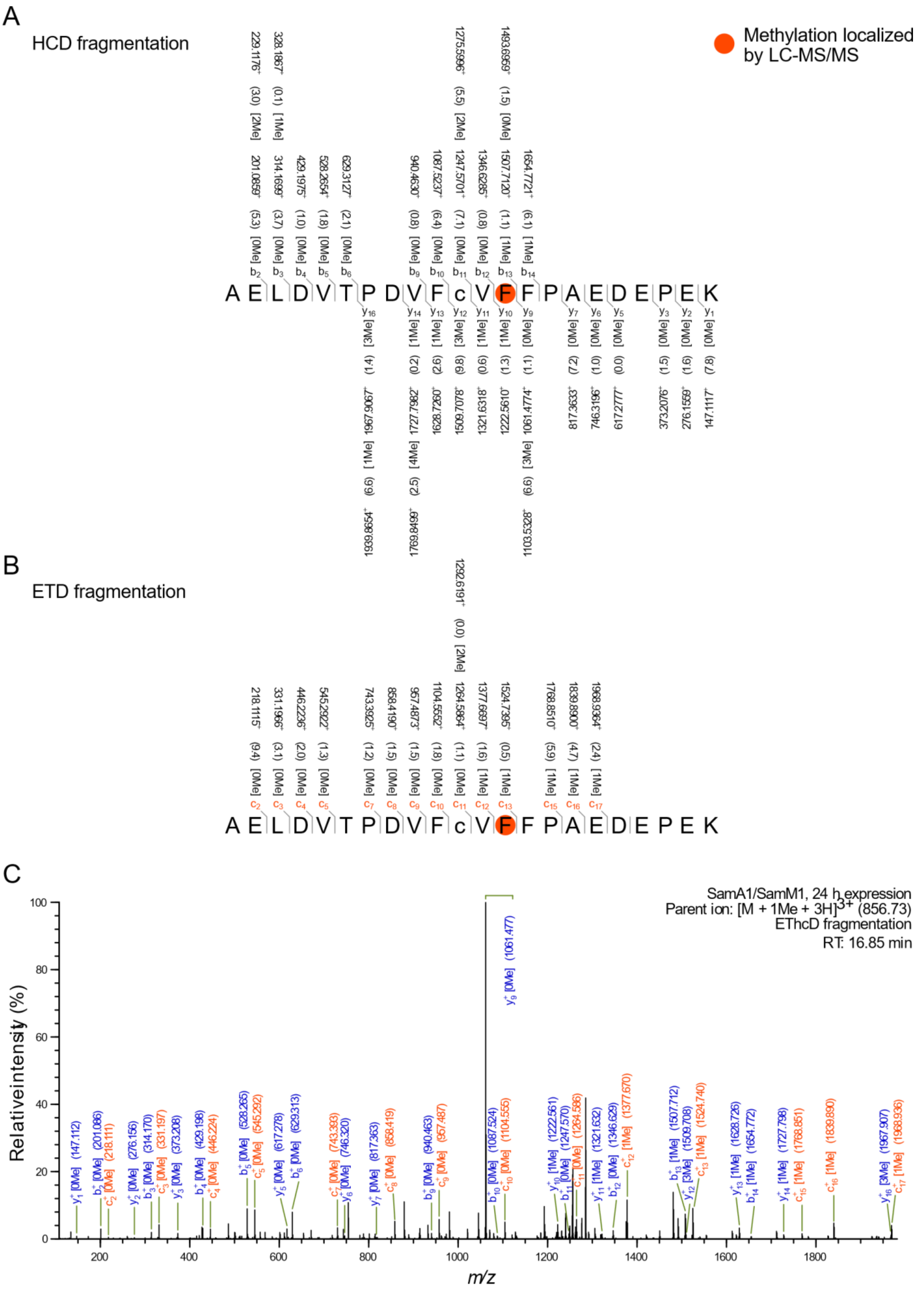

A
HCD fragmentation
Methylation localized by LC-MS/MS
A E L D V T P D V F c V F F P A E D E P E K
B
ETD fragmentation
A E L D V T P D V F c V F F P A E D E P E K
C
SamA1/SamM1, 24 h expression
Parent ion: [M + 1Me + 3H]3+ (856.73)
EThcD fragmentation
RT: 16.85 min
Relative intensity (%)
m/z

**Figure S23: MS2 fragmentation of methylated SamA1.** SamA1, purified from a co-expression with *samM1,* in-gel digested with trypsin, then analyzed using high resolution LC-MS/MS with EThcD fragmentation. Methylation state of each peptide fragment is included in brackets, where 'Me' marks a mass shift corresponding to methylation, and methylated residues are indicated by orange circles. Fragment $b^+$, $y^+$, and $c^+$ ions are labeled above and below the peptide sequence with the observed m/z and the deviation from the theoretical mass in parentheses (in ppm). A mass cutoff of 10.0 ppm and minimum relative peak intensity of 0.1% was used for the annotated masses. **A.** Fragment $b^+$ and $y^+$ ions from HCD fragmentation **B.** $c^+$ ions from ETD fragmentation. **C.** MS2 spectrum with observed $b^+$, $y^+$, and $c^+$ ions (orange) labeled.

A

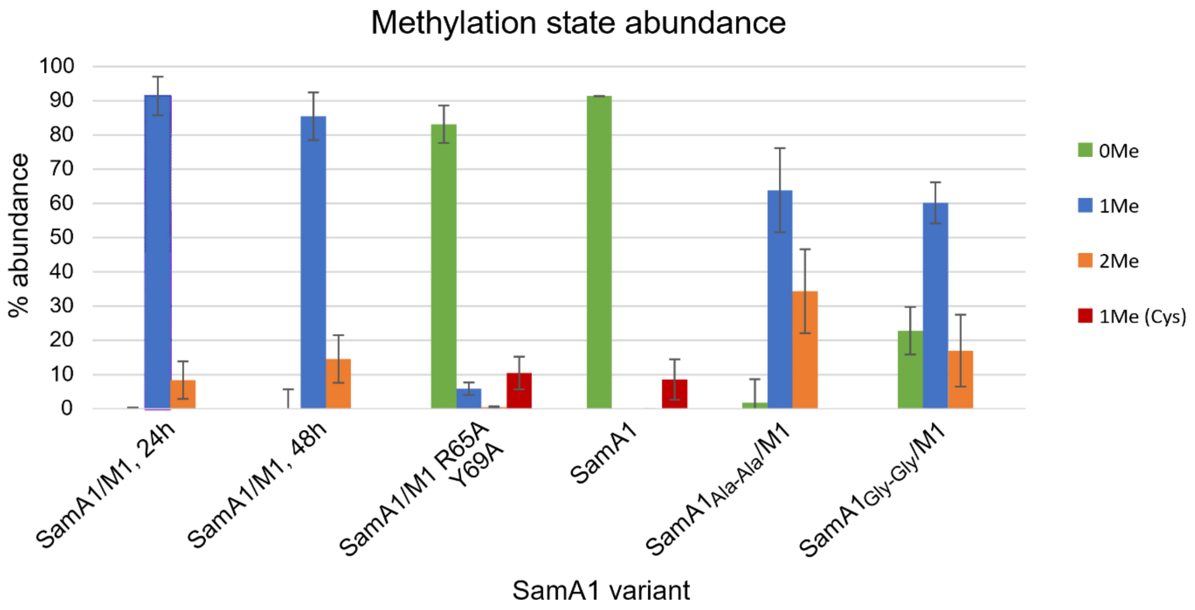


B

| Variant | 0Me | 1Me | 1Me (Cys) | 2Me |
|---|---|---|---|---|
| SamA1/M1, 24h | 0.20% +/- 0.22 | 91.40% +/- 5.67 | | 8.38% +/- 5.44 |
| SamA1/M1, 48h | 0.00% +/- 0.01 | 85.45% +/- 6.94 | | 14.53% +/- 6.94 |
| SamA1/M1 R65A Y69A | 83.12% +/- 3.18 | 5.86% +/- 1.81 | 10.4% +/- 4.80 | 0.51% +/- 0.19 |
| SamA1 | 91.40% +/- 5.90 | | 8.59% +/- 5.90 | |
| $SamA1_{Ala-Ala}$ /M1 | 1.75% +/- 0.61 | 63.86% +/- 12.2 | | 34.38% +/- 12.2 |
| $SamA1_{Gly-Gly}$ /M1 | 22.80% +/- 4.52 | 60.19% +/- 6.02 | | 16.99% +/- 10.5 |

**Figure S24: Calculated methylation state abundances of SamA1 variants.** Abundances of 0-2 methylation states for SamA1 and variants determined through in-gel digestion and high-resolution LC-MS/MS with HCD fragmentation. Calculated abundance includes $2^+$ and $3^+$ charged states with a 10 amu mass difference cutoff. Samples were prepared in technical triplicate. **A.** Bar graph of relative abundance of 0 methylations (green), 1 methylation at position 52 (blue), 1 methylation on Cys 50 (red), and 2 methylations (orange). Standard deviation is shown as error bars. **B.** Table of calculated abundances for each variant and methylation state.

A

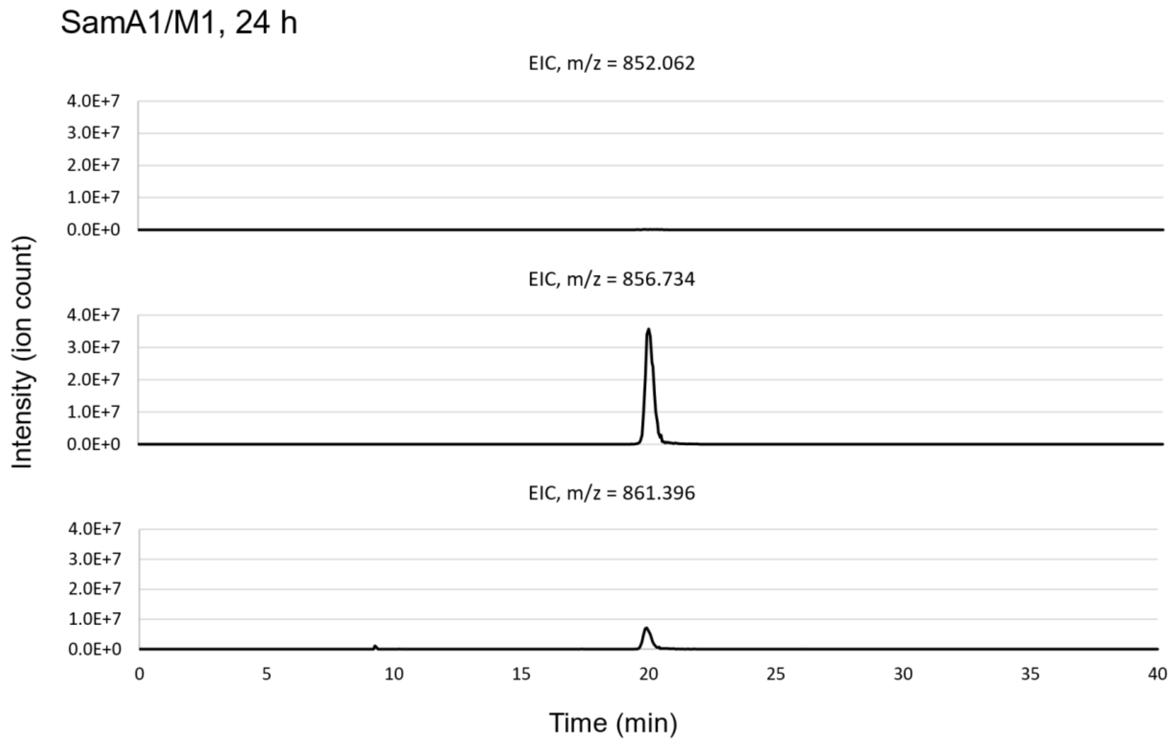
SamA1/M1, 24 h
EIC, m/z = 852.062
4.0E+7
3.0E+7
2.0E+7
1.0E+7
0.0E+0
EIC, m/z = 856.734
4.0E+7
3.0E+7
2.0E+7
1.0E+7
0.0E+0
EIC, m/z = 861.396
4.0E+7
3.0E+7
2.0E+7
1.0E+7
0.0E+0
Intensity (ion count)
0
5
10
15
20
25
30
35
40
Time (min)

B

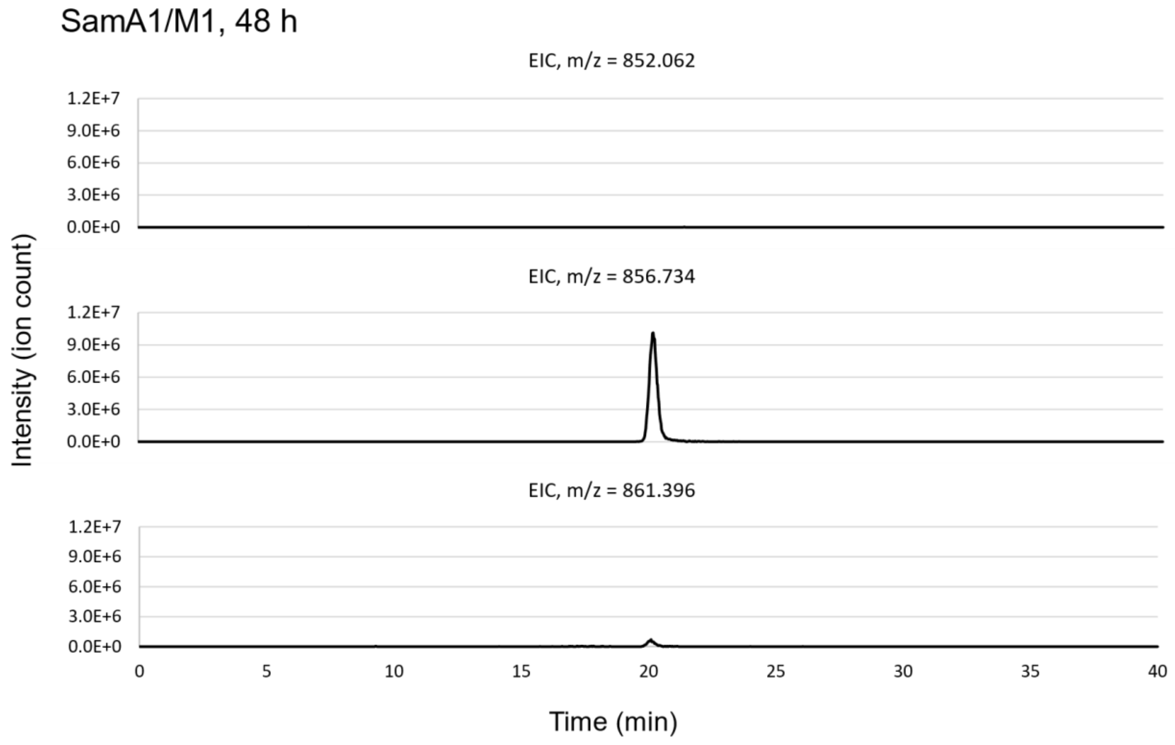
SamA1/M1, 48 h
EIC, m/z = 852.062
1.2E+7
9.0E+6
6.0E+6
3.0E+6
0.0E+0
EIC, m/z = 856.734
1.2E+7
9.0E+6
6.0E+6
3.0E+6
0.0E+0
EIC, m/z = 861.396
1.2E+7
9.0E+6
6.0E+6
3.0E+6
0.0E+0
Intensity (ion count)
0
5
10
15
20
25
30
35
40
Time (min)

C

SamA1/M1 R65A Y69A

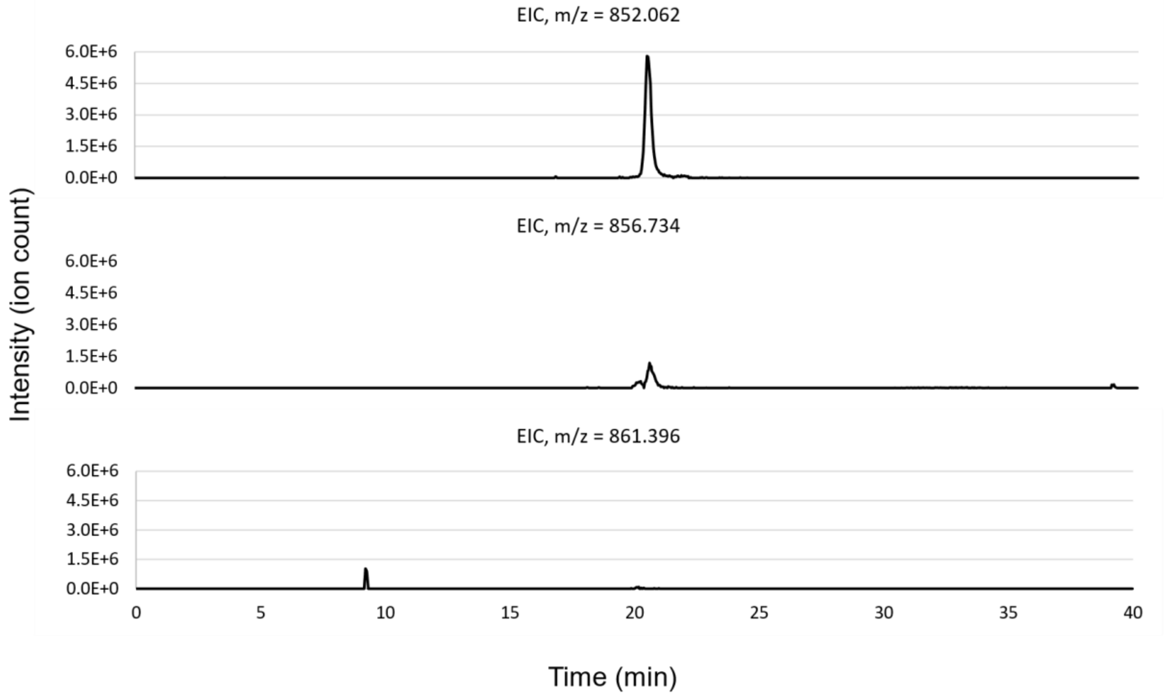


D

SamA1

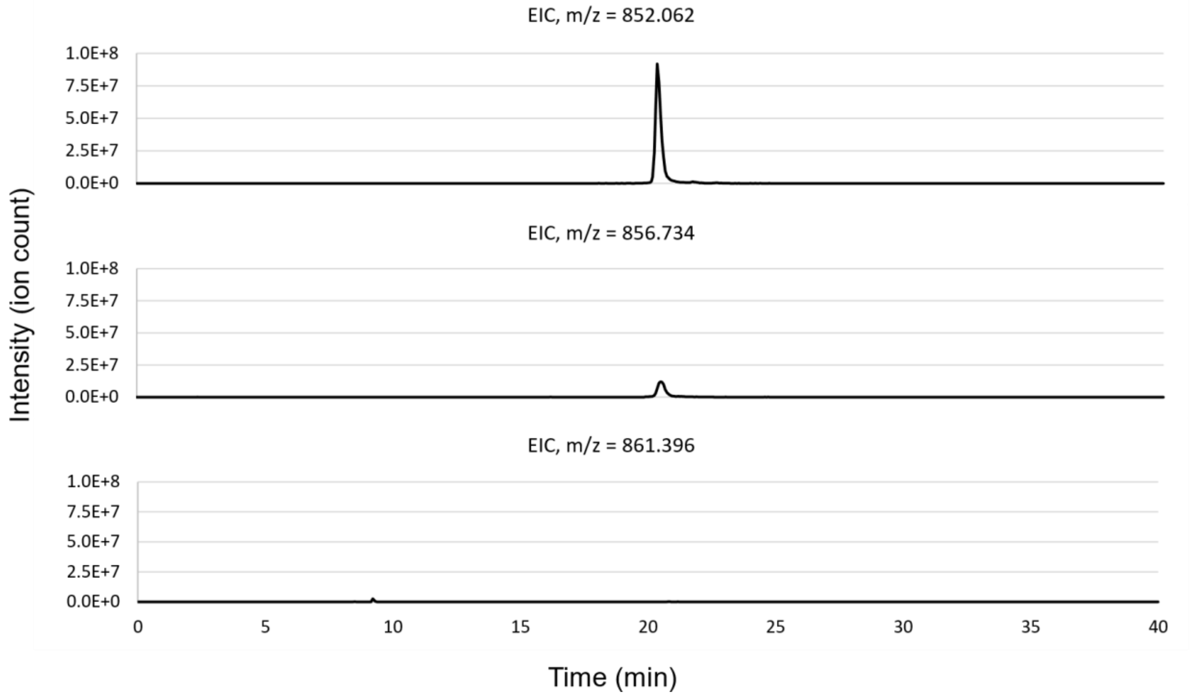

E

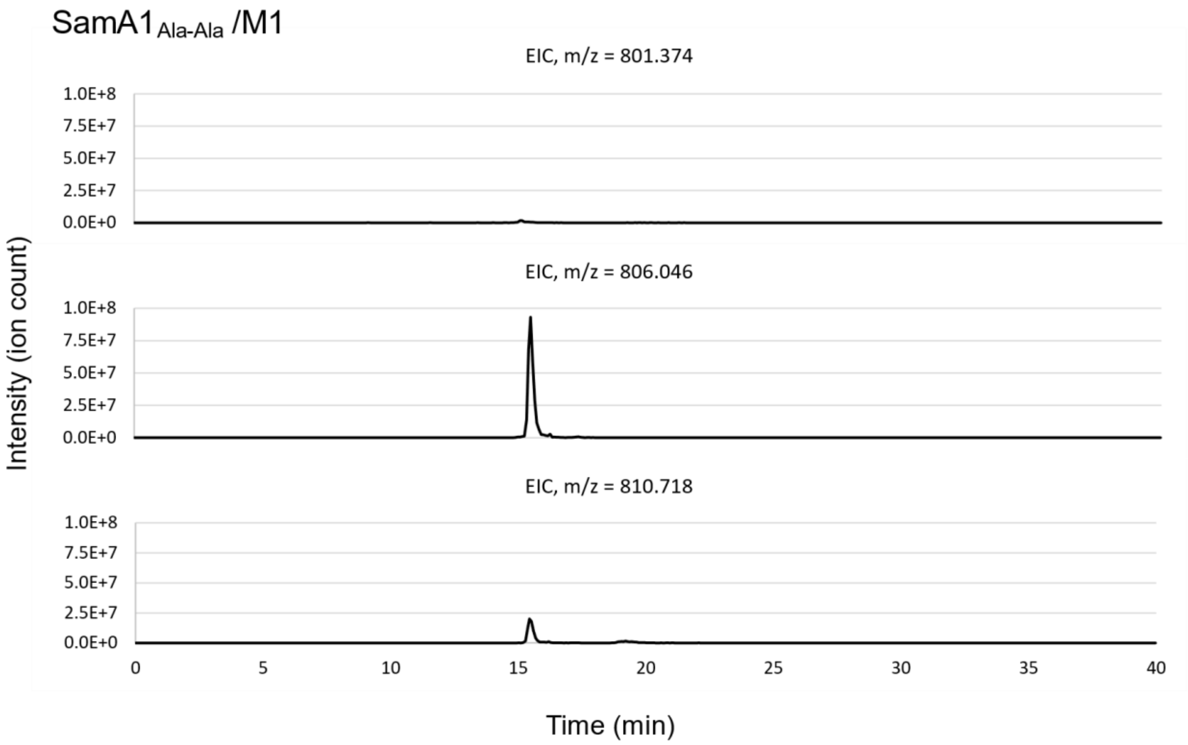
SamA1Ala-Ala /M1
EIC, m/z = 801.374
EIC, m/z = 806.046
EIC, m/z = 810.718
1.0E+8
7.5E+7
5.0E+7
2.5E+7
0.0E+0
Intensity (ion count)
0
5
10
15
20
25
30
35
40
Time (min)

F

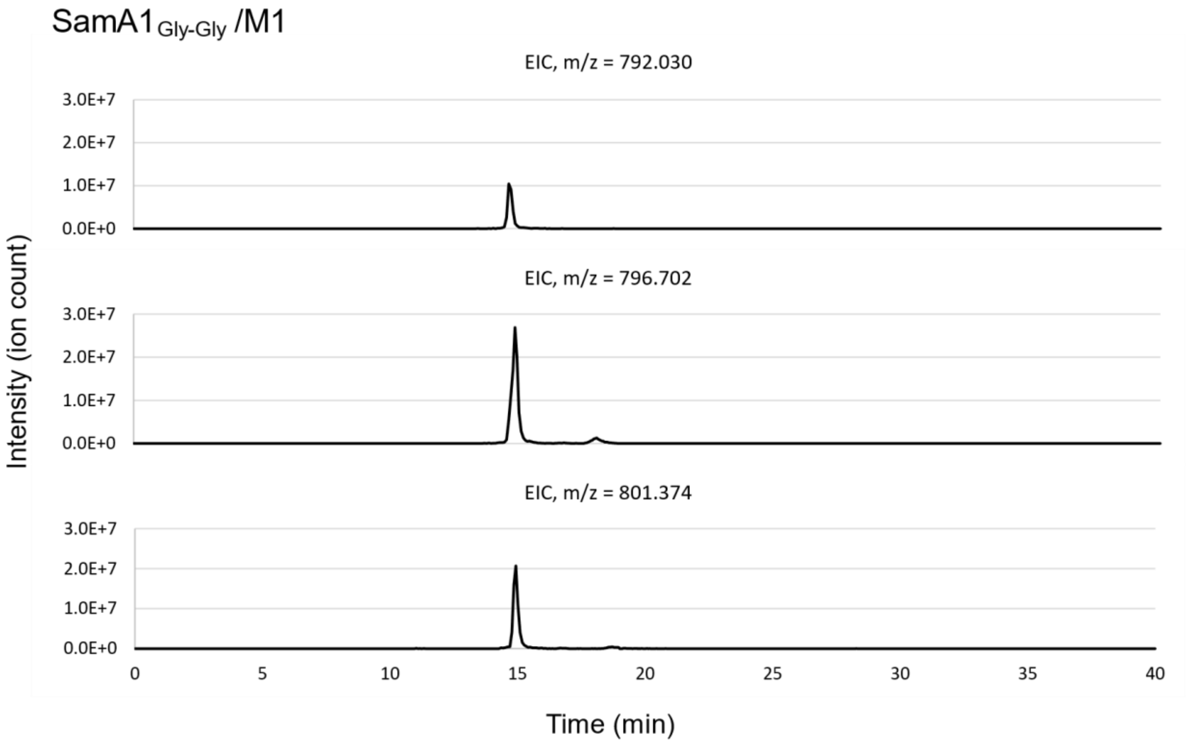
SamA1Gly-Gly /M1
EIC, m/z = 792.030
EIC, m/z = 796.702
EIC, m/z = 801.374
3.0E+7
2.0E+7
1.0E+7
0.0E+0
Intensity (ion count)
0
5
10
15
20
25
30
35
40
Time (min)

**Figure S25: LC-MS1 EICs for 0-2 methylation states of trypsinized SamA1 variants.** Extracted ion chromatograms (EICs) from a single replicate for each SamA1 variant. Included are masses for 0 (top), 1 (middle), and 2 (bottom) methylations at a $3^+$ charge state with a mass difference range of +/- 10 amu. **A.** EICs for SamA1/M1 expressed for 24 h. **B.** EICs for SamA1/M1 expressed for 48 h. **C.** EICs for SamA1/M1 R65A Y69A expressed for 48 h. **D.** EICs for SamA1 expressed for 24 h. **E.** EICs for $SamA1_{Ala\text{-}Ala}$/M1 expressed for 24 h. **F.** EICs for $SamA1_{Gly\text{-}Gly}$/M1 expressed for 24 h.

A

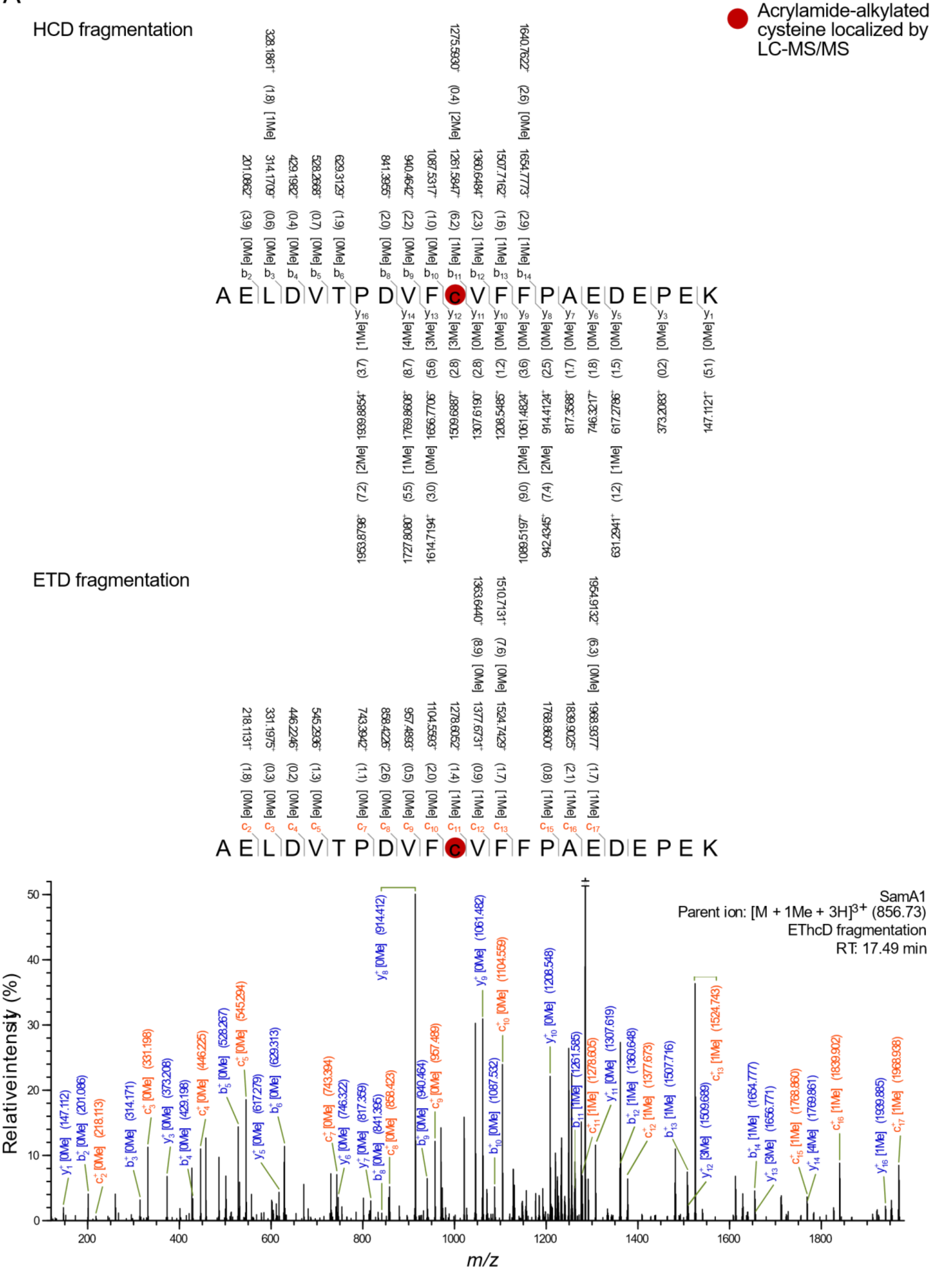

Acrylamide-alkylated cysteine localized by LC-MS/MS
HCD fragmentation
A E L D V T P D V F c V F F P A E D E P E K
ETD fragmentation
A E L D V T P D V F c V F F P A E D E P E K
SamA1
Parent ion: [M + 1Me + 3H]3+ (856.73)
EThcD fragmentation
RT: 17.49 min
Relative intensity (%)
m/z

B

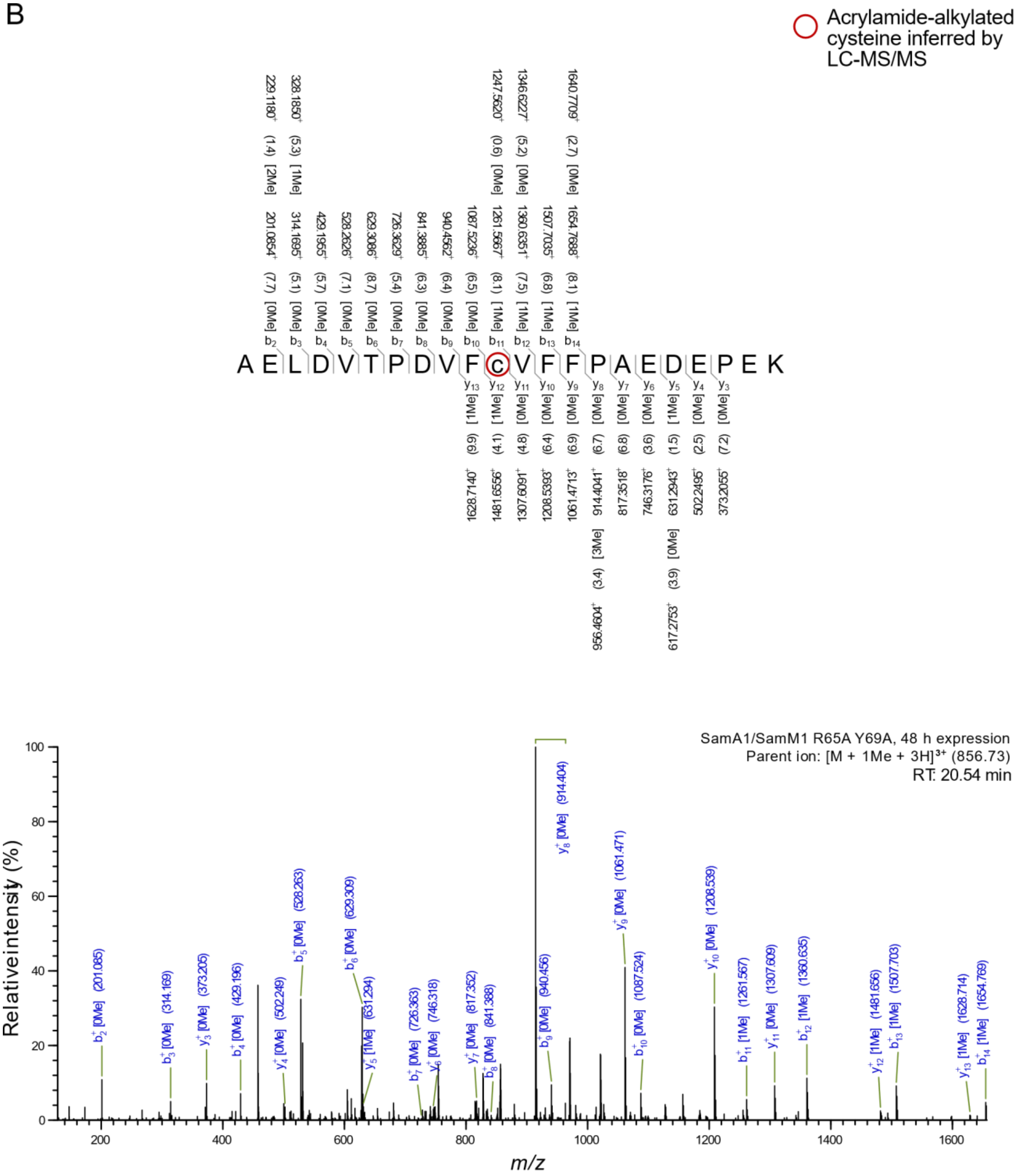
Acrylamide-alkylated cysteine inferred by LC-MS/MS
A E L D V T P D V F C V F F P A E D E P E K
SamA1/SamM1 R65A Y69A, 48 h expression
Parent ion: [M + 1Me + 3H]3+ (856.73)
RT: 20.54 min
Relative intensity (%)
m/z

C

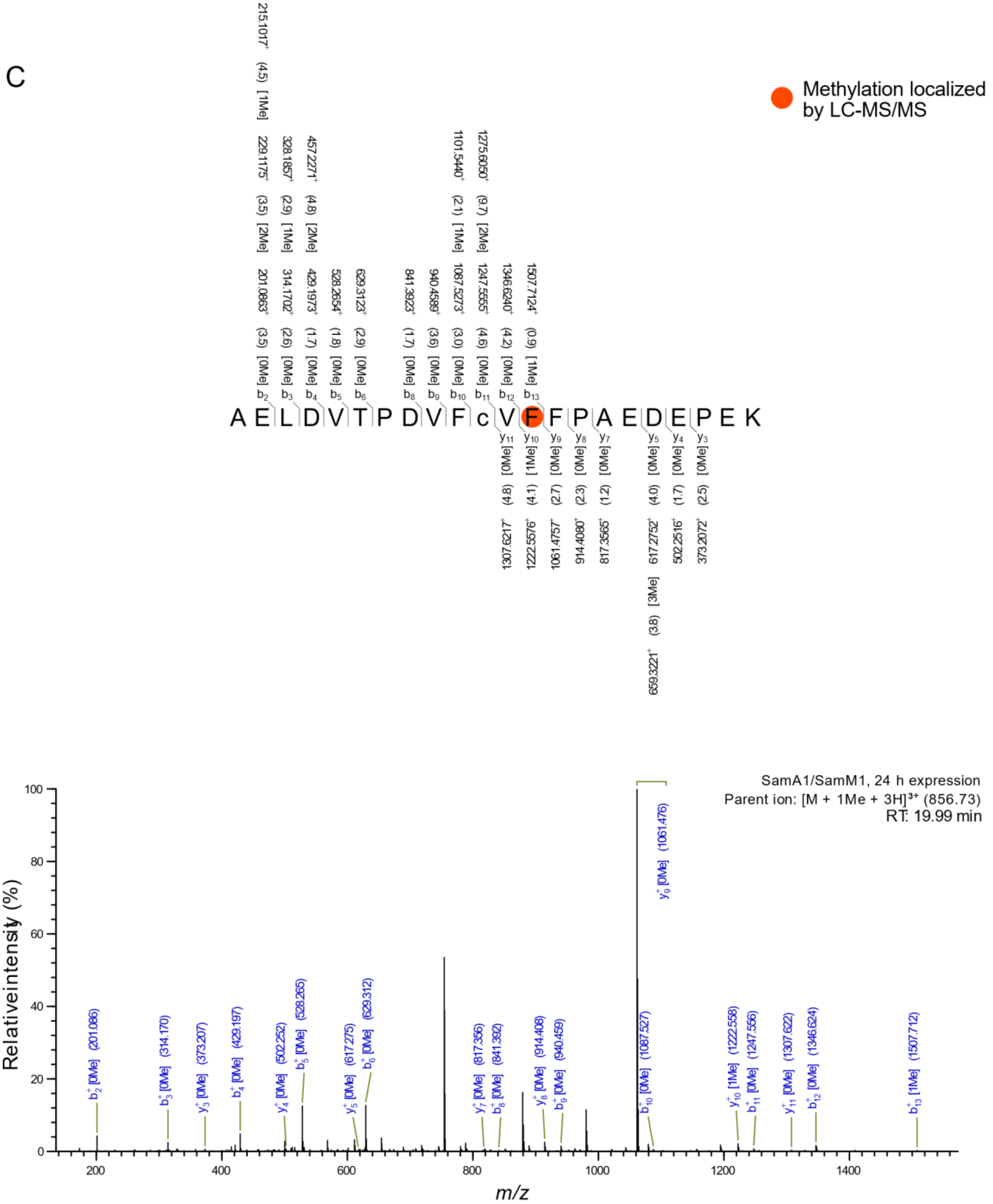
Methylation localized by LC-MS/MS
A E L D V T P D V F c V F F P A E D E P E K
SamA1/SamM1, 24 h expression
Parent ion: [M + 1Me + 3H]3+ (856.73)
RT: 19.99 min
Relative intensity (%)
m/z

D

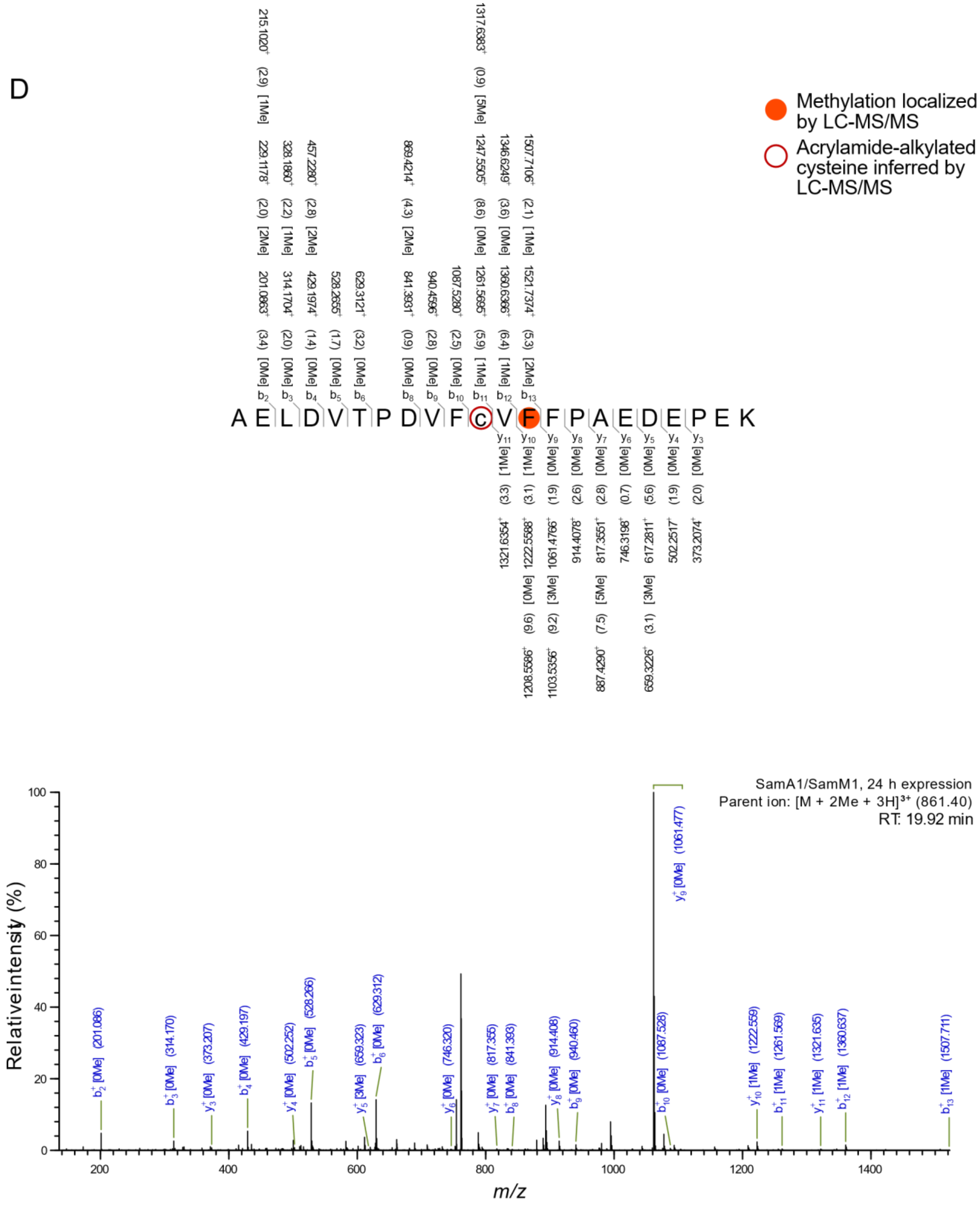
Methylation localized by LC-MS/MS
Acrylamide-alkylated cysteine inferred by LC-MS/MS
A E L D V T P D V F c V F F P A E D E P K
SamA1/SamM1, 24 h expression
Parent ion: [M + 2Me + 3H]3+ (861.40)
RT: 19.92 min
Relative intensity (%)
m/z

E

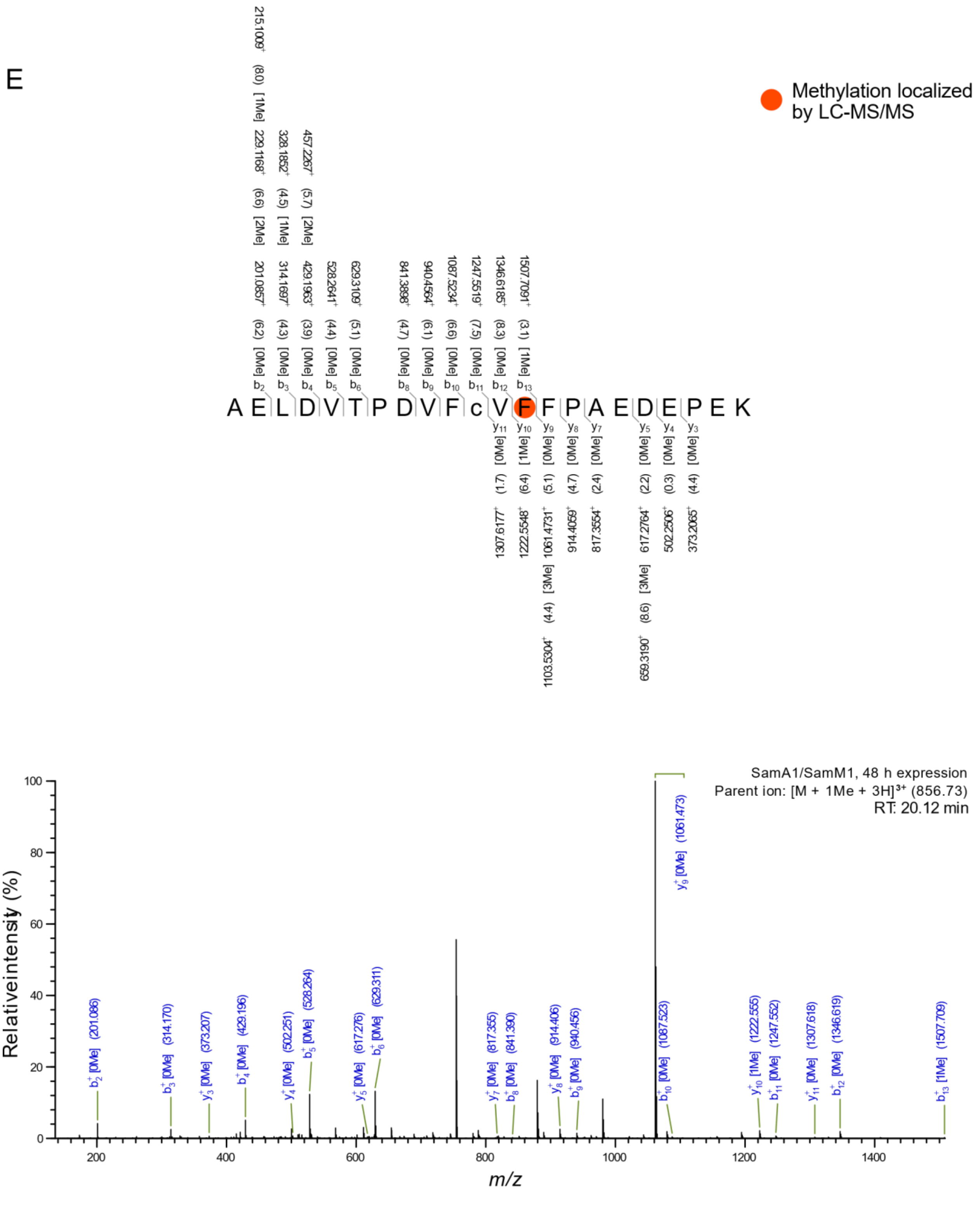

Methylation localized by LC-MS/MS
A E L D V T P D V F c V F F P A E D E P K
SamA1/SamM1, 48 h expression
Parent ion: [M + 1Me + 3H]3+ (856.73)
RT: 20.12 min
Relative intensity (%)
m/z

F

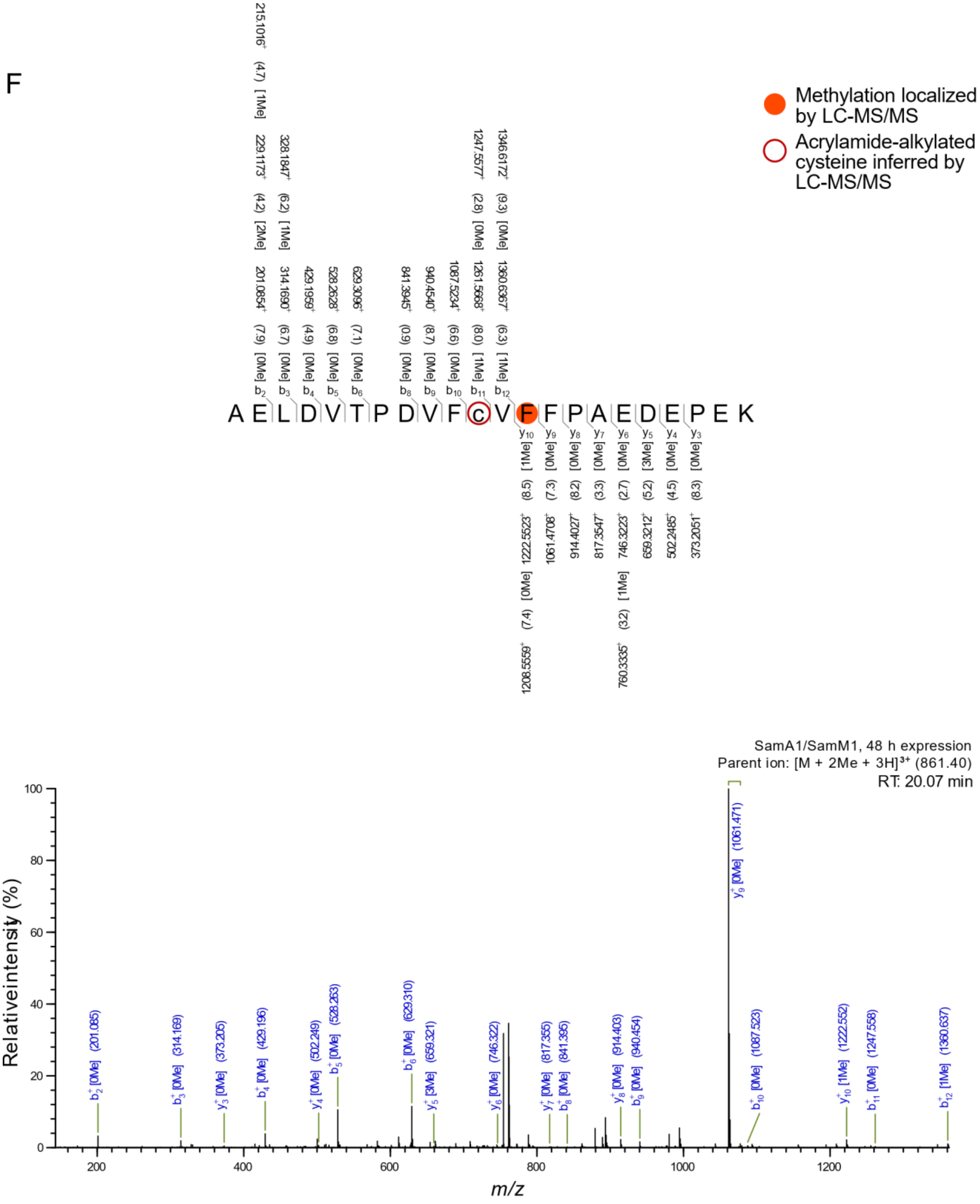
Methylation localized by LC-MS/MS
Acrylamide-alkylated cysteine inferred by LC-MS/MS
A E L D V T P D V F C V F F P A E D E P E K
SamA1/SamM1, 48 h expression
Parent ion: [M + 2Me + 3H]3+ (861.40)
RT: 20.07 min
Relative intensity (%)
m/z

G

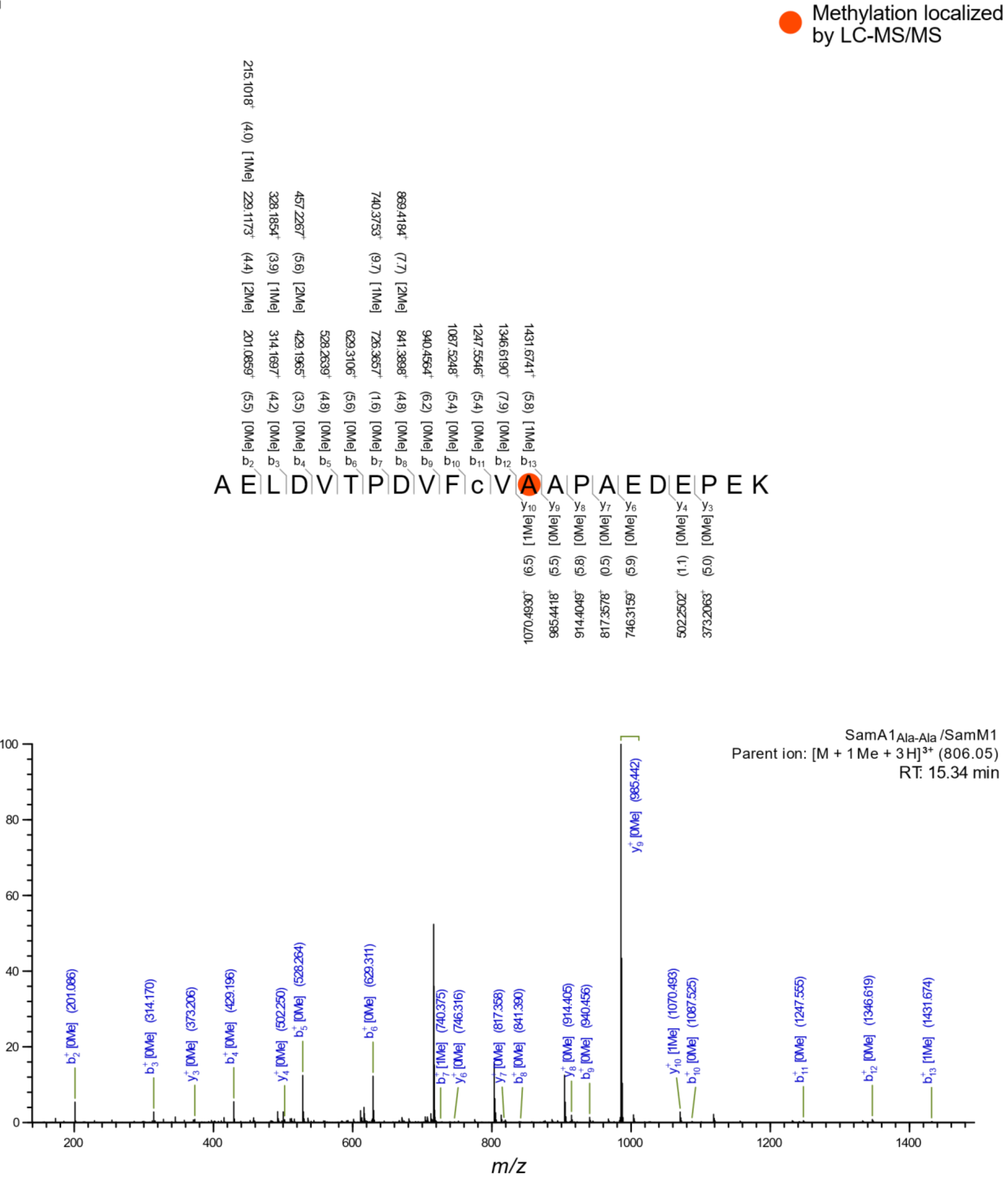

Methylation localized by LC-MS/MS
A E L D V T P D V F c V A A P A E D E P E K
SamA1Ala-Ala /SamM1
Parent ion: [M + 1Me + 3H]3+ (806.05)
RT: 15.34 min
Relative intensity (%)
m/z

H

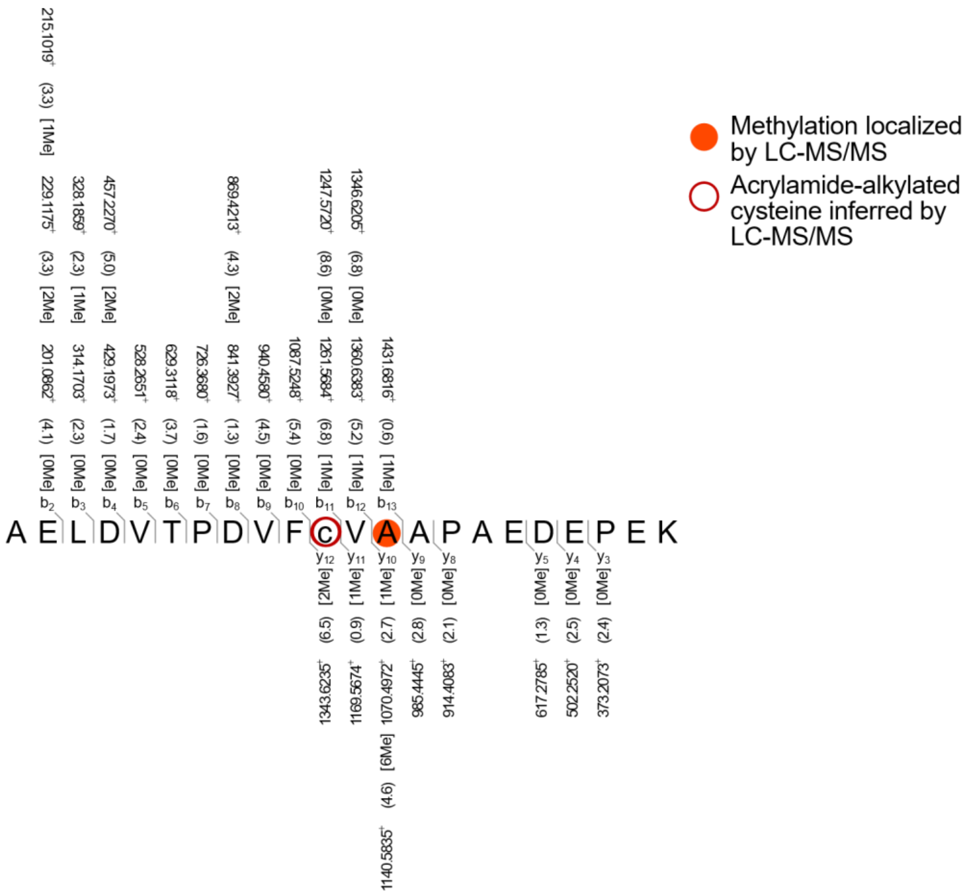
Methylation localized by LC-MS/MS
Acrylamide-alkylated cysteine inferred by LC-MS/MS
A E L D V T P D V F c V A A P A E D E P K

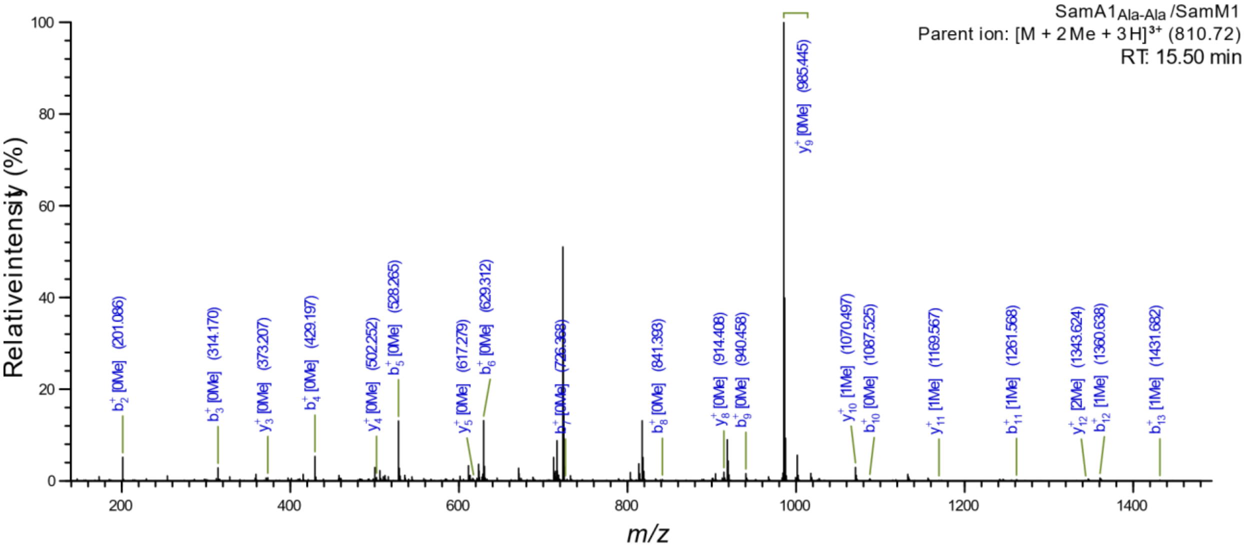
SamA1$_{Ala-Ala}$ /SamM1
Parent ion: [M + 2 Me + 3 H]$^{3+}$ (810.72)
RT: 15.50 min
Relative intensity (%)
m/z

I

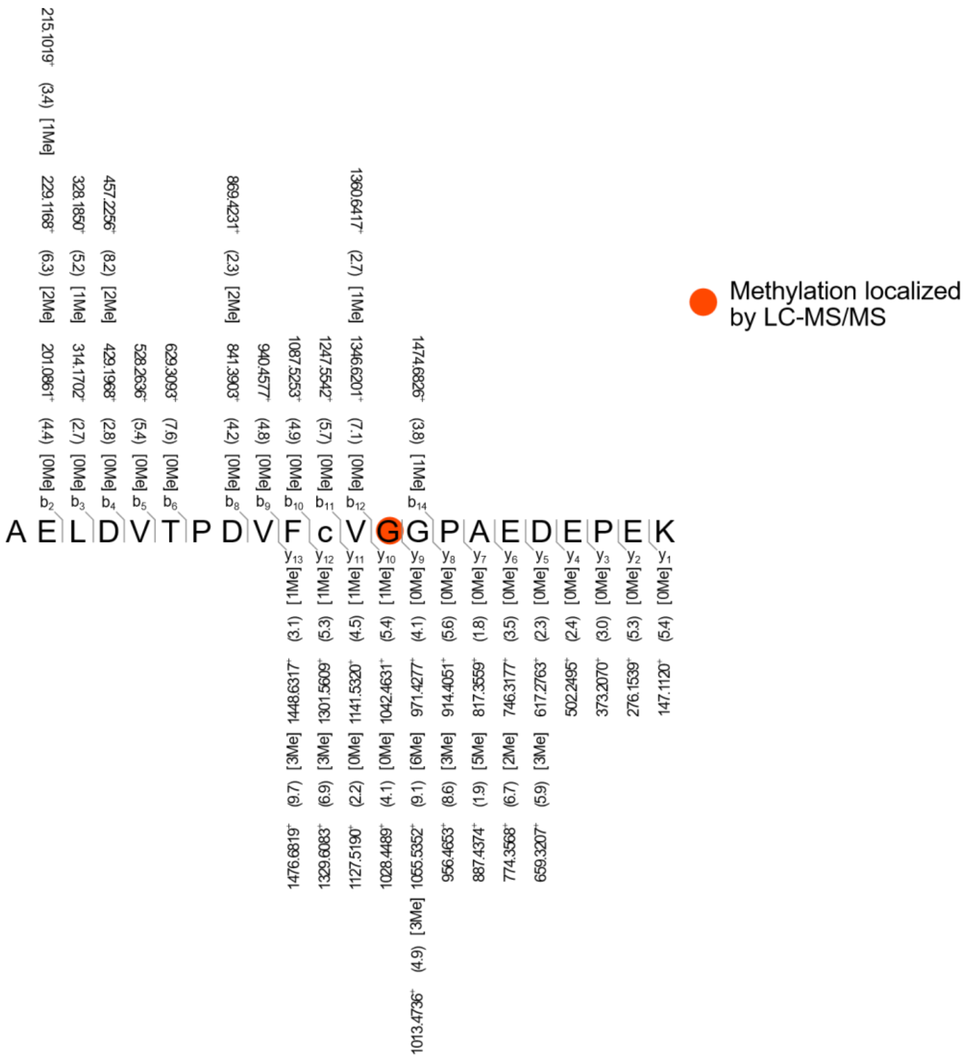
Methylation localized by LC-MS/MS
A E L D V T P D V F c V G G P A E D E P K

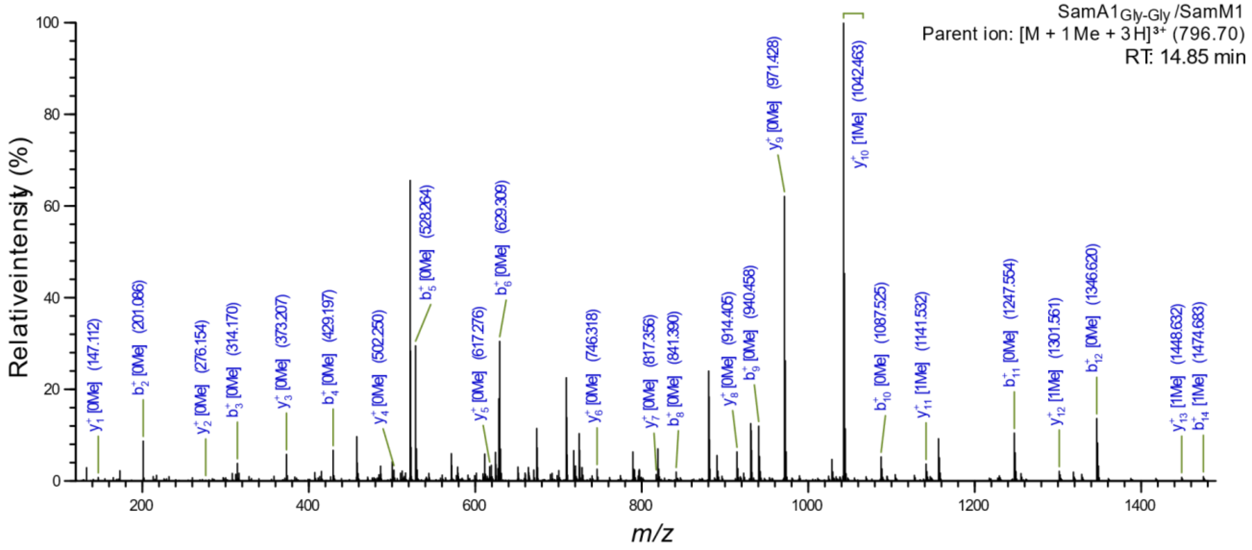
SamA1Gly-Gly /SamM1
Parent ion: [M + 1 Me + 3 H]3+ (796.70)
RT: 14.85 min
Relative intensity (%)
m/z

J

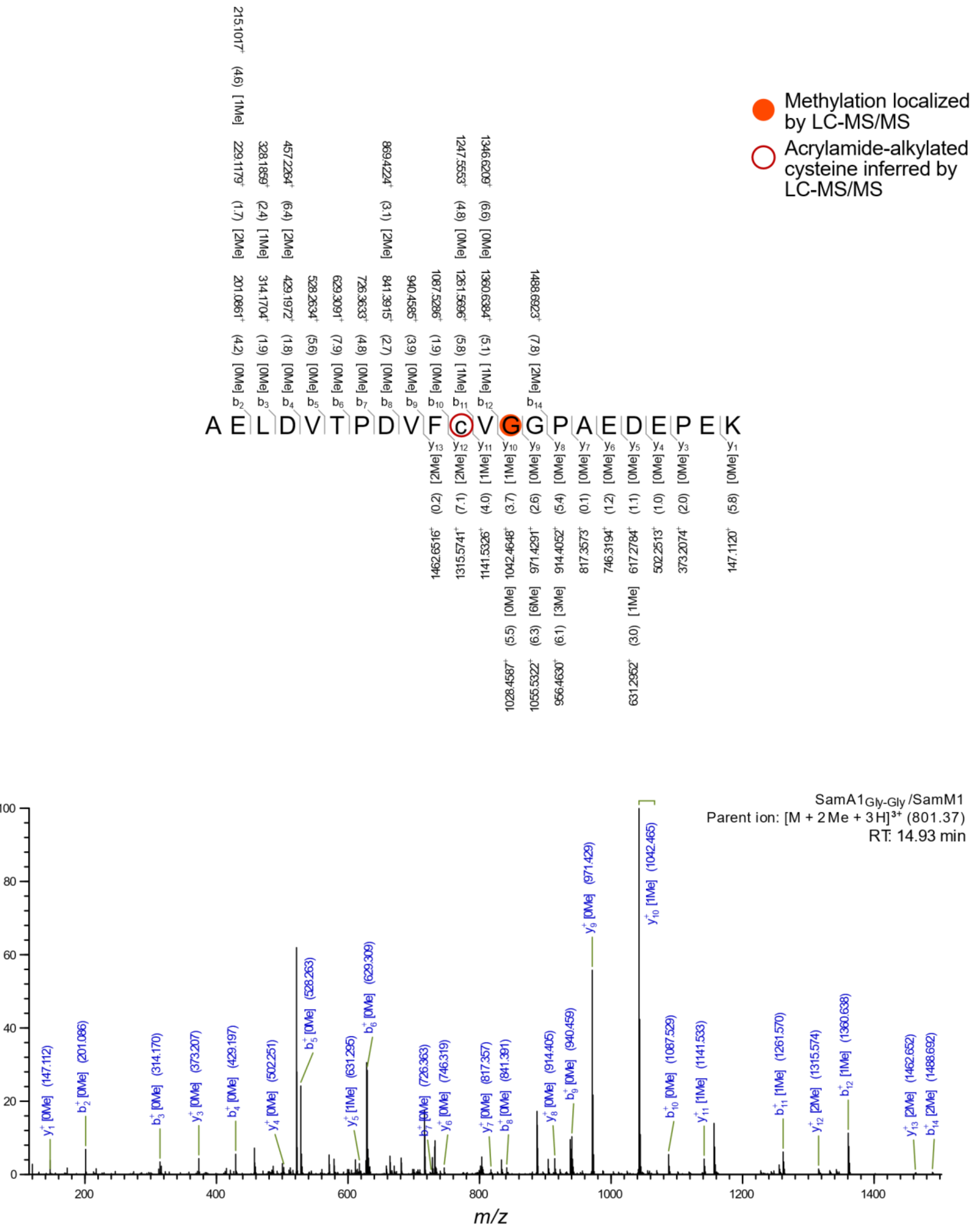


**Figure S26: MS2 fragmentation of SamA1 variants.** SamA1 and variants in-gel digested and analyzed using high-resolution LC-MS/MS. Methylation state of each peptide fragment is included in brackets, where 'Me' marks a mass shift corresponding to methylation, and methylated residues are indicated by open or filled circles. Fragments $b^+$, $y^+$, and $c^+$ ions (orange if applicable) are labeled above and below the peptide sequence with the observed m/z and the deviation from the theoretical mass in parentheses (in ppm). The lowercase ‘c’ represents the expected mass of a cysteine treated with iodoacetamide (IAA). A mass cutoff of 10.0 ppm and minimum relative peak intensity of 0.1% was used for the annotated masses. Panel **A** localizes a mass shift corresponding to methylation to Cys50 that is independent of SamM1 activity using EThcD fragmentation. Methylation localized to this position by HCD in panels **B**, **D**, **F**, **H**, and **J** are inferred as the same artifact. **A.** Singly methylated SamA1 expressed without SamM1 fragmented by EThcD. Red filled circle indicates localization of the mass shift corresponding to a methylation to a site other than on the backbone nitrogen. The observed methylation at Cys50 is likely due to alkylation by unpolymerized acrylamide during SDS-PAGE, resulting in a mass shift of 14 from the expected IAA treated mass. This modification is not seen in samples pre-treated with IAA before SDS-PAGE (data not shown). **B.** Singly methylated SamA1 expressed with SamM1 R65A Y69A (low activity variant), fragmented by HCD. Red open circle indicates the artifactual observed methylation at Cys50. **C.** Singly methylated SamA1 expressed with SamM1 for 24 h, fragmented by HCD. **D.** Doubly methylated SamA1 expressed with SamM1 for 24 h, fragmented by HCD. Red open circle indicates the artifactual observed methylation at Cys50. **E.** Singly methylated SamA1 expressed with SamM1 for 48 h, fragmented by HCD. **F.** Doubly methylated SamA1 expressed with SamM1 for 48 h, fragmented by HCD. Red open circle indicates the artifactual observed methylation at Cys50. **G.** Singly methylated SamA1$_{Ala\text{-}Ala}$ expressed with SamM1 for 24 h, fragmented by HCD. **H.** Doubly methylated SamA1$_{Gly\text{-}Gly}$ expressed with SamM1 for 24 h, fragmented by HCD. Red open circle indicates the artifactual observed methylation at Cys50. **I.** Singly methylated SamA1$_{Gly\text{-}Gly}$ expressed with SamM1 for 24 h, fragmented by HCD. **J.** Doubly methylated SamA1$_{Gly\text{-}Gly}$ expressed with SamM1 for 24 h, fragmented by HCD. Red open circle indicates the artifactual observed methylation at Cys50.

A

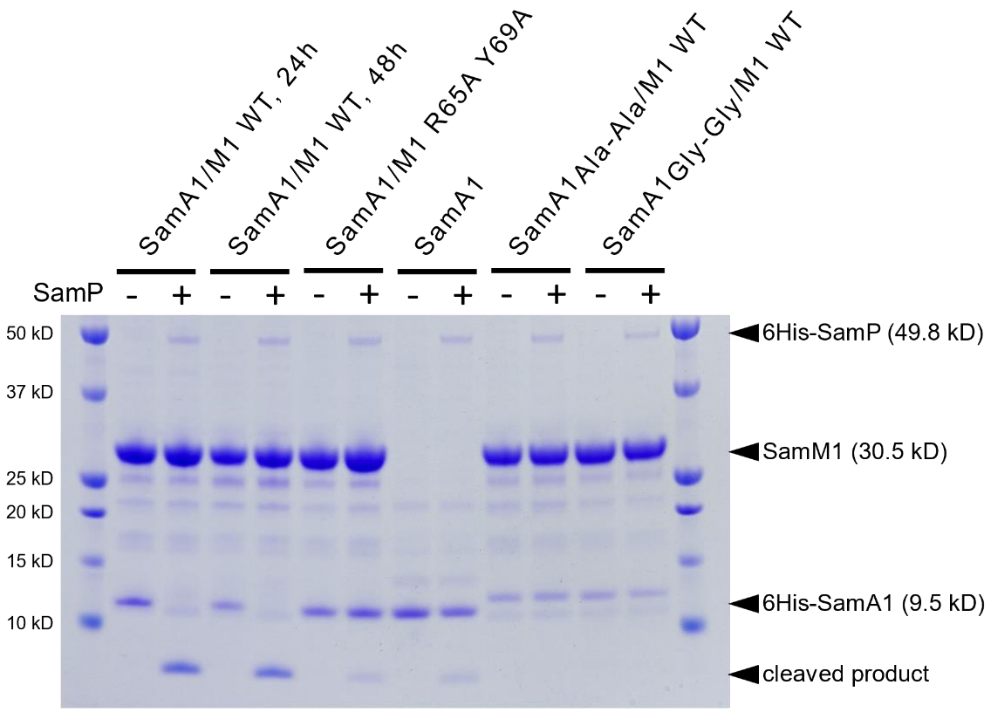
SamA1/M1 WT, 24h
SamA1/M1 WT, 48h
SamA1/M1 R65A Y69A
SamA1
SamA1 Ala-Ala/M1 WT
SamA1 Gly-Gly/M1 WT
SamP
- + - + - + - + - + - +
50 kD
37 kD
25 kD
20 kD
15 kD
10 kD
6His-SamP (49.8 kD)
SamM1 (30.5 kD)
6His-SamA1 (9.5 kD)
cleaved product

B

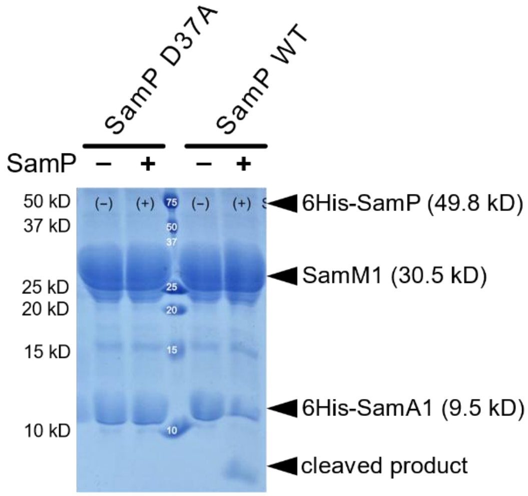
SamP D37A
SamP WT
SamP
− + − +
50 kD
37 kD
25 kD
20 kD
15 kD
10 kD
6His-SamP (49.8 kD)
SamM1 (30.5 kD)
6His-SamA1 (9.5 kD)
cleaved product

C

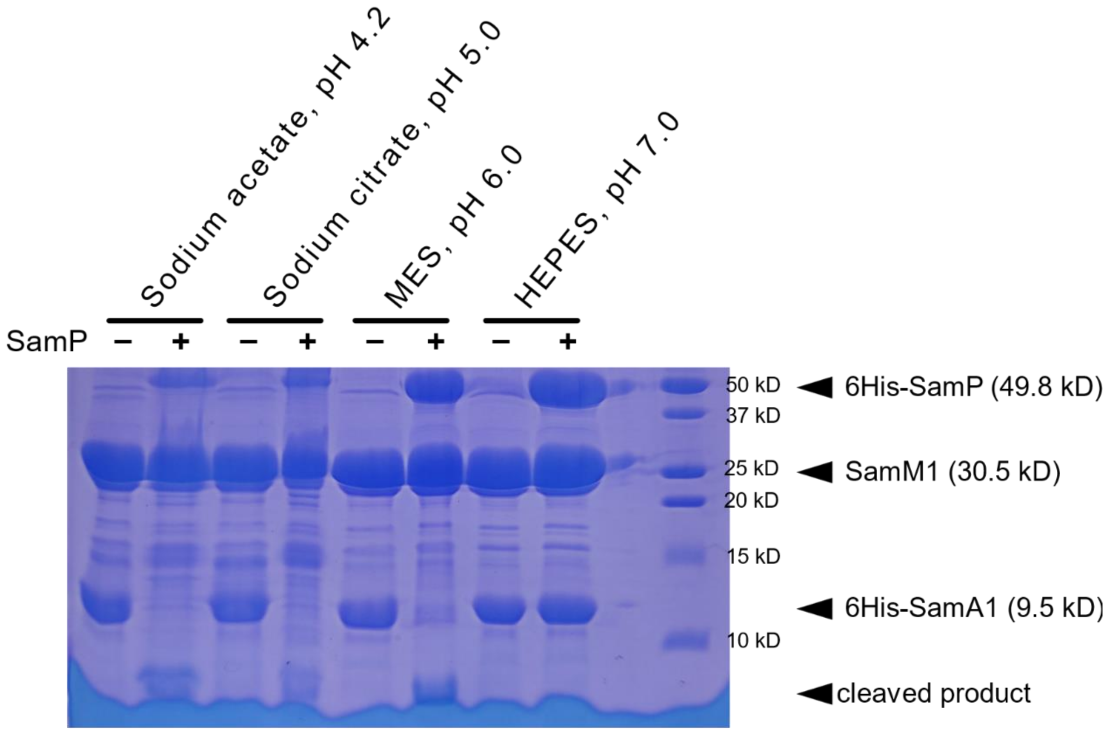
Sodium acetate, pH 4.2
Sodium citrate, pH 5.0
MES, pH 6.0
HEPES, pH 7.0
SamP – + – + – + – +
50 kD
37 kD
25 kD
20 kD
15 kD
10 kD
6His-SamP (49.8 kD)
SamM1 (30.5 kD)
6His-SamA1 (9.5 kD)
cleaved product

D

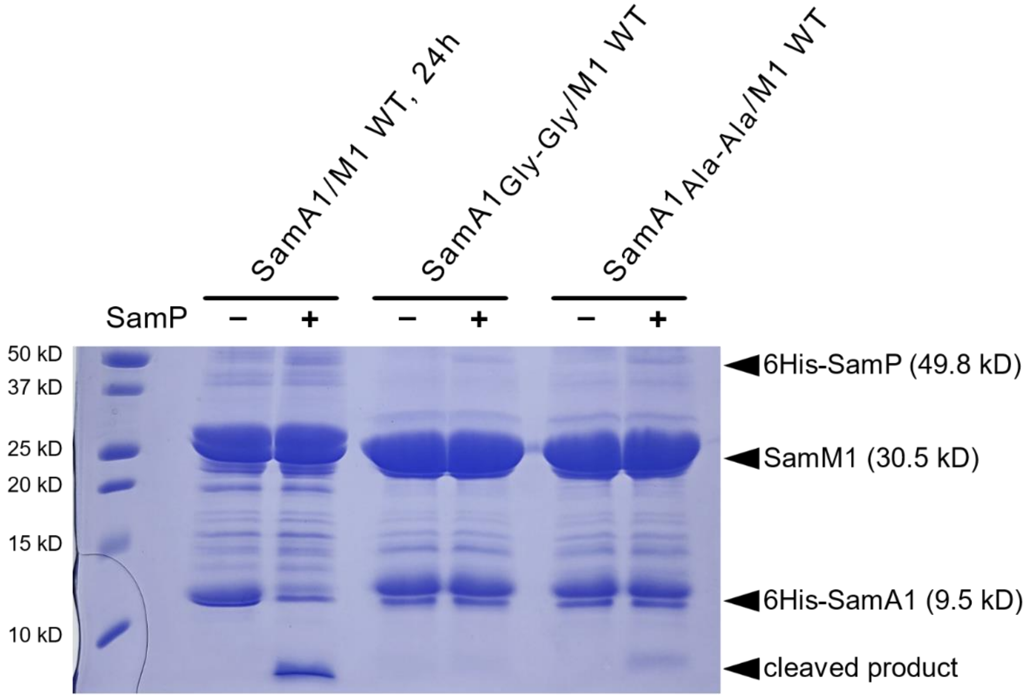
SamA1/M1 WT, 24h
SamA1 Gly-Gly/M1 WT
SamA1 Ala-Ala/M1 WT
SamP – + – + – +
50 kD
37 kD
25 kD
20 kD
15 kD
10 kD
6His-SamP (49.8 kD)
SamM1 (30.5 kD)
6His-SamA1 (9.5 kD)
cleaved product

**Figure S27: SDS-PAGE gels of SamP *in vitro* reactions.** Cleavage of His-SamA1/M1 by SamP was tested in *in vitro* reactions at 37 °C for 30-60 min. Following the reactions, tubes were centrifuged to pellet precipitant and the supernatant was run and visualized by SDS-PAGE. Negative controls indicated by "-" were performed that excluded SamP (WT or D37A) from the reaction. **A.** SamP cleavage of methylated SamA1 (SamA1/M1 24 and 48 h), un- or low-methylated SamA1 (SamA1/M1 R65A Y69A and SamA1), and SamA1 variants (SamA1$_{Ala-Ala}$/M1 and SamA1$_{Gly-Gly}$/M1) was tested *in vitro* using a 1:100 protease to substrate ratio in 30-min reactions. **B.** Cleavage of His-SamA1/M1 by WT and D37A SamP was tested *in vitro* using a 1:100 protease to substrate ratio in 60-min reactions. **C.** SamP cleavage of His-SamA1/M1 was tested *in vitro* at various pH values using a 1:10 protease to substrate ratio in 60-min reactions. **D.** SamP cleavage of SamA1$_{Ala-Ala}$ or SamA1$_{Gly-Gly}$ co-expressed with SamM1 was performed *in vitro* using a 1:100 protease to substrate ratio in 60-min reactions.

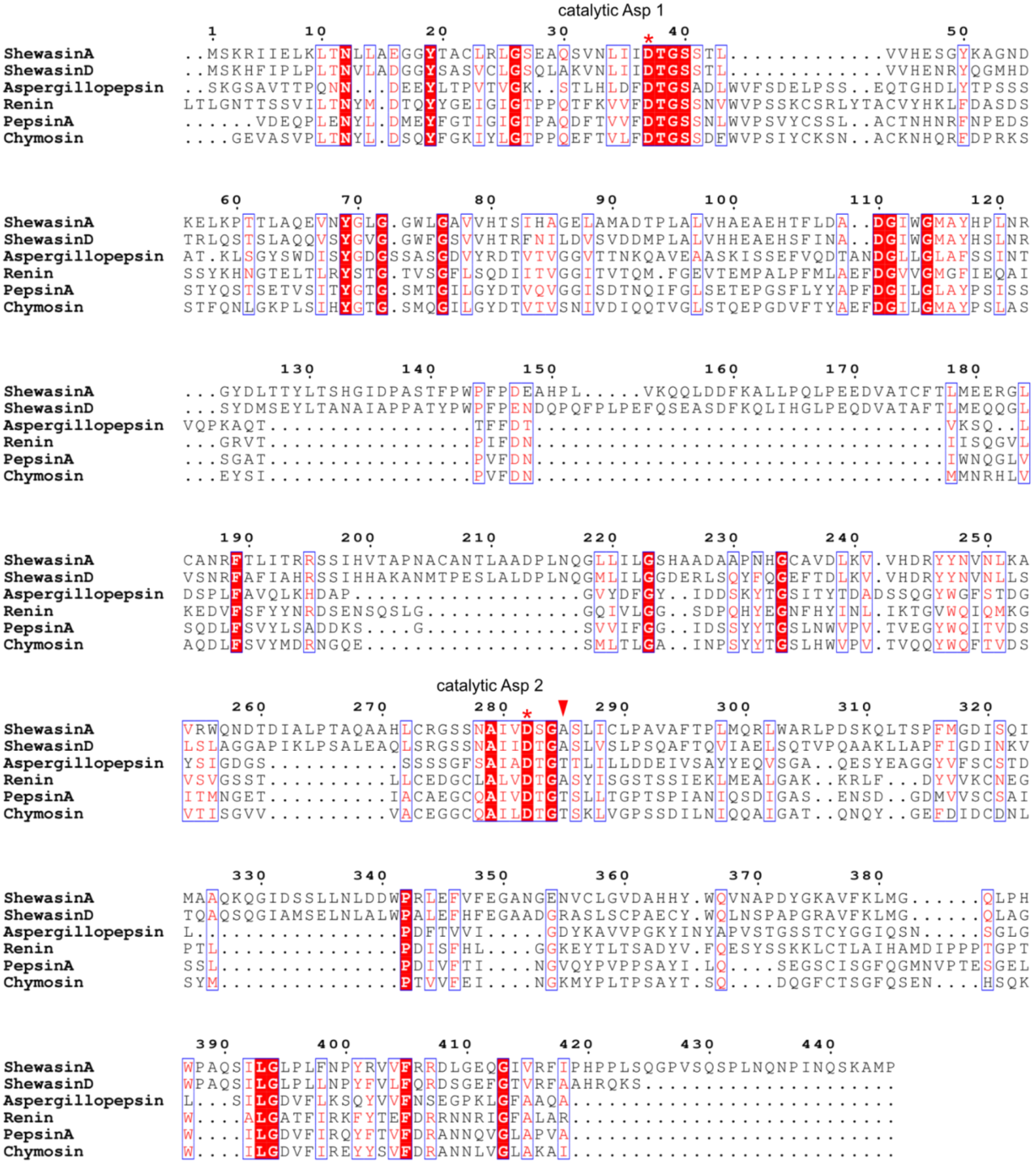


**Figure S28: Alignment of A1 peptidase family members.** MUSCLE alignment of shewasins A and D, with the eukaryotic A1 family peptidases pepsin A (*Homo sapiens*), chymosin (*Bos taurus*), aspergillopepsin (*Aspergillus phoenicis*), and renin (*Homo sapiens*). The N-terminal signaling peptides have been removed from the eukaryotic sequences. The catalytic Asp residues at positions 37 and 282 are labelled and marked with a red asterisk and the Thr → Ala substitution at position 285 is marked by a red arrow. Alignment figure was generated using ESPript 3.

A

SamP *in vitro* reactions, MS1 TICs

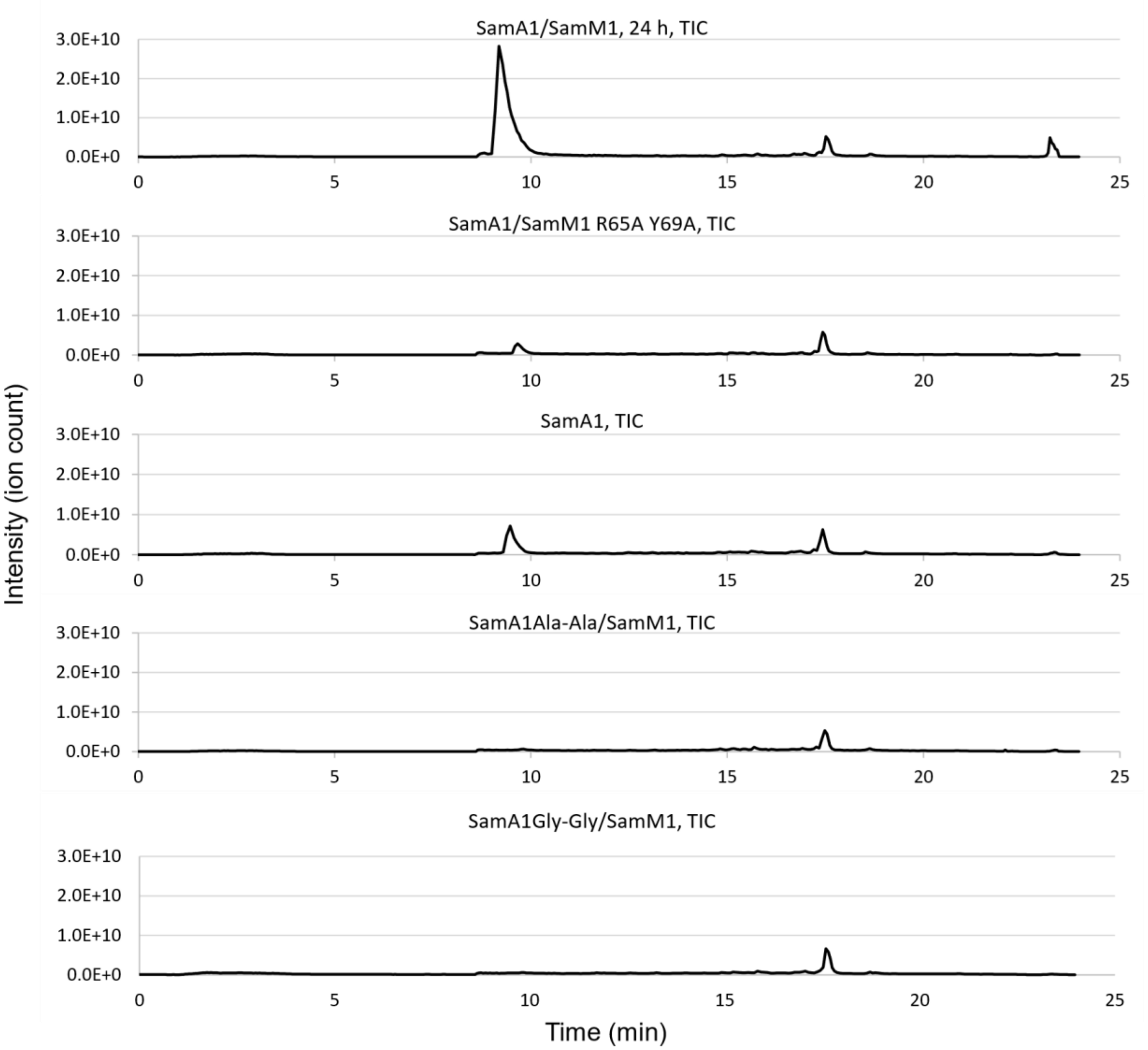


B

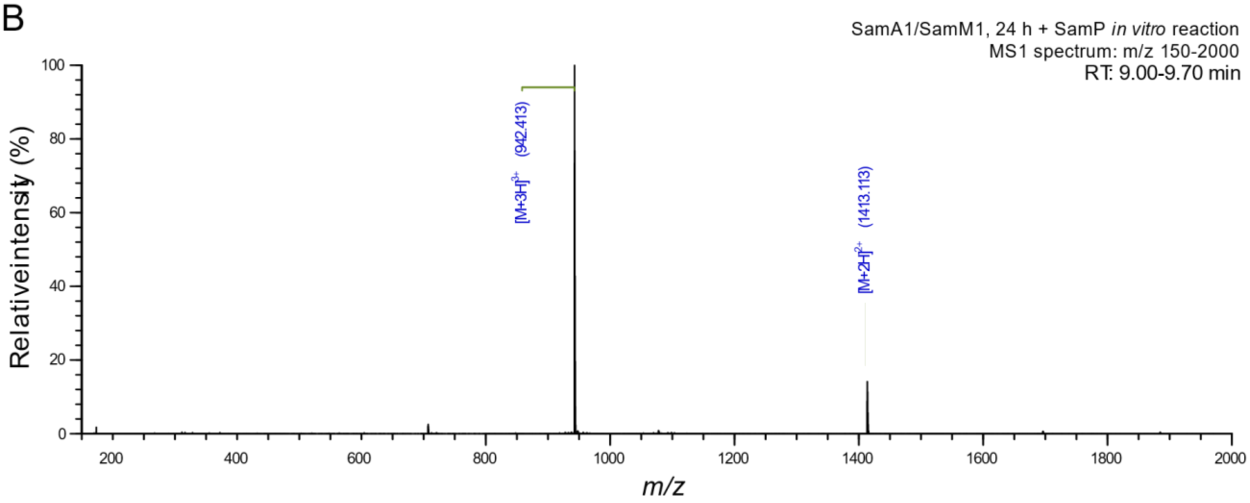

C

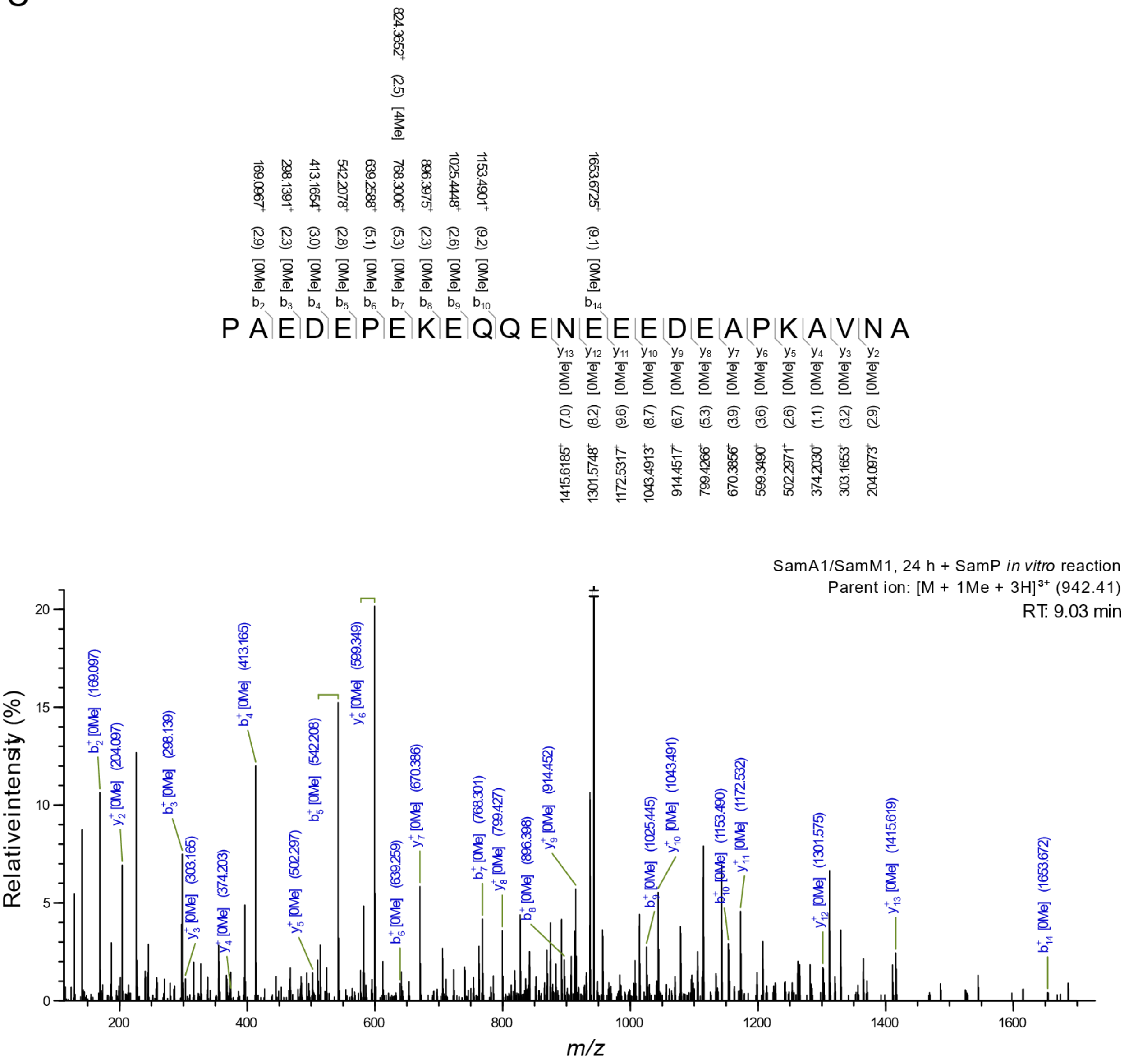


**Figure S29: LC-MS/MS analysis of SamP *in vitro* reactions.** SamP *in vitro* reactions with SamA1 and variants were analyzed by LC-MS/MS. **A.** MS1 total ion chromatograms (TICs) of each reaction with intensity in ion count shown on the y-axis. Each TIC is titled by SamA1 sample. **B.** MS1 spectrum for the dominant peak in the SamA1/M1 24 h + SamP *in vitro* reaction over a retention time range of 9.0-9.7 min and a mass range from 150-2000 m/z. The two most dominant peaks are labeled by observed mass and charge state. Both peaks correspond to a calculated neutral mass of 2824.21 - 2824.23. **C.** MS2 spectrum from a SamA1/M1, 24 h + SamP *in vitro* reaction for parent ion 942.41 at retention time 9.03 min, fragmented by HCD. Fragment $b^+$ and $y^+$ ions are labeled above and below the peptide sequence with the observed m/z and the deviation from the theoretical mass in parentheses (in ppm). A mass cutoff of 10.0 ppm and minimum relative peak intensity of 0.1% was used for the annotated masses.

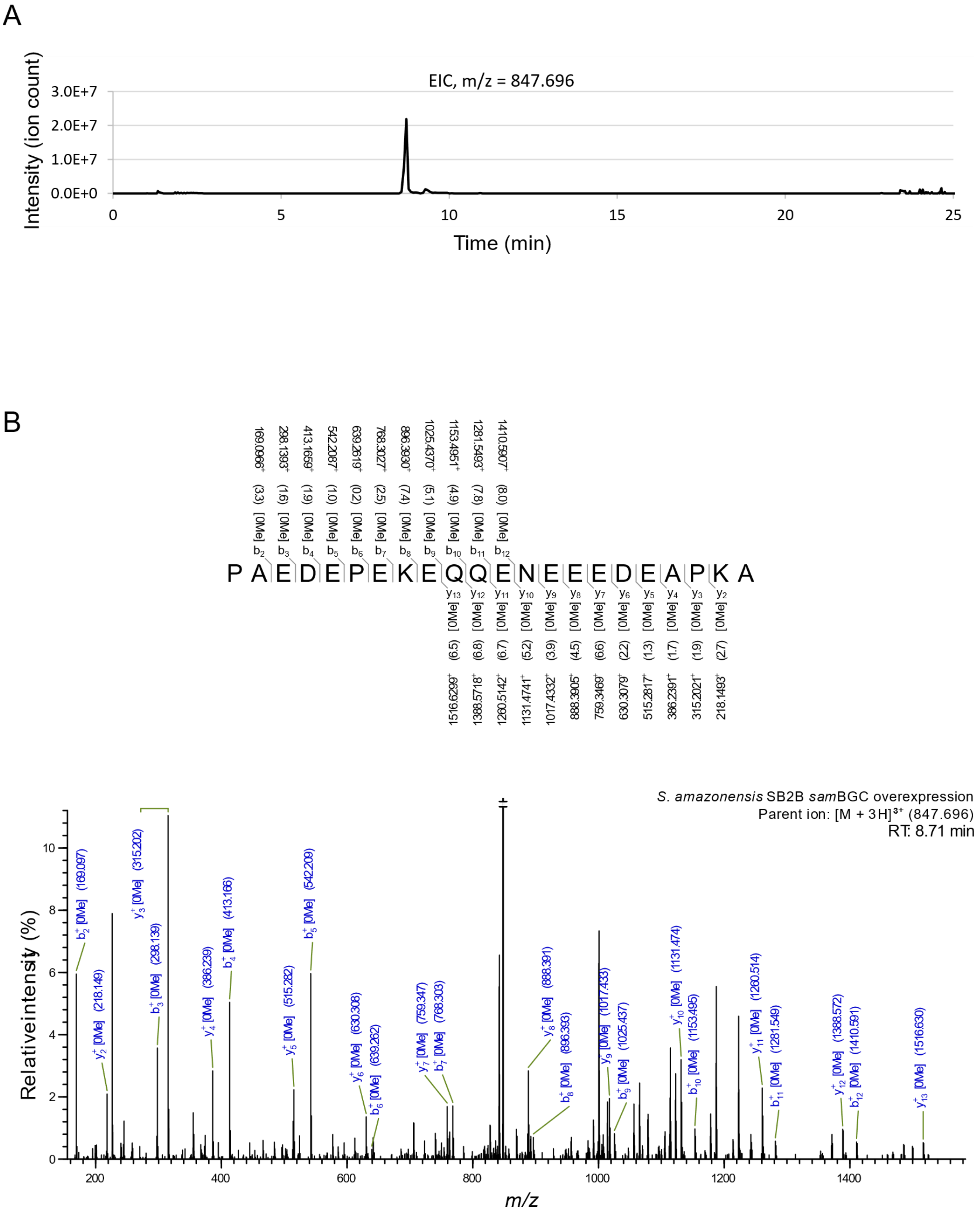


**Figure S30: LC-MS/MS analysis of metabolite extract from *sam*BGC overexpression.** High-resolution LC-MS/MS analysis of a metabolite that matches the portion of SamA1 C-terminal of the SamP cleavage site. **A.** MS1 extracted ion chromatogram of the parent ion m/z = 847.696$^{+3}$

(calculated neutral mass of 2540.064) with a mass difference range of +/- 10 amu. **B.** MS2 spectrum for parent ion 847.696, fragmented by HCD. Fragment $b^+$ and $y^+$ ions are labeled above and below the peptide sequence with the observed m/z and the deviation from the theoretical mass in parentheses (in ppm). A mass cutoff of 10.0 ppm and minimum relative peak intensity of 0.1% was used for the annotated masses.
